# The formal demography of kinship II: Multistate models, parity, and sibship

**DOI:** 10.1101/2020.03.23.003848

**Authors:** Hal Caswell

## Abstract

**Background:** Recent kinship models focus on the age structures of kin as a function of the age of the focal individual. However, variables in addition to age have important impacts. Generalizing age-specific models to multistate models including other variables is an important and hitherto unsolved problem.

**Objectives:** Our aim is to develop a multistate kinship model, classifying individuals jointly by age and other criteria (generically, “stages”).

**Methods:** We use the vec-permutation method to create multistate projection matrices including age- and stage-dependent survival, fertility, and transitions. These matrices operate on block-structured population vectors that describe the age×stage structure of each kind of kin, at each age of a focal individual.

**Results:** The new matrix formulation is directly comparable to, and greatly extends, the recent age-classified kinship model of Caswell (2019a). As an application, we derive a model that includes age and parity. We obtain, for all types of kin, the joint age×parity structure, the marginal age and parity structures, and the (normalized) parity distributions, at every age of the focal individual. We show how to use the age×parity distributions to calculate the distributions of sibship sizes of kin.

As an example, we apply the model to Slovakia (1960–2014). The results include a dramatic shift in the parity distribution as the frequency of low-parity kin increased and that of high-parity kin decreased.

**Contribution:** The new model extends the formal demographic analysis of kinship to age×stage-classified models. In addition to parity, other stage classifications, including marital status, maternal age effects, and sex are now open to analysis.

## 1 Introduction

The goal of the formal demographic analysis of kinship is to infer the kinship network implied by a set of vital rates. Goodman, Keyfitz, and Pullum (1974) presented a rigorous analysis that gave the expected numbers of kin of a focal individual of a specified age. Caswell (2019a) presented a new approach using matrix methods that gives not only the expected numbers, but also the expected age structures of any kind of kin, living or dead, at any age of the focal individual. It permits the calculation of, inter alia, the prevalence of diseases, dependency ratios, and the experience of the death of relatives.

The models of Goodman, Keyfitz, and Pullum (1974) and Caswell (2019a) both assume that age is the only factor governing the survival and fertility of kin. In those models, the age schedules of mortality and fertility are necessary and sufficient to determine the complete kinship network. They also assume that the only interesting property of the kin is age, or something that can be calculated as a function of age alone.

To the extent to which other characteristics, in addition to age, influence survival and fertility (and it is easy to think of such characteristics), the assumption of age specificity is a limitation. In this paper we generalize the Caswell (2019a) kinship model to incorporate other factors in addition to age. We refer to these factors generically as “stages” and to the resulting models as multistate or age-stage models.^1^ After presenting the general multistate kinship framework, we apply it to the case of maternal parity. Parity (the number of children born to a woman up to a given age) influences fertility, and probably mortality, and incorporating parity into the analysis provides extra information about family structures and kinship (Schoen, 2019b,a). The demographic and evolutionary implications of kinship and family structures are many and have been explored from many perspectives (e.g., Blake, 1989; Wachter, 1997; Hammel, 2005; Hrdy, 2009; Willekens, van Imhoff, and Wright, 2015; Tanskanen and Danielsbacka, 2019).

The multistate approach introduced here, when applied to parity dynamics, expands the range of kinship issues that can be explored. The parity of an individual determines the number of children that she will have to support. Increased numbers of siblings is known to be associated with lower levels of intellectual and educational achievement (Downey, 2001); the resource dilution model posits that increased sibship size requires parental resources to be spread more thinly among more children (Blake, 1989). Downey (2001) reviews a variety of evidence supporting this effect of sibship size. Kalmijn and van de Werfhorst (2016) have recently explored the gender relations of resource dilution, and find that brothers have more of an effect than sisters. The forthcoming two-sex version of the kinship model will be useful for exploring this relationship further. Sonneveldt, Plosky, and Stover (2013) report that the increased child mortality of high parity births may be partially due to reduced coverage of needed health interventions. The resource dilution hypothesis has implications across generations. Zhao and Zhang (2019) found that grandparental care of grandchildren is diluted by siblings of the parents in China, and that the availability of this childcare support influences planning by mothers for an additional child.

The effects of children on parents are diverse. Cools, Markussen, and Strøm (2017) analyze the impact of children on women’s labor force participation and career prospects. They find that increased numbers of children reduce women’s employment, work hours, and earnings. They also found strong and long-lasting negative effects on career outcomes (employment at high ranking firms) for college-educated women. On the other hand, loneliness in late life is associated with a variety of adverse health conditions and increased mortality risks (e.g., Valtorta et al., 2016). van den Broek, Tosi, and Grundy (2019) report that, in both Eastern and Western Europe, increased numbers of children and the experience of having at least one grandchild, was associated with a reduced risk of late-life loneliness.

There have been dramatic changes in sibship size patterns in many countries. Fahey (2017) analyzes the situation in the United States, and concludes that there has been a “revolution in family circumstances” since the 1970s. He documents a dramatic difference between black and white families; the sibship sizes of black children dropping even more precipitously than those of white children. Schoen (2016) analyzed parity-specific life tables for U.S. women from 2005–2010. Using an assumption of no mortality during the reproductive period, he found an increase in the proportion of parity 0 and a corresponding decrease in proportions at parity 3 or higher. We will see in Sections 6 and 7 that such changes follow naturally from the changes in mortality and fertility over the last 50 years in the particular case of Slovakia.

Kinship networks provide pathways for all kinds of intergenerational support. Parent-child and grandparent-grandchild interactions are obviously important; Song and Mare (2019) have emphasized the importance of multigenerational exposures. On a larger scale, (Kramer, 2019) argues that the human life history, characterized by earlier weaning and shorter birth intervals than is the case in other primates, results in larger kinship groups and that the resulting inter-generational support and transfer of resources among relatives has played a major role in the demographic success of humans as a species.

The analysis to be presented here, and that of Caswell (2019a), together provide a way to explore a very diverse set of properties of the kinship network, as it develops through the life of a focal individual and as it changes over time in response to changes in mortality, fertility, and stage transitions.

### 1.1 Notation and terminology

In what follows, matrices are denoted by upper case bold characters (e.g., **U**) and vectors by lower case bold characters (e.g., **a**). Vectors are column vectors by default; **x**^T^ is the transpose of **x**. The *i*th unit vector (a vector with a 1 in the *i*th location and zeros elsewhere) is **e**_*i*_. The vector **1** is a vector of ones, and the matrix **I** is the identity matrix. When necessary, subscripts are used to denote the size of a vector or matrix; e.g., **I**_*ω*_ is an identity matrix of size *ω* × *ω*.

Matrices and vectors with a tilde (e.g., **Ũ** or **ã**) are block-structured, jointly classifying individuals by age and stage. Late in the analysis, we will have occasion to use two tildes (e.g., 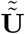). We feel apologetic about this. The symbol ° denotes the Hadamard, or element-by-element product (implemented by .* in Matlab and by * in R). The symbol ⊗ denotes the Kronecker product. The vec operator stacks the columns of a *m* × *n* matrix into a *mn* × 1 column vector. The notation ‖**x**‖ denotes the 1-norm of **x**. On occasion, Matlab notation will be used to refer to rows and columns; e.g., **F**(*i*,:) and **F**(:, *j*) refer to the *i*th row and *j*th column of the matrix **F**.

For clarity, we distinguish *structure* vectors, which give the number of individuals in each state, and *distribution* vectors, which give the proportions of individuals in each state. This is not a standard terminology, but is useful here. If the population structure is a zero vector (e.g., the age structure of the children of Focal at an age before reproduction), the distribution vector is undefined.

## 2 A brief review of the age-specific kinship model

Because the structure and derivation of the multistate kinship model follow closely the age-classified model of Caswell (2019a), we begin with a brief synopsis of that model. Kin are defined relative to a focal individual named (for purposes of reference) Focal. Focal is a member of a population in which all individuals are subject to the same age schedules of mortality and fertility. These are captured in survival and fertility matrices **U** and **F**; e.g., for the case of four age classes,

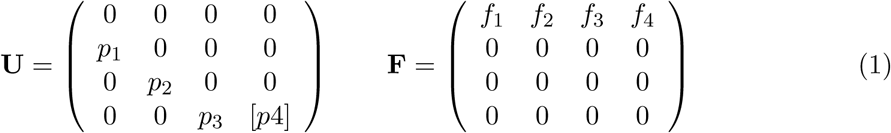

where *p*_*i*_ is the probability of survival and *f*_*i*_ the effective fertility of age class *i*. The brackets around *p*_4_ indicate that it is the survival probability of an optional open-ended age interval. It is assumed that **U** and **F** are time-invariant and have been in effect long enough that the stable age distribution can be used to calculate the age distribution of mothers. That age distribution is given by the right eigenvector **w** corresponding to the dominant eigenvalue *λ* of the projection matrix **A** = **U** + **F**, normalized to sum to one.

The key to the model of Caswell (2019a) is the recognition that the kin of any type can be treated as a population, and projected from one age of Focal to the next. The kinship network of Focal is diagrammed in Figure 1. The model describes female kin through female lines of descent. The kin of any type, at age *x* of Focal, are treated as a population described by an age structure vector denoted by the letters in Figure 1 (i.e., **a**(*x*) for daughters, **b**(*x*) for granddaughters, etc.). The symbol **k**(*x*) refers to some generic type of kin; the dynamics of this age structure are given by

**Figure 1:**
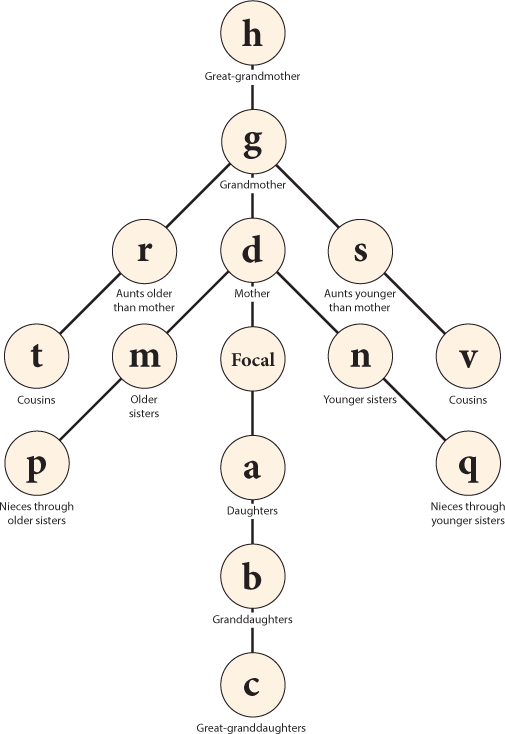
The network of kin defined in Goodman, Keyfitz, and Pullum (1974) and Keyfitz and Caswell (2005), with symbols (**ã**, 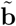, etc.) used to denote the age structure vectors of each type of kin of Focal.

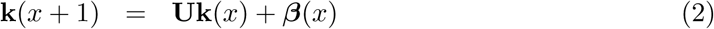

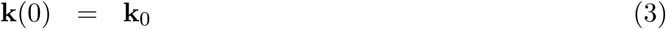

The term **Uk**(*x*) applies the survival matrix to the age structure at age *x*. The kin at age *x* of Focal survive and advance in age according to the survival schedule. The term ***β***(*x*) is a reproductive subsidy vector, which gives the production of new kin by some other type of kin (e.g., granddaughters are produced by the reproduction of daughters). The initial condition **k**_0_ specifies the age structure of the kin at the birth of Focal. For example, **a**_0_ = **b**_0_ = **c**_0_ = 0 because we may be quite sure that Focal has no daughters, granddaughters, or great-granddaughters at her birth.

It is certain that Focal has one mother alive at the time of her birth, but the age of this mother is unknown. However, the distribution of the ages of mothers in the stable population is known, and is given by

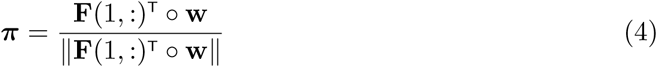

where **F**(1,:) is the first row of **F** and ‖·‖ denotes the 1-norm. At Focal’s birth, the model assumes that her mother is a randomly selected individual from this age distribution, so that

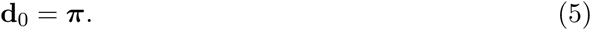

By applying this procedure methodically to all the kin in Figure 1, Caswell (2019a) derived the projection models for all kin, leading to the results given in Table 1.

**Table 1:**
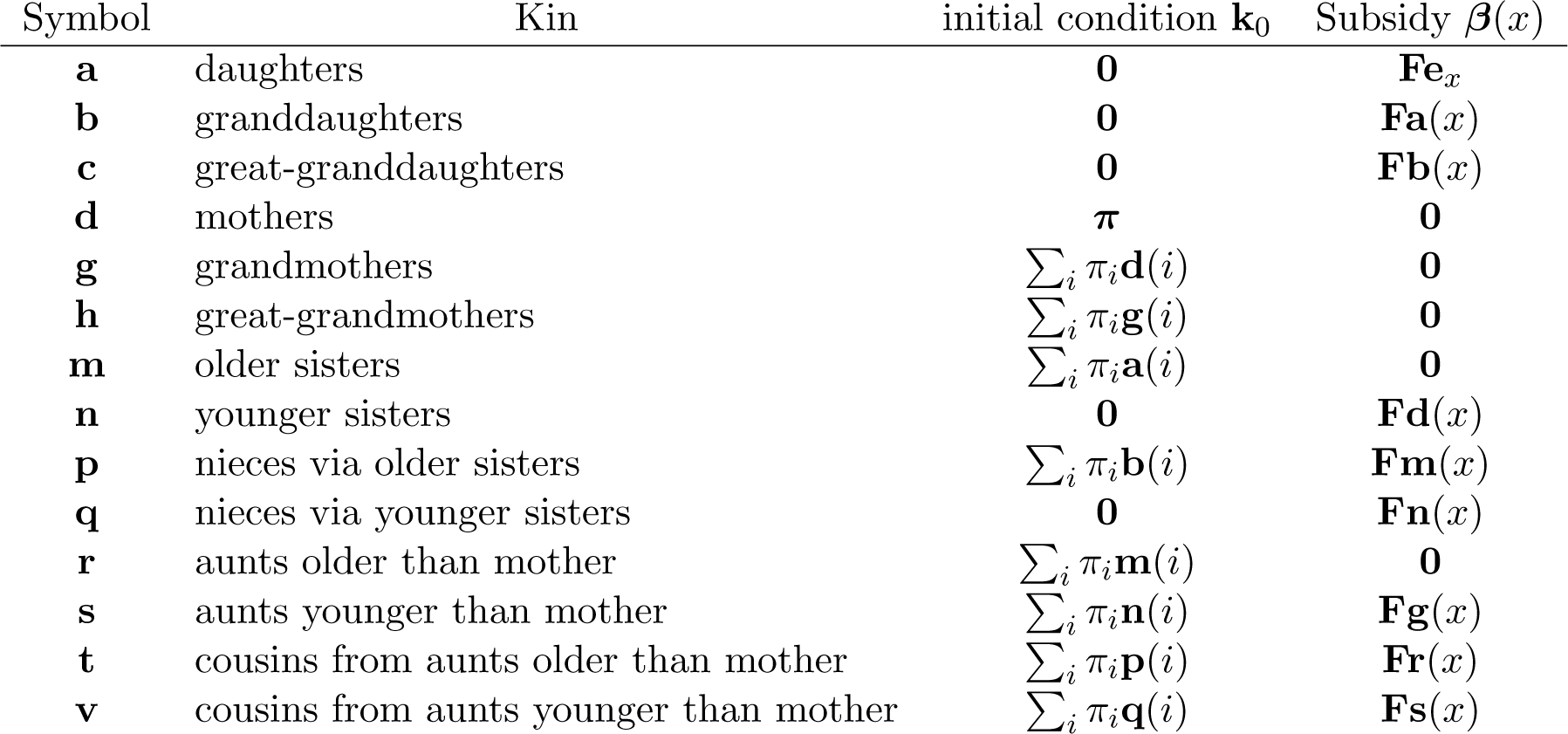
The age-classified kinship model of Caswell (2019a). Compare this with the age×stage-classified model summarized in Table 2.

| Symbol | Kin | initial condition $\mathbf{k}_0$ | Subsidy $\beta(x)$ |
| --- | --- | --- | --- |
| **a** | daughters | $\mathbf{0}$ | $\mathbf{F}\mathbf{e}_x$ |
| **b** | granddaughters | $\mathbf{0}$ | $\mathbf{F}\mathbf{a}(x)$ |
| **c** | great-granddaughters | $\mathbf{0}$ | $\mathbf{F}\mathbf{b}(x)$ |
| **d** | mothers | $\boldsymbol{\pi}$ | $\mathbf{0}$ |
| **g** | grandmothers | $\sum_i \pi_i \mathbf{d}(i)$ | $\mathbf{0}$ |
| **h** | great-grandmothers | $\sum_i \pi_i \mathbf{g}(i)$ | $\mathbf{0}$ |
| **m** | older sisters | $\sum_i \pi_i \mathbf{a}(i)$ | $\mathbf{0}$ |
| **n** | younger sisters | $\mathbf{0}$ | $\mathbf{F}\mathbf{d}(x)$ |
| **p** | nieces via older sisters | $\sum_i \pi_i \mathbf{b}(i)$ | $\mathbf{F}\mathbf{m}(x)$ |
| **q** | nieces via younger sisters | $\mathbf{0}$ | $\mathbf{F}\mathbf{n}(x)$ |
| **r** | aunts older than mother | $\sum_i \pi_i \mathbf{m}(i)$ | $\mathbf{0}$ |
| **s** | aunts younger than mother | $\sum_i \pi_i \mathbf{n}(i)$ | $\mathbf{F}\mathbf{g}(x)$ |
| **t** | cousins from aunts older than mother | $\sum_i \pi_i \mathbf{p}(i)$ | $\mathbf{F}\mathbf{r}(x)$ |
| **v** | cousins from aunts younger than mother | $\sum_i \pi_i \mathbf{q}(i)$ | $\mathbf{F}\mathbf{s}(x)$ |

## 3 Multistate kinship: the vec-permutation model

We turn now to the multistate generalization of the age-specific kin model. We develop the model carefully, because it is critical in order to the eventual result.

The multistate model classifies individuals jointly by age and some other characteristic, referred to generically as “stage.” In Section 5, the stage variable will be parity. The model is constructed using the vec-permutation matrix approach introduced by Hunter and Caswell (2005) and described in detail in Caswell et al. (2018); it has been applied, inter alia, to frailty (Caswell, 2014; Hartemink, Missov, and Caswell, 2017), latent heterogeneity (Hartemink and Caswell, 2018; Jenouvrier et al., 2018; van Daalen and Caswell, 2020), maternal age effects (Hernández et al., 2019), socioeconomic inequality (Caswell, 2019b), epidemiology (Klepac and Caswell, 2011), and genetics (de Vries and Caswell, 2019).

The details of the method, and many demographic applications, are described in Caswell et al. (2018).

Suppose the population has *ω* age classes and *s* stages, and let *k*_*ij*_ denote the number of individuals, of some type of kin, in stage *i* and age class *j*. The state of the kin population is given by the population structure vector

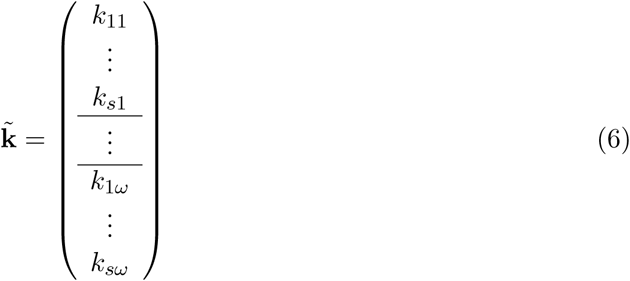

where *k*_*ij*_ is the number of kin of stage *j* in age class *i*. That is, 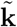 is a block-structured vector, each block of which contains the stage structure within one of the age classes.

The following sets of matrices, together, define the age- and stage-dependent demographic rates.

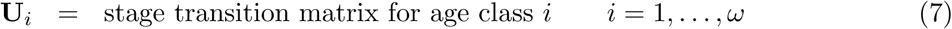

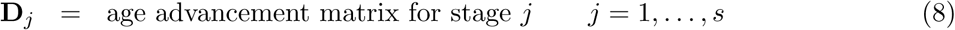

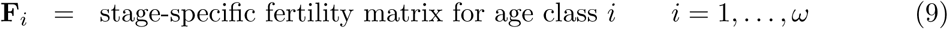

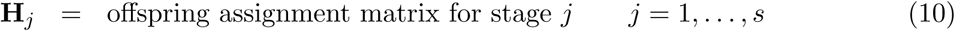

The matrices **U**_*i*_ and **F**_*i*_ are of dimension *s* × *s*. The matrices **D**_*j*_ and **H**_*j*_ are of dimension *ω* × *ω*.

The entries in **U**_*i*_ are probabilities of transitions among stages. If mortality is not included in the **U**_*i*_, the entries are transition probabilities conditional on survival, and **U**_*i*_ is column-stochastic. If mortality is included, **U**_*i*_ is column sub-stochastic.

The matrix **D**_*j*_ advances individuals of stage *j* from one age class to the next. If mortality is accounted for in the **U**_*i*_, then **D**_*j*_ is a matrix with ones on the subdiagonal and zeros elsewhere. If mortality is not included in the **U**_*i*_, **D**_*j*_ contains age-specific survival probabilities for stage *j* on the subdiagonal and zeros elsewhere.

The matrix **F**_*i*_ captures stage-specific fertility; its (*k, ℓ*) entry is the per capita production of stage *k* offspring by stage *ℓ* individuals, in age class *i*. The matrix **H**_*j*_ assigns the offspring of individuals in stage *j* to the appropriate age class. If offspring are born into the first age class, then **H**_*j*_ contains ones in the first row and zeros elsewhere.

Use these matrices to construct block diagonal matrices; e.g., from the **U**_*i*_, construct

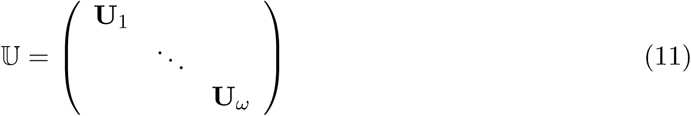

with a similar construction for 𝔻, 𝔽, and ℍ. These block-diagonal matrices are all of dimension *sω* × *sω*.

To project the joint age×stage structure of the population, construct matrices **Ũ** and 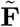 as

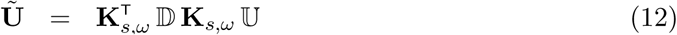

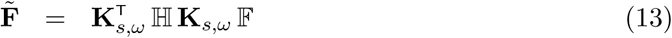

where **K**_*s,ω*_ is the vec-permutation matrix appropriate to the numbers of stages and age classes (Henderson and Searle, 1981; Caswell et al., 2018). The matrices **Ũ** and 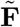 are block structured; their rows and columns inherit the same age-stage structure as 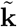 in (6). The matrix **Ũ** captures survival, transitions, and aging of extant individuals as they move among age-stage categories. The matrix 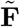 captures the production of new individuals by fertility, and the assignment of those newborn individuals into appropriate age-stage categories. The matrices **Ũ** and 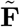 play the same role in the multistate model as the matrices **U** and **F** play in the age-classified model of Caswell (2019a).

The model simplifies in various ways depending on whether individual stage categories are fixed or dynamic. If the stages are fixed, as for example birth weight or mother’s age at birth, then the **U**_*i*_ are diagonal matrices. If stages are dynamic, as for example parity or martial status, then the structure of the **U**_*i*_ reflects the possible transitions among stages at each age.

The structure of the fertility matrices **F**_*i*_ depends on how the stage of the mother affects the stage of the offspring. If all offspring are born into the same stage (e.g., all children are born with parity 0), then **F**_*i*_ will contain non-zero entries only in the row corresponding to that stage. If offspring can be born into multiple stages, as for example when stages are defined by maternal age at birth, then the pattern of entries in the **F**_*i*_ will reflect this transmission.

## 4 Multistate kinship calculations

### 4.1 Projecting the kin populations

The population projection matrix is given by 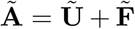. As in Caswell (2019a) we assume that Focal is a member of a population with the stable age-stage distribution implied by **Ã**. This stable distribution is given by the right eigenvector 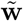 (scaled to sum to 1) of **Ã** corresponding to its largest eigenvalue.

The joint age×stage distribution of mothers in the stable population is

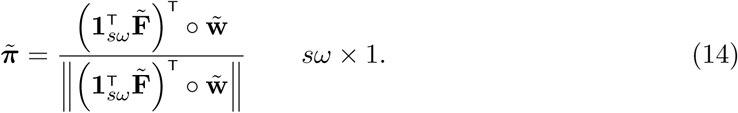

The marginal age distribution of mothers in the stable population plays an important role; it is

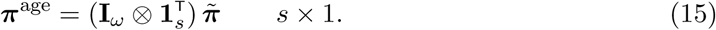

The multistate model admits the possibility that mothers may produce more than one type of offspring, which appear in different stages. These must be combined somehow in order to define the age of mothers of “offspring.” The term 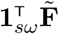 in equation (14) simply adds them up. One could instead combine them with some kind of weighted sum. I do not consider this further here.

Given the multistate model, the dynamics of the population 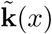 of some type of kin, jointly classified by age and stage, are given by

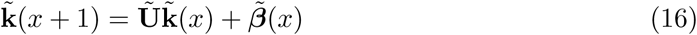

with initial condition

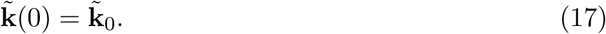

The vector 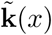 gives the joint age×stage structure of the population at age *x* of Focal. The marginal age and stage structures are calculated from 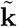 as

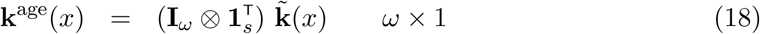

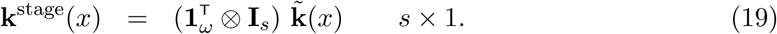

Normalizing the structure vectors **k**^age^ and **k**^stage^ so that they sum to 1 gives the proportional age and stage distributions, respectively.

This construction reduces the multistate model to the same mathematical form as the age-classified model.^2^ This makes it possible to incorporate any kind of age- and stage-specific rates into the projection of kin. Because they share the same mathematical structure, the derivations for the age×stage-classified model follow the same steps as those of the age-classified model. In an appropriate matrix-oriented language (Matlab or R) the multistate code is almost identical to the age-classified code, once the block structured matrices **Ũ** and 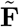 are created using (12) and (13).

### 4.2 The core kinship network

The core of the kinship network in Figure 1 is comprised of Focal, her mother, her daughters, and her older and younger sisters, as shown in Figure 2. The model for the entire network can be derived from this core (Caswell, 2019a). Schoen (2019a) presented a similar core, differing mainly in that it did not separate older and younger sisters as is done here. The following sections will derive the dynamics of each component of the core network.

**Figure 2:**
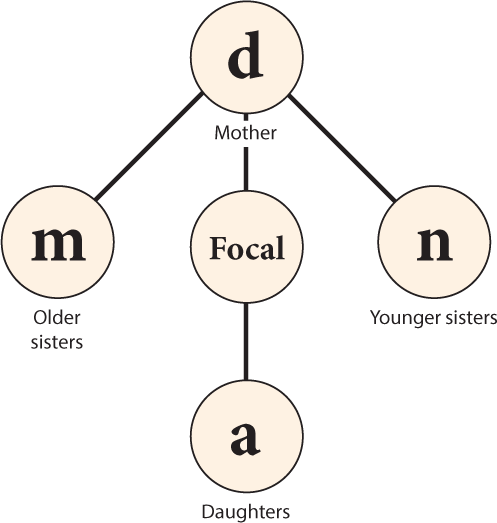
The mother-daughter-sister core of the kinship network in Figure 1

#### 4.2.1 The dynamics of Focal

In the age-classified model, the dynamics of Focal were trivial. Focal was an individual female assumed to be alive at age *x*. Thus the age distribution of Focal at age *x* was **e**_*x*_, a unit vector of length *ω* with a 1 in the *x*th place and zeros elsewhere. No further attention was needed.

In the multistate model, Focal is still an individual assumed to be alive at age *x*, but she is described by a joint age×stage distribution. This age×stage distribution will change as Focal ages and moves among stages. Thus our analysis begins with the dynamics of Focal, in essence treating her as one of her own relatives. We define

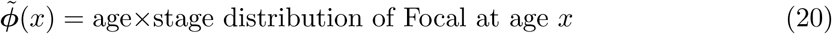

Because Focal is by definition alive, the dynamics of 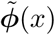 are obtained by conditioning on survival at each age up to *x*, by writing **Ũ** as

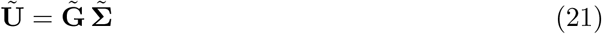

where 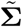 is a diagonal matrix with survival probabilities on the diagonal,

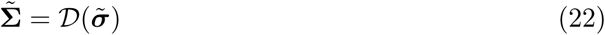

With 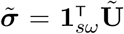 being the vector of survival probabilities, given by the column sums of **Ũ**. The matrix 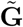 contains transition probabilities conditional on survival.

The age×stage distribution of Focal at age *x*, conditional on survival, satisfies

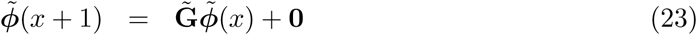

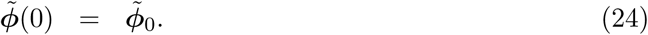

The initial condition 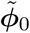 is the joint age×stage distribution of children, at birth, in the stable population,

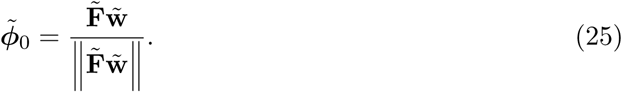

#### 4.2.2 Daughters of Focal

Daughters are the result of the reproduction of Focal. Since Focal is assumed to be alive at age *x*, the subsidy vector for daughters is 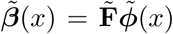, where 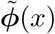 is the age×stage structure vector for Focal at age *x*. Because we may be sure that Focal has no daughters when she is born, the initial condition is **ã**_0_ = **0**. Thus

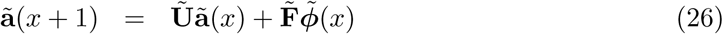

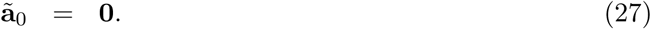

Because there is always exactly one of Focal, the age×stage structure vector is also the age×stage distribution vector.

#### 4.2.3 The mother of Focal

The population 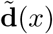 of mothers at age *x* of Focal consists of at most a single individual (step-mothers are not considered here), who is known to be alive at *x* = 0. Because no new mothers appear after Focal’s birth, the subsidy term is 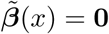.

In the age-classified model (Caswell, 2019a), the age of Focal’s mother at *x* = 0 is unknown, but has the distribution ***π***. Thus the initial population **d**_0_ of the mothers in that model was a mixture of unit vectors **e**_*i*_ (a vector of length *ω* with a one in the *i*th entry and zeros elsewhere), with mixing distribution given by ***π***. The result was an initial condition **d**_0_ = ***π***.

In the multistate model, the age and stage of Focal’s mother at *x* = 0 are unknown, but their joint distribution is given by 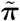, with the marginal age distribution ***π***^age^. The stage distribution of Focal’s mother is a mixture of the conditional stage distributions (conditional on mother’s age), with mixing distribution ***π***^age^. Define the conditional age×stage distribution of mothers, conditional on a specified age, as

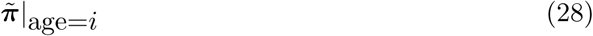

This conditional distribution is given by

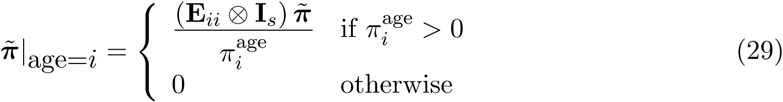

Then it follows that

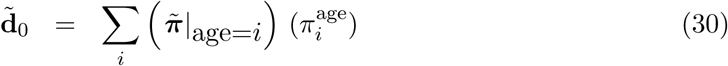

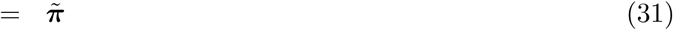

In other words, just as the initial condition for mothers in the age-classified model was given by the age distribution ***π*** of mothers in the stable population, in the age×stage-classified model, it is given by the age×stage distribution 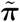 of mothers in the stable population.

The initial condition must sometimes be adjusted to satisfy additional constraints beyond the requirement that Focal’s mother be alive at her birth. For example, if stages represent parity classes, the mother of Focal cannot be in parity state 0. This requires modification of the distribution 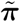, but leads to the same eventual calculations (see Section 5).

#### 4.2.4 Older sisters of Focal

The age×stage structure vector of the older sisters of Focal is denoted by 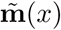. Once Focal is born, she can accumulate no more older sisters, so the subsidy term is 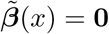. At Focal’s birth, her older sisters are the children **ã**(*i*) of the mother of Focal at the age *i* of Focal’s mother at Focal’s birth. This age is unknown, so the initial condition 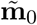 is a mixture of the age structures of children with a mixing distribution given by the marginal age distribution of mothers ***π***^age^:

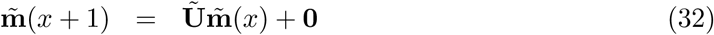

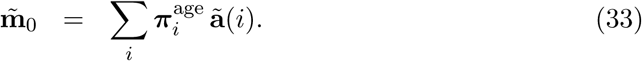

#### 4.2.5 Younger sisters of Focal

Focal has no younger sisters (**ñ**) when she is born, so the initial condition is **n**_0_ = **0**. Younger sisters are the children of Focal’s mother, so the subsidy term is the reproduction of Focal’s mother at age *x* of Focal. Thus

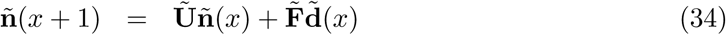

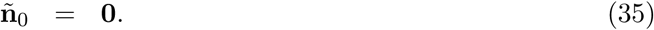

### 4.3 From the core kin to the rest of the kinship network

The age×stage structures of daughters, mothers, and sisters of Focal follow the same dynamics as the corresponding age structures in the age-classified model, only replacing the matrices **U** and **F** with **Ũ** and 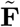 and modifying the initial conditions appropriately. The extension of this core to the entire kinship network (granddaughters, nieces, cousins, etc.) follows closely the derivations presented in Caswell (2019a). The results are shown in Table 2; comparison with Table 1 shows how similar the results are, once the age×stage model is formulated using the vec permutation matrix. For the curious, the complete set of derivations is given in Appendix Appendix A. As in the age-classified model, each kin type depends only on kin types above it in the table. Thus there are no circular dependencies to render the model insoluble. Note also that the side chains through nieces, cousins, etc. can be extended (to grand nieces, second cousins, etc.) just as the chains of descendants and ancestors are extended in Caswell (2019a).

**Table 2:** Summary of the components of the age×stage-classified kinship model. Matrices and vectors bearing tildes (e.g., **ã**) inherit the age×stage structure given by equation (6). Compare with the age-classified summary in Table 1. The vector ***π***^age^ is the marginal distribution of the ages of mothers in the stable population.

| Symbol | Kin | initial condition $\mathbf{k}_0$ | Subsidy $\beta(x)$ |
| --- | --- | --- | --- |
| $\tilde{\phi}$ | Focal | $\tilde{\phi}_0$ | $\mathbf{0}$ |
| $\tilde{\mathbf{a}}$ | daughters | $\mathbf{0}$ | $\tilde{\mathbf{F}}\tilde{\phi}(x)$ |
| $\tilde{\mathbf{b}}$ | granddaughters | $\mathbf{0}$ | $\tilde{\mathbf{F}}\tilde{\mathbf{a}}(x)$ |
| $\tilde{\mathbf{c}}$ | great-granddaughters | $\mathbf{0}$ | $\tilde{\mathbf{F}}\tilde{\mathbf{b}}(x)$ |
| $\tilde{\mathbf{d}}$ | mothers | $\tilde{\pi}$ | $\mathbf{0}$ |
| $\tilde{\mathbf{g}}$ | grandmothers | $\sum_i \pi_i^{\text{age}} \tilde{\mathbf{d}}(i)$ | $\mathbf{0}$ |
| $\tilde{\mathbf{h}}$ | great-grandmothers | $\sum_i \pi_i^{\text{age}} \tilde{\mathbf{g}}(i)$ | $\mathbf{0}$ |
| $\tilde{\mathbf{m}}$ | older sisters | $\sum_i \pi_i^{\text{age}} \tilde{\mathbf{a}}(i)$ | $\mathbf{0}$ |
| $\tilde{\mathbf{n}}$ | younger sisters | $\mathbf{0}$ | $\tilde{\mathbf{F}}\tilde{\mathbf{d}}(i)$ |
| $\tilde{\mathbf{p}}$ | nieces via older sisters | $\sum_i \pi_i^{\text{age}} \tilde{\mathbf{b}}(i)$ | $\tilde{\mathbf{F}}\tilde{\mathbf{m}}(x)$ |
| $\tilde{\mathbf{q}}$ | nieces via younger sisters | $\mathbf{0}$ | $\tilde{\mathbf{F}}\tilde{\mathbf{n}}(i)$ |
| $\tilde{\mathbf{r}}$ | aunts older than mother | $\sum_i \pi_i^{\text{age}} \tilde{\mathbf{m}}(x)$ | $\mathbf{0}$ |
| $\tilde{\mathbf{s}}$ | aunts younger than mother | $\sum_i \pi_i^{\text{age}} \tilde{\mathbf{n}}(i)$ | $\tilde{\mathbf{F}}\tilde{\mathbf{g}}(x)$ |
| $\tilde{\mathbf{t}}$ | cousins: aunts older than mother | $\sum_i \pi_i^{\text{age}} \tilde{\mathbf{p}}(i)$ | $\tilde{\mathbf{F}}\tilde{\mathbf{r}}(x)$ |
| $\tilde{\mathbf{v}}$ | cousins: aunts younger than mother | $\sum_i \pi_i^{\text{age}} \tilde{\mathbf{q}}(i)$ | $\tilde{\mathbf{F}}\tilde{\mathbf{s}}(x)$ |

## 5 Multistate kinship: Age and parity

Parity, the number of live births that a female has had, is a particularly interesting stage variable because of its close connection to fertility. Familiar age-specific fertility rates are an aggregation of age×parity-specific rates, Ignoring the parity dimension can be criticized fo failing “to capture the underlying variance of parity heterogeneity and its implications for kinship” (Schoen, 2019b). Parity dynamics change over time and are known to differ among populations and among social groups within populations (e.g., Preston, 1976).

Individuals move through parity stages, from parity 0 to parity 1 at the first birth, from parity 1 to parity 2 at the second birth, and so on. (We neglect multiple births.) The probabilities of these transitions depend on age and appear in the matrices **U**_*i*_.

Parity also can affect maternal mortality, although the details are unclear. Barclay and Kolk (2019) found a U-shaped relationship between maternal mortality and parity, with a minimum at parity 2. There is some evidence that higher parity is associated with increased risk of certain cancers (Plesko et al., 1985; Kravdal, Glattre, and Haldorsen, 1991). Higher parity is also associated with increased infant and child mortality (Sonneveldt, Plosky, and Stover, 2013). In the absence of sufficiently detailed data, we do not consider the effects of parity on mortality here. However, if such data were available, mortality effects would appear in the multistate model as part of the age advancement matrices **D**_*j*_ in equation (12).

### 5.1 Age and parity: matrix construction

In an age×parity model, the matrices **U**_*i*_ in equation (11) describe transitions among parity stages, conditional on survival, for a woman of age class *i*. Suppose (as is the case for data in the Human Fertility Database) that six parity classes are recognized (0,1,2,3,4,5+). The parity transition matrix for age class *i* is then

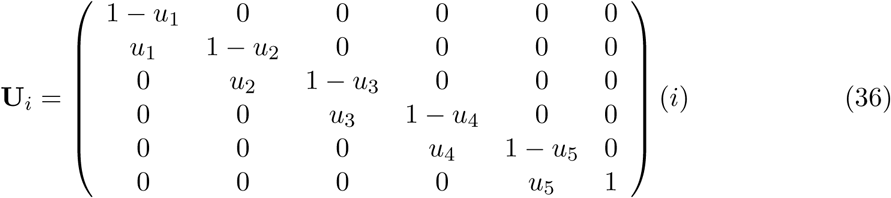

where *u*_*k*_(*x*) is the probability of a *k*th birth, over the next time interval, to a woman of age *i* and parity *k* − 1. The parity class 5+ contains women of parity 5 and all higher parities. In the Human Fertility Database, the transition probabilities are obtained (Jasilioniene et al., 2019, p. 51) from conditional parity-specific birth rates *m*_*k*_ as

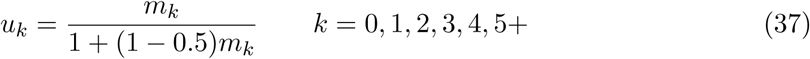

Because the transition probabilities in the **U**_*i*_ are conditional on survival, the age advancement matrices **D**_*j*_ contain age-specific survival probabilities for women of each parity class. Suppose, for example, that *ω* = 4; then

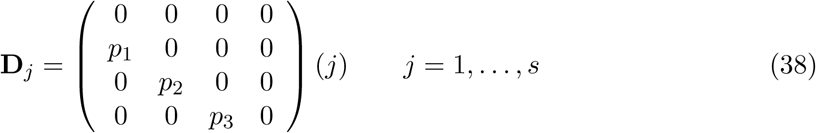

where *p*_*k*_ = 1 − *q*_*k*_ is the probability of survival of an individual in age class *k* and parity stage *j*. In the example we will consider below, all the **D**_*j*_ are equal because we lack parity-specific mortality schedules, but we give the general formulation here for future reference. An analysis including parity-specific mortality would be useful for exploring the evolutionary demography of parity progression.

The matrix **F**_*i*_ contains stage-specific fertility for females in age class *i*. In an age×parity model, reproduction is associated with transitions among the parity classes. The probabilities of those transitions are included in the **U** matrices, so that

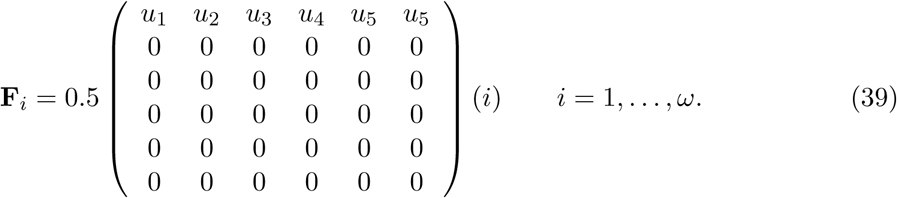

Entries appear only in the first row because all offspring are born into parity stage 0. This kinship model includes only female kin, so the factor 0.5 assumes an even sex ratio at birth. The Human Fertility Database does not differentiate among individuals within the 5+ parity stage, so they are assigned the birth probability (*u*_5_) of the highest stage recorded. Because all offspring are born into the first age class, the age assignment matrices **H**_*j*_ are the same for all parity classes; e.g., for *ω* = 4

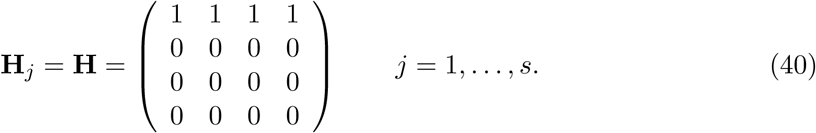

### 5.2 Age and parity: initial conditions

The age×parity model requires some modification of the initial condition 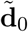 for the mother of Focal. Focal must have exactly one mother, she must be alive, and being a mother, she may not be in parity class 0. Thus the initial vector 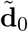 is a mixture, over the marginal distribution of mother’s ages, of age×parity vectors 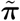 with parity 0 removed and then normalized to sum to 1. Using this modified vector in place of 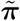, the calculation of 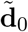 follows the steps in Section 4.2.3. Because backwards parity transitions are impossible, these conditions on 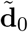 also guarantee that 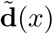 at later ages will not contain parity 0.

Similarly, Focal’s grandmother and great-grandmother (and further generations if included) may not be in parity class 0. Because the initial condition for grandmothers is a mixture of vectors for the mothers,

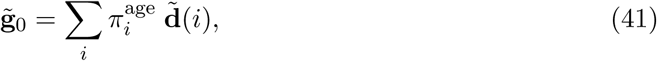

the constraint will also be satisfied for grandmothers and great-grandmothers.

## 6 A case study: age and parity in Slovakia

As an example we apply the age×parity analysis to Slovakia, using data from the Human Fertility and Human Mortality databases. The Human Fertility Database (Human Fertility Database, 2019) contains data on age- and age×parity-specific fertility for, as of this writing, thirty-one countries. We selected the series for Slovakia as a case study because, of all the countries currently in the database, Slovakia has one of the longest time series of data (1950– 2014)^3^ and one of the steepest declines in total fertility rates, from TFR = 3.6 in 1950 to TFR = 1.5 in 2014. Over the same period, life expectancy at birth increased from 62.5 to 80.3 years. The mean age at birth changed little, from 28.6 to 29.1 years.

The HFD file of conditional age-specific fertility rates contains, for each age, the probability of a first, second, third, fourth, and fifth or higher birth. These are the probabilities of transition from parity 0 to 1, from 1 to 2, from 2 to 3, from 3 to 4, and from 4 to 5 or more, respectively. These probabilities were used to create the parity transition matrices **U**_*i*_ and the fertility matrices **F**_*i*_, for each age class *i* (see equations 36 and 39).

Mortality schedules were extracted from the Human Mortality Database (2019). The age-specific period probabilities of death *q*_*x*_ were used to create the age advancement matrices **D**_*j*_ in equation (38).

The age-parity model produces a rich output data structure (Fig. 3), including the full age×parity structure, at every age of Focal, in every year, for each of the 14 types of kin in the network. Incorporating more types of kin into the network as described in Caswell (2019a) would extend this even further. So would incorporating dead as well as living kin, which we will encounter below in Section 7.

**Figure 3:**
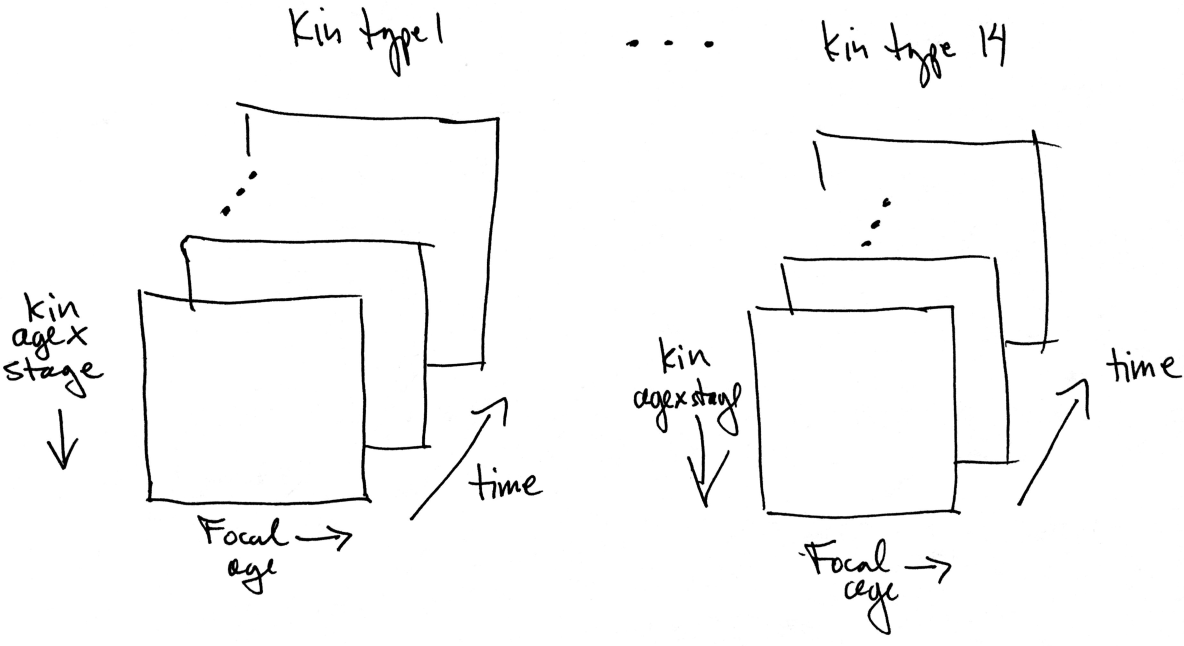
The data structure produced by the multistate kinship model applied to the kinship network of Fig. 1.

Unless your visualizing abilities are far better than average, exploring this five-dimensional (age of kin, stage of kin, age of Focal, year, type of kin) data structure requires simplifications. Here, we present

- a comparison of the joint age×parity structure at ages 20 and 60 of Focal in the years 1960 and 2014,
- the marginal parity structure over time (1960–2014) for ages 20 and 60 of Focal,
- the marginal parity structure as a function of the age of Focal, in 1960 and 2014,
- the marginal parity distributions of Focal and of her kin as a function of the age of Focal, in 1960 and 2014, and
- the proportion of low parity (0 or 1) kin over time (1960–2014), at ages 20 and 60 of Focal.

We will show results for selected types of kin.^4^ Mothers and daughters are presented because they are the kin most closely related to Focal. We have also included aunts; the role of aunts, especially childless aunts in providing support to their nieces has been the subject of much discussion. The internet proclaims that “aunts may be just as important as moms” and PANKs (professional aunts, no kids) are claimed to play a particularly significant role in the lives of their nieces and nephews.^5^

More rigorously, Nitsch, Faurie, and Lummaa (2014) found almost no effect of childless aunts and uncles on survival of nieces and nephews in 18th century Finland. Sear and Mace (2008) reviewed 45 studies of effects of kin on child survival and found no effects of aunts, but positive effects of maternal grandmothers and siblings.

As in the age-classified analysis of Japan in Caswell (2019a), these results should be taken as an example of what can be obtained from an age×parity model, not as a detailed analysis of the demography of Slovakia. For convenience, we will describe the results as applying to Slovakia in a specified year, rather than the more accurate but cumbersome description of a stable population experiencing the demographic rates of Slovakia in a specified year..

### 6.1 Age×parity structure

The model provides the age×parity structure of each type of kin, in each year, at each age of Focal (Figure 3). Results, for aunts at ages 20 and 60 of Focal, in 1960 and 2014, are shown in Fig. 4. From these age structures, one could calculate, e.g., the mean age within each parity class, the mean parity within each age class, and so on.

**Figure 4:**
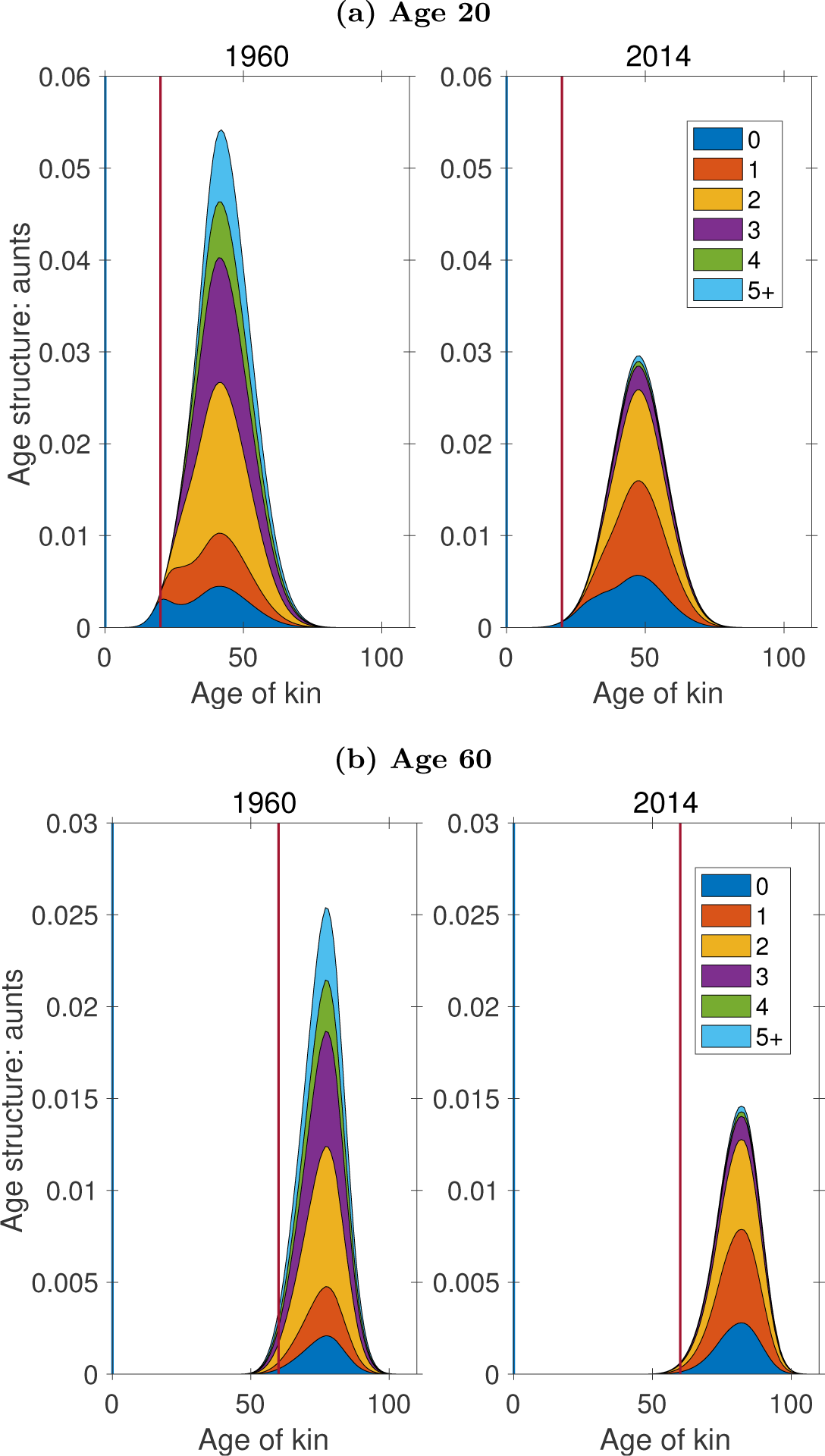
Expected age and parity structure of aunts, at ages 20 and 60 of Focal. Vertical line indicates age of Focal for reference.

In Slovakia the frequency of parity 0 aunts increased, while the frequency of high-parity aunts declined, between 1960 and 2014. We explore this pattern in more detail by examining the temporal dynamics of marginal parity distributions in the next section.

### 6.2 Parity structure over time

The marginal parity structure, obtained by applying equation (19) to the age×parity structure, gives the abundance of each parity class, integrated over age. Figures 5–8 show these parity structures for daughters, mothers, sisters, and aunts over time, at ages 20 and 60 of Focal.

**Figure 5:**
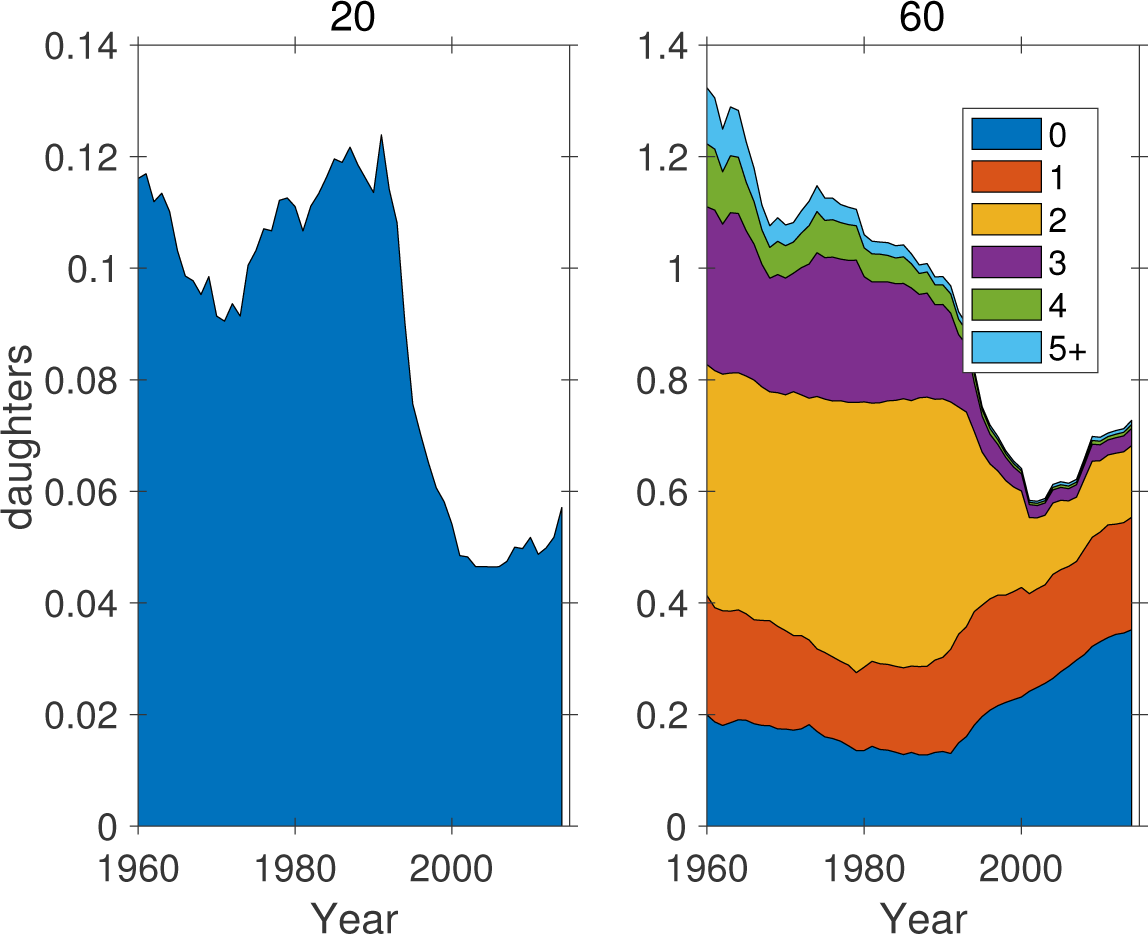
The marginal parity structure of daughters as a function of time, for ages 20 and 60 of Focal. Note different ordinate scales.

**Figure 6:**
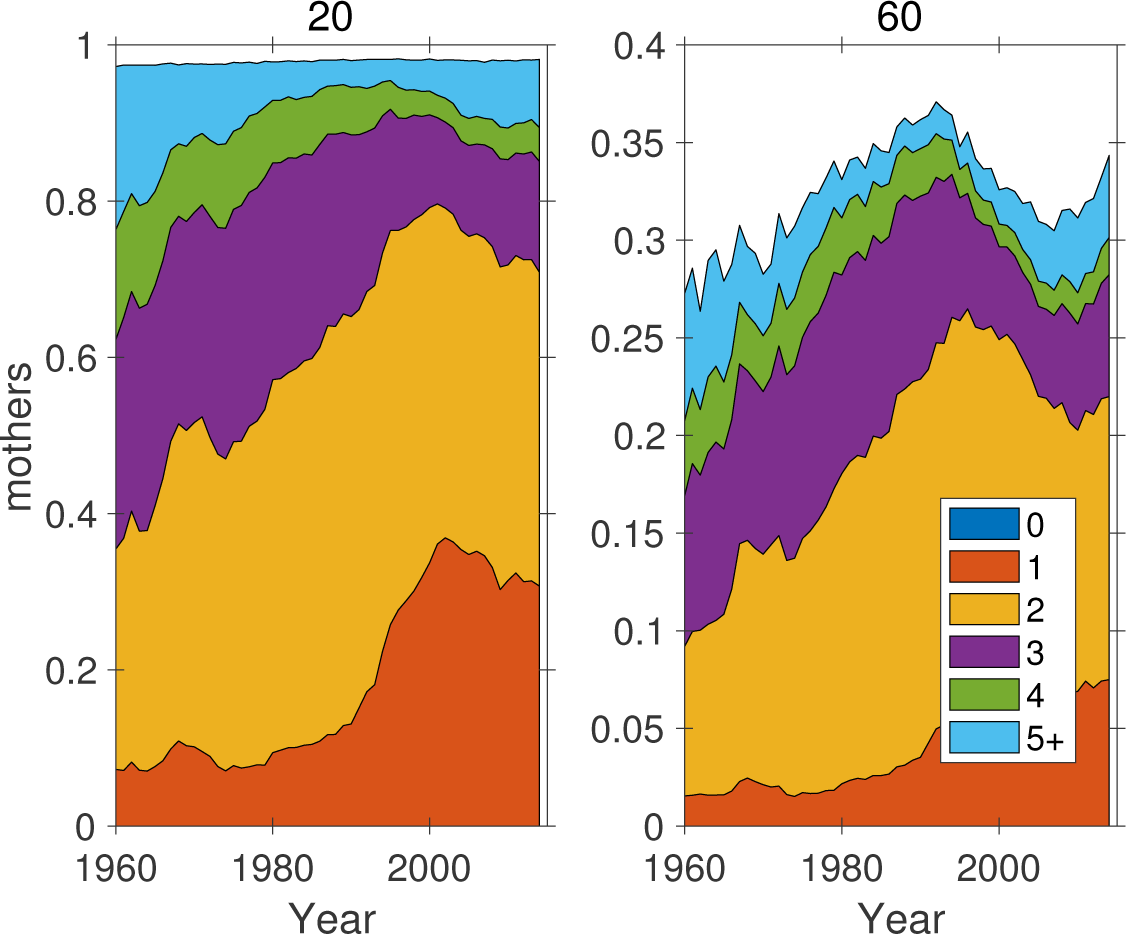
The marginal parity structure of mothers as a function of time, for ages 20 and 60 of Focal. Note different ordinate scales.

**Figure 7:**
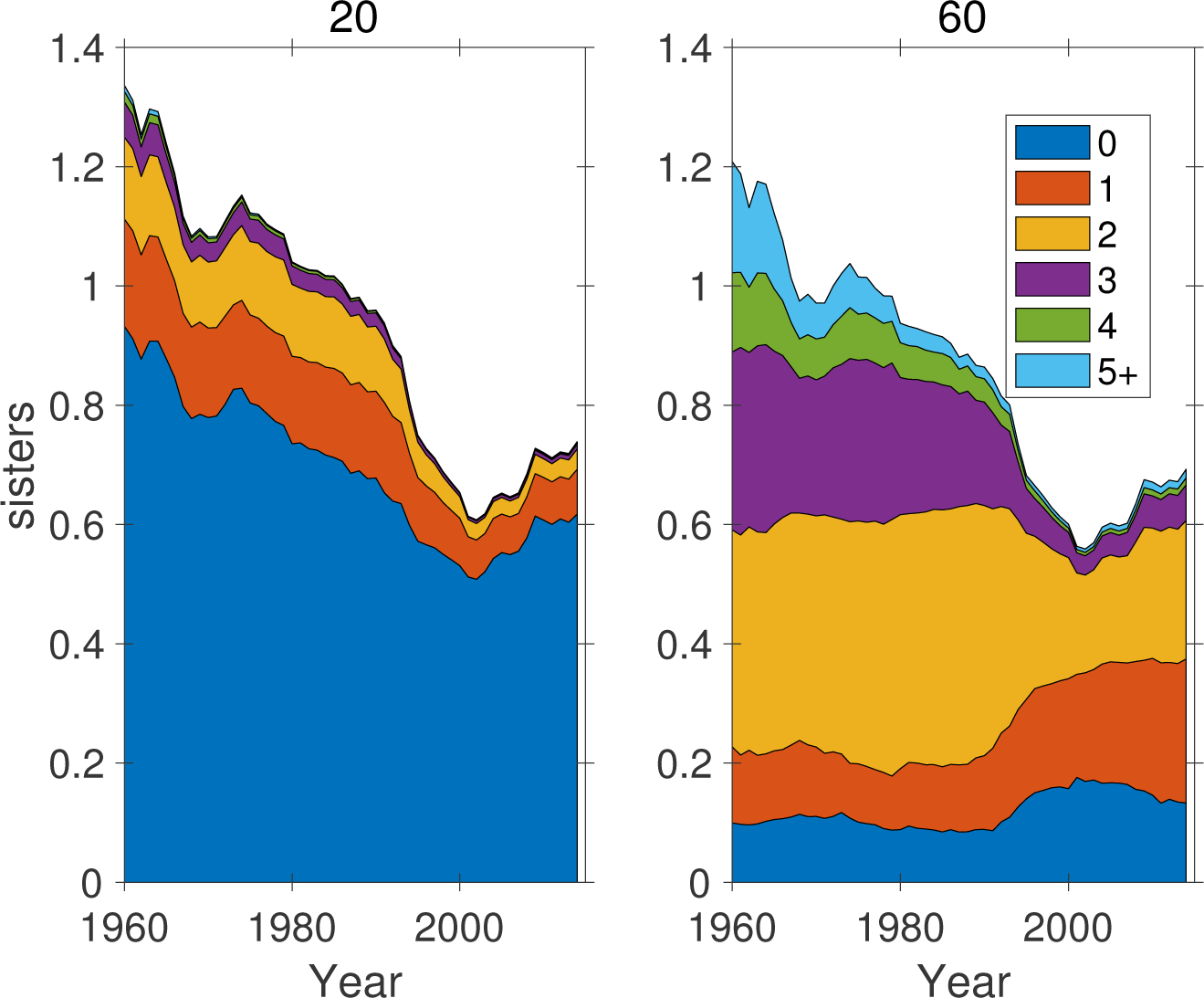
The marginal parity structure of sisters as a function of time, for ages 20 and 60 of Focal.

**Figure 8:**
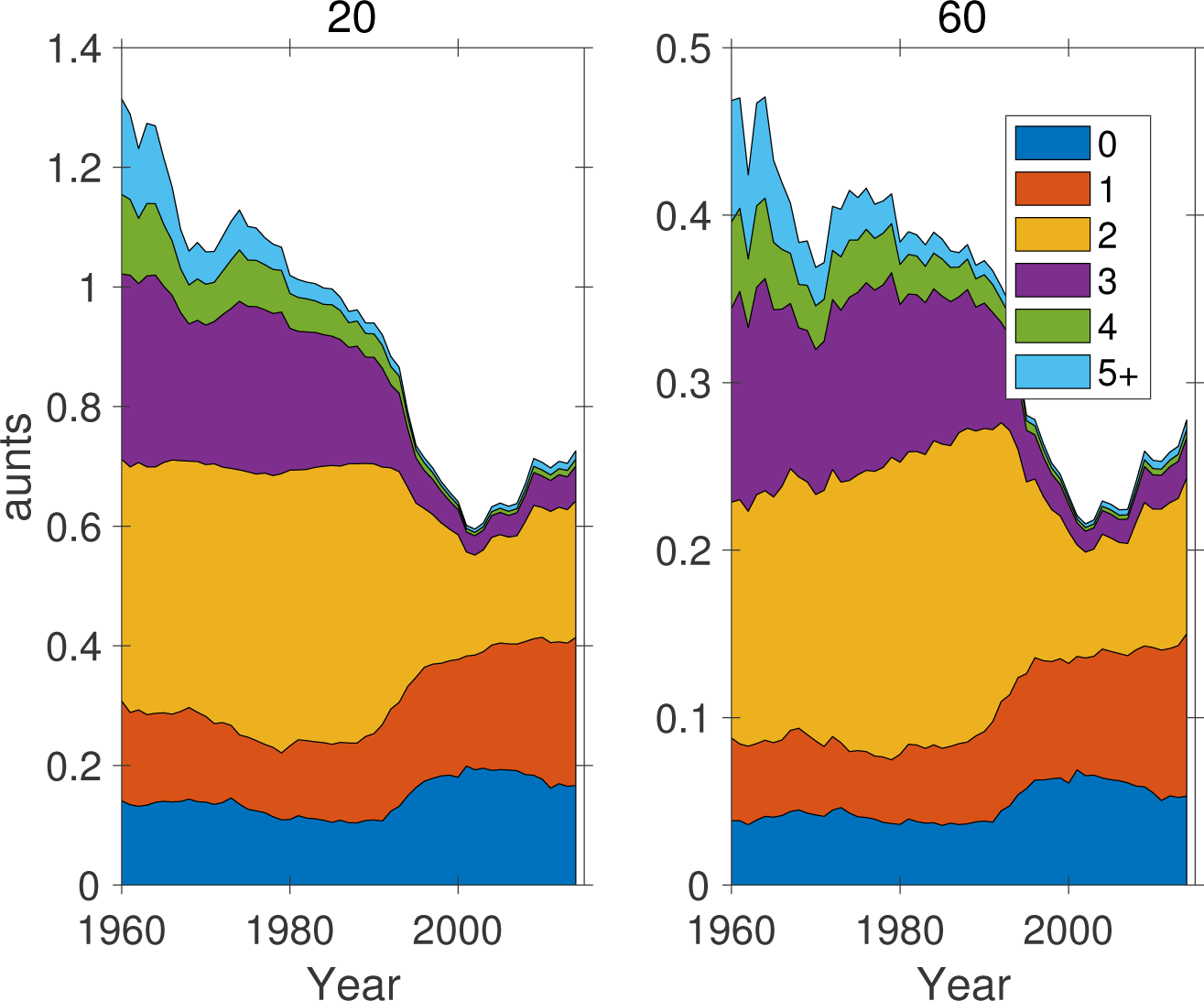
The marginal parity structure of aunts as a function of time, for ages 20 and 60 of Focal. Note different ordinate scales.

The number of Focal’s daughters at age 20 fell precipitously around 1990. At age 20 of Focal, her daughters have yet to reproduce and are all at parity 0 (Fig. 5). At age 60, the number of daughters of Focal follows the same pattern seen at age 20. However, we see that the decline in the number of daughters is accompanied by a dramatic shift in parity composition; by 2014, parity classes 3, 4, and 5+ have nearly disappeared.

A similar shift in parity composition is seen in the mothers of Focal, with an increase in parity classes 1 and 2 (Fig. 6). The same is true for the sisters and the aunts of Focal, which also decrease in abundance over time (Figs. 7 and 8).

### 6.3 Parity structure by age of Focal

Figures 9–12 show the marginal parity structures, in 1960 and 2014, as a function of the age of Focal. Under 2014 conditions, Focal has fewer daughters, and many fewer daughters in high parity stages, than in 1960. The survival of mothers is slightly better in 2014, and the reduction in high parity mothers is apparent. The pattern for sisters is similar to that for daughters. In 2014, Focal has only about half the number of aunts as in 1960, and again with fewer high parity individuals.

**Figure 9:**
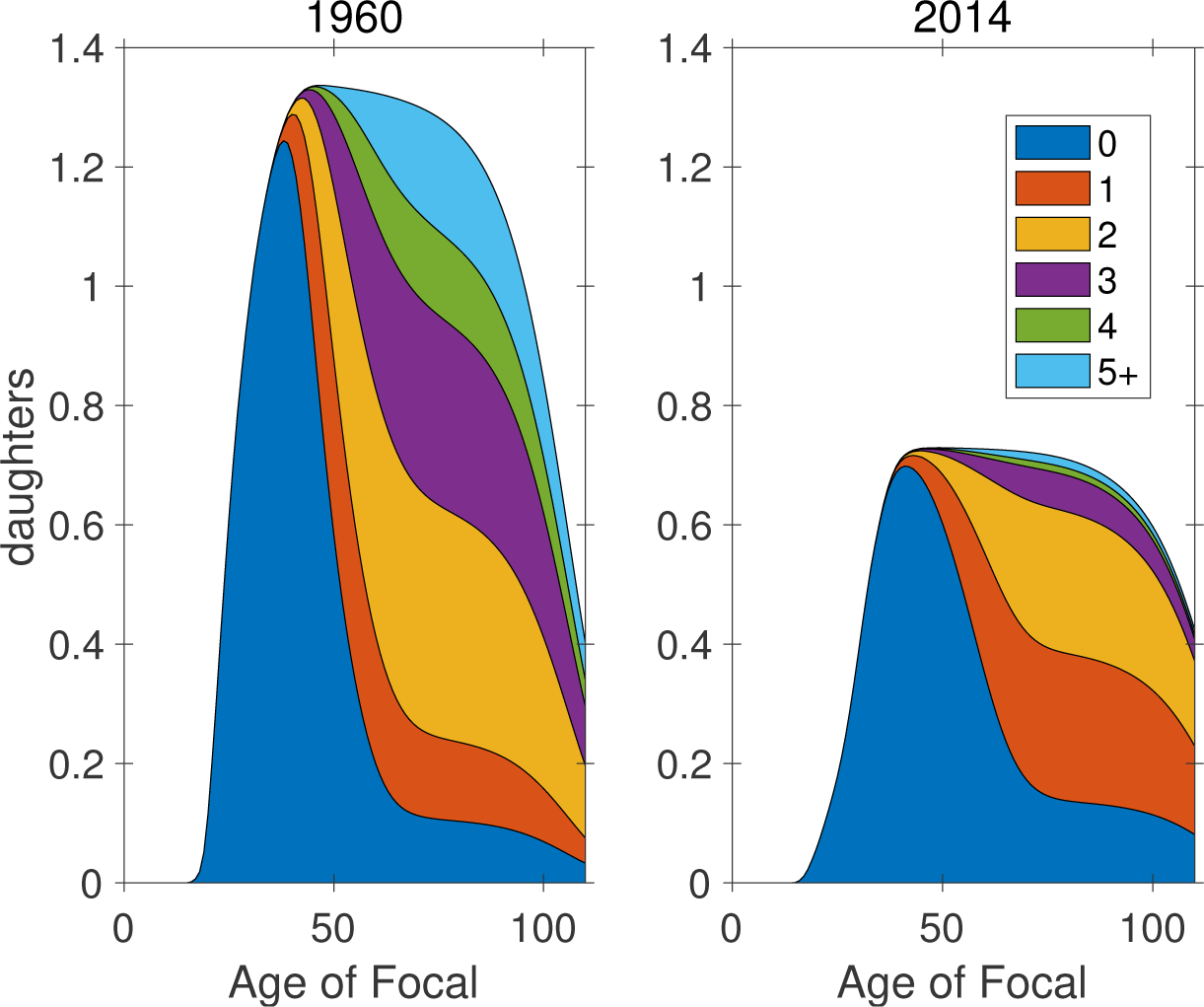
The marginal parity structure of daughters as a function of the age of Focal

**Figure 10:**
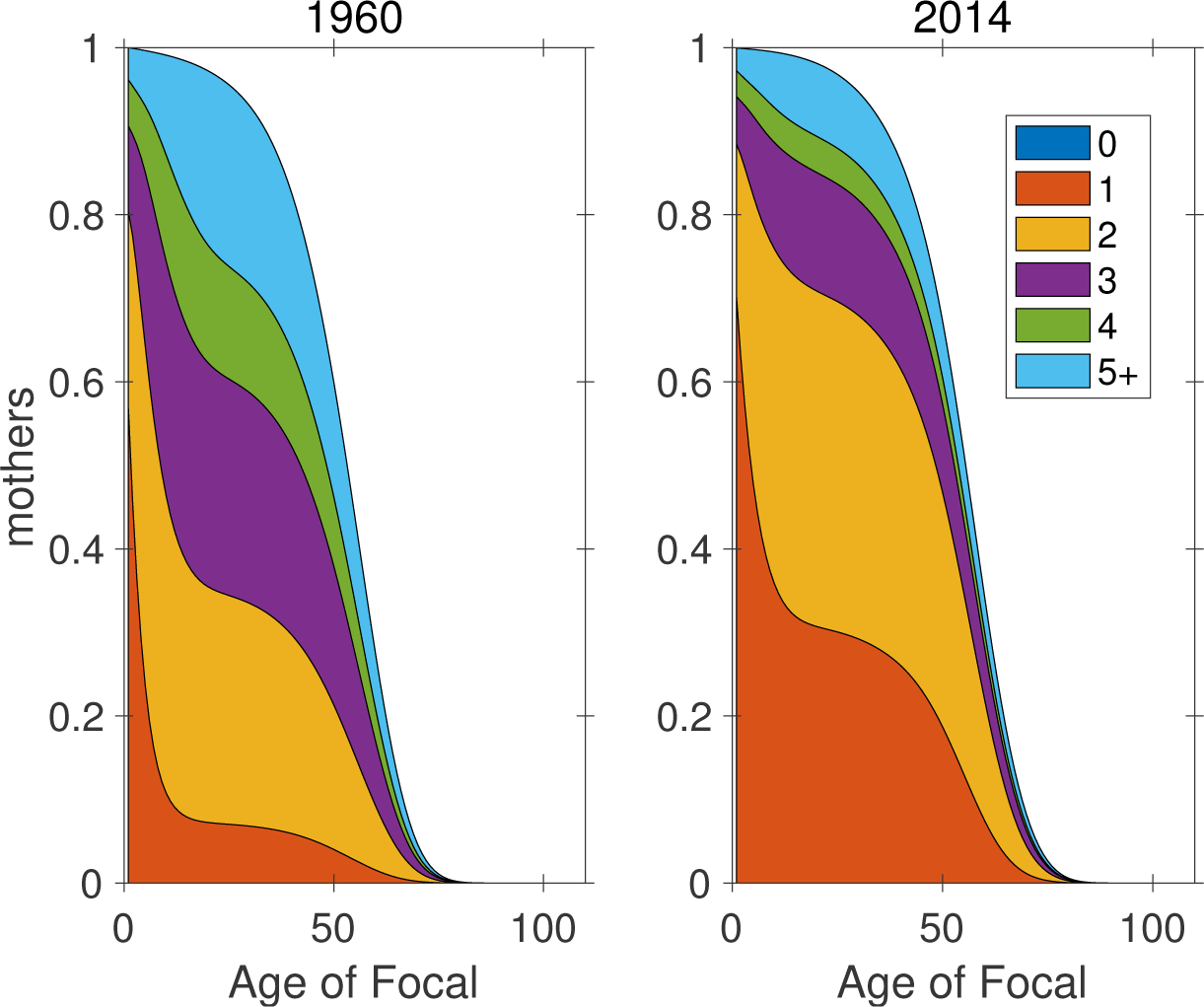
The marginal parity structure of mothers as a function of the age of Focal.

**Figure 11:**
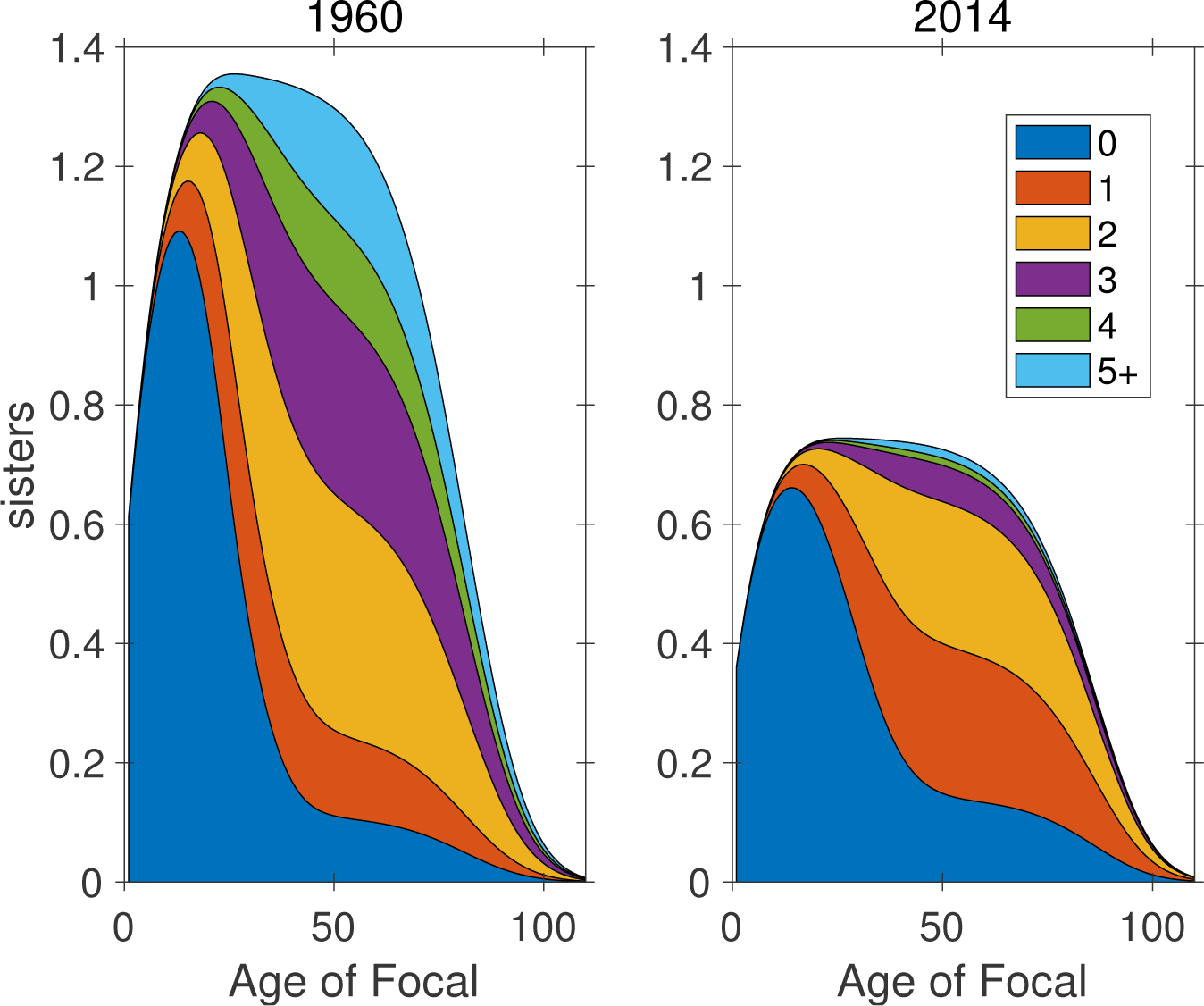
The marginal parity structure of sisters as a function of the age of Focal.

**Figure 12:**
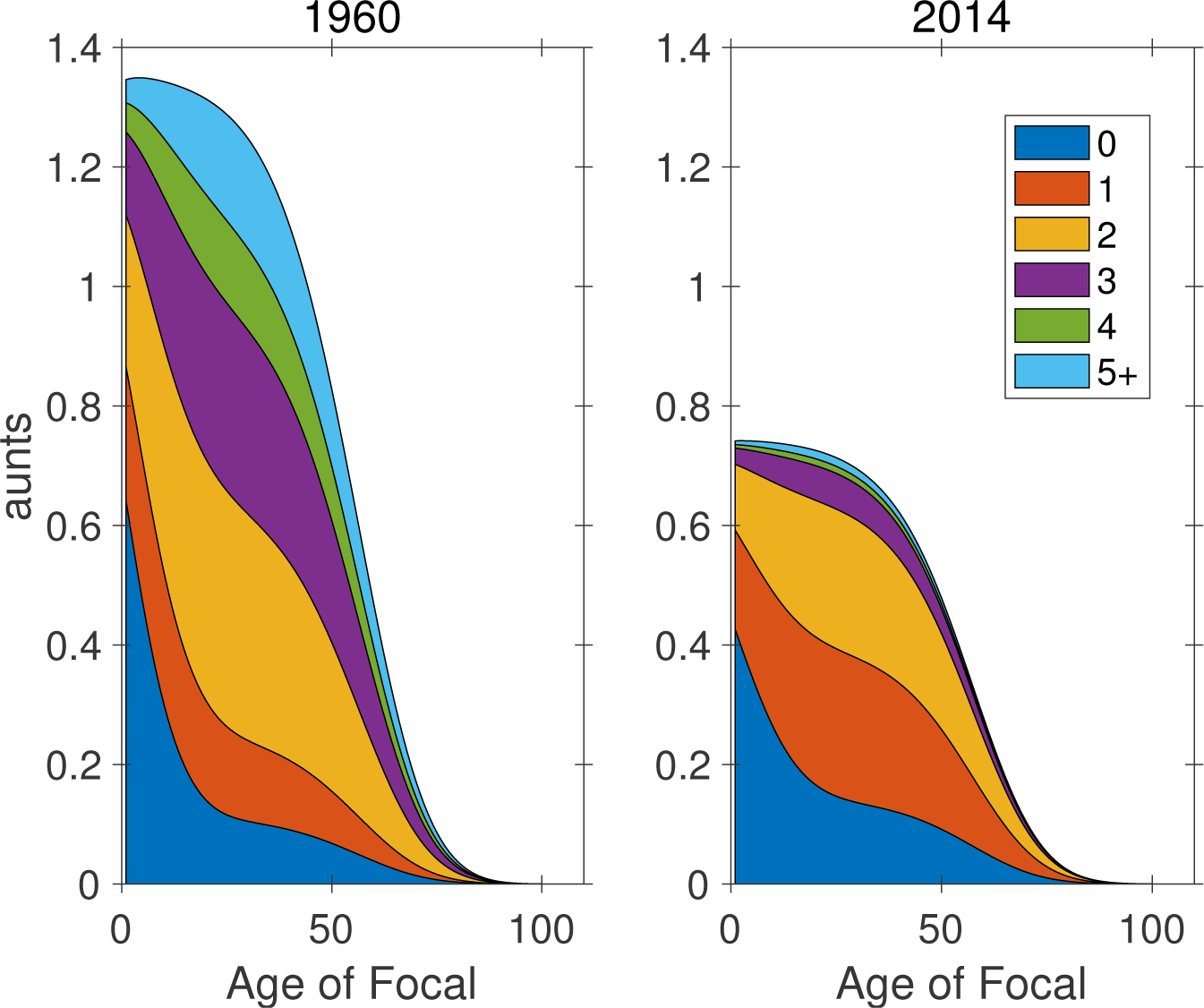
The marginal parity structure of aunts as a function of the age of Focal.

### 6.4 Parity distribution by age of Focal

From the previous figures, it is apparent that the parity composition of the kinship network in Slovakia changed dramatically between 1960 and 2014, and equally that it changes over the lifetime of Focal. This section focuses on the parity composition by plotting the marginal parity distributions (obtained by normalizing the marginal parity structure to sum to one) for selected kin.

Figure 13 presents the parity distribution of Focal herself. At birth, of course, Focal is in parity class 0. By age 50 she has stopped reproducing, and can no longer move among parity classes. Because mortality is independent of parity, the parity distribution remains fixed after this age. Between birth and age 50, the parity distribution gradually fills in, from 1 to 5+.

**Figure 13:**
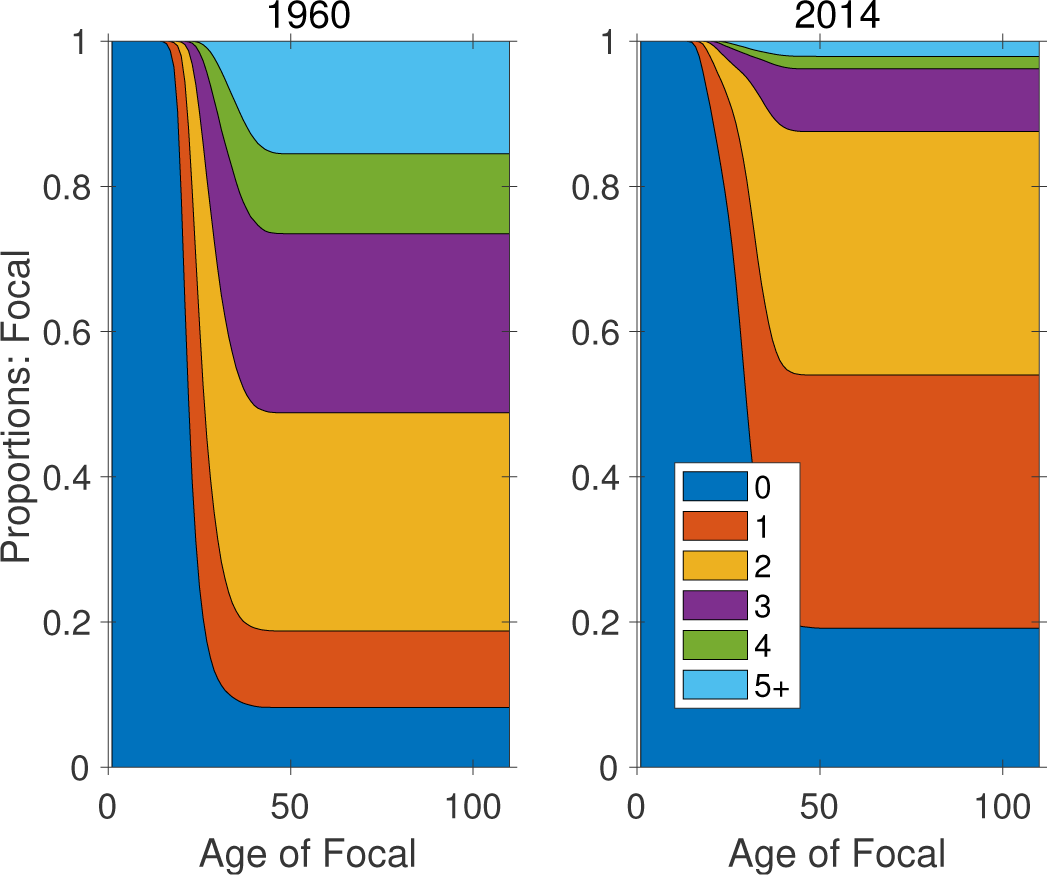
The marginal parity distribution of Focal as a function of age, in 1960 and 2014.

The probability of being at high parity declined strikingly between 1960 and 2014. The probability of parity 1, for example, increased more than three-fold, at the expense of parity 3, 4, and 5+.

Figures 14–17 show the marginal parity distributions of daughters, mothers, sisters, and aunts. In all four cases, there has been a dramatic increase in the proportion of low parity, and a decrease in the proportion of high parity kin. For example, at the age of 25, under 2014 rates, Focal is more than four times as likely to be an only child (i.e., for her mother to be in parity class 1) than was the case in 1960 (Figure 15). Similarly, sisters and aunts in parity classes 0 and 1 are almost three times as likely in 2014, compared to 1960.

**Figure 14:**
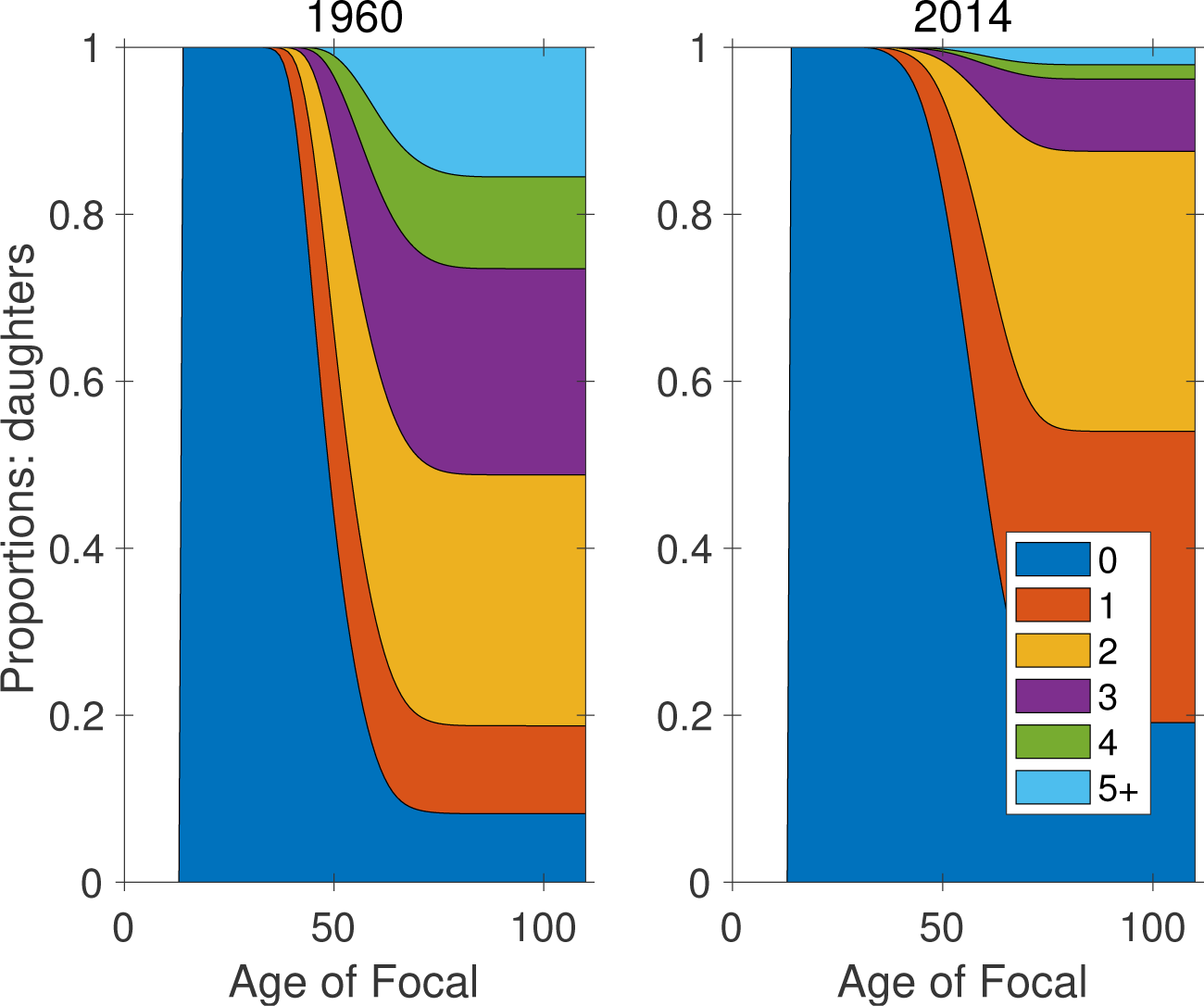
The marginal parity distribution of daughters as a function of age of Focal.

**Figure 15:**
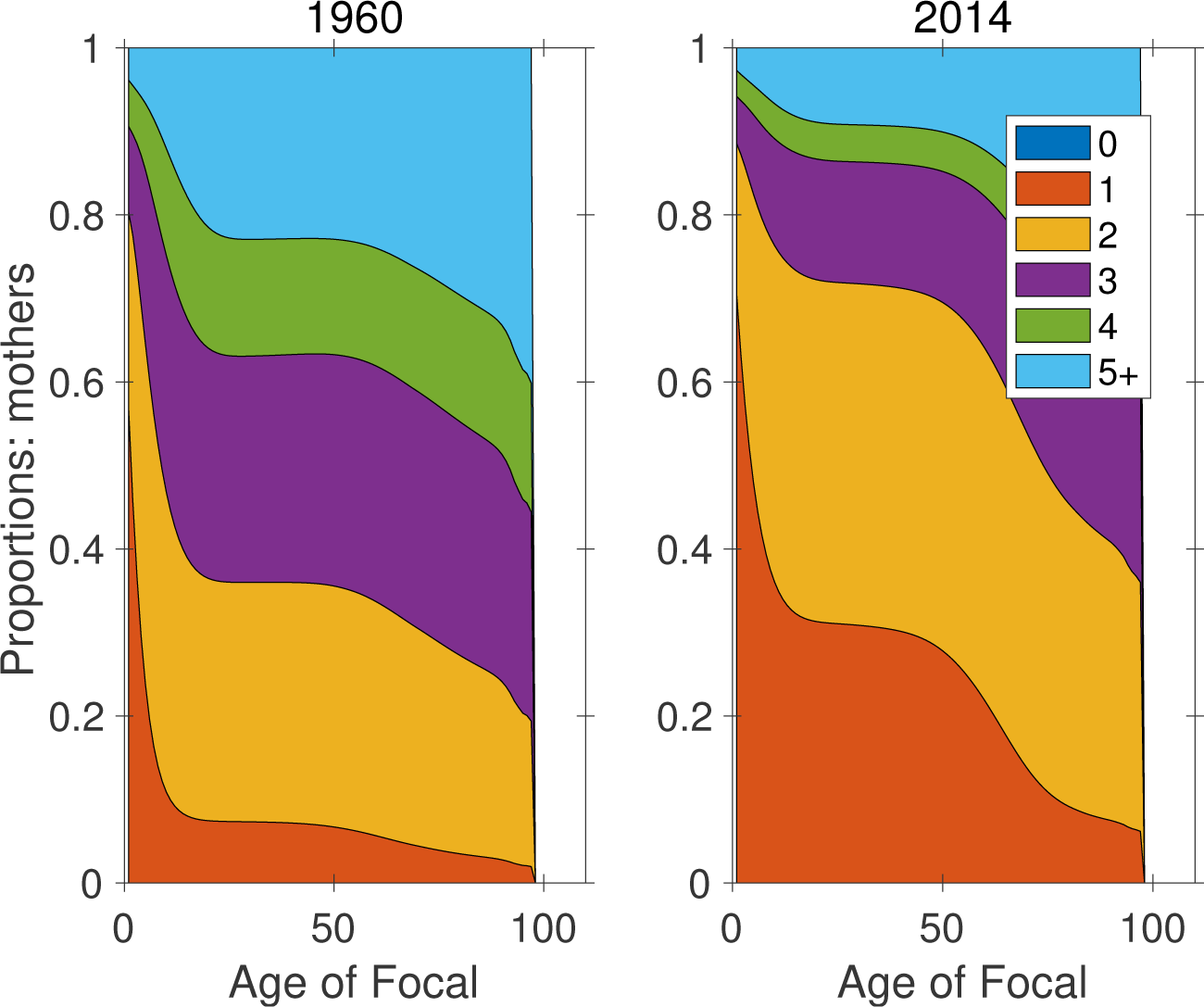
The marginal parity distribution of mothers as a function of age of Focal.

**Figure 16:**
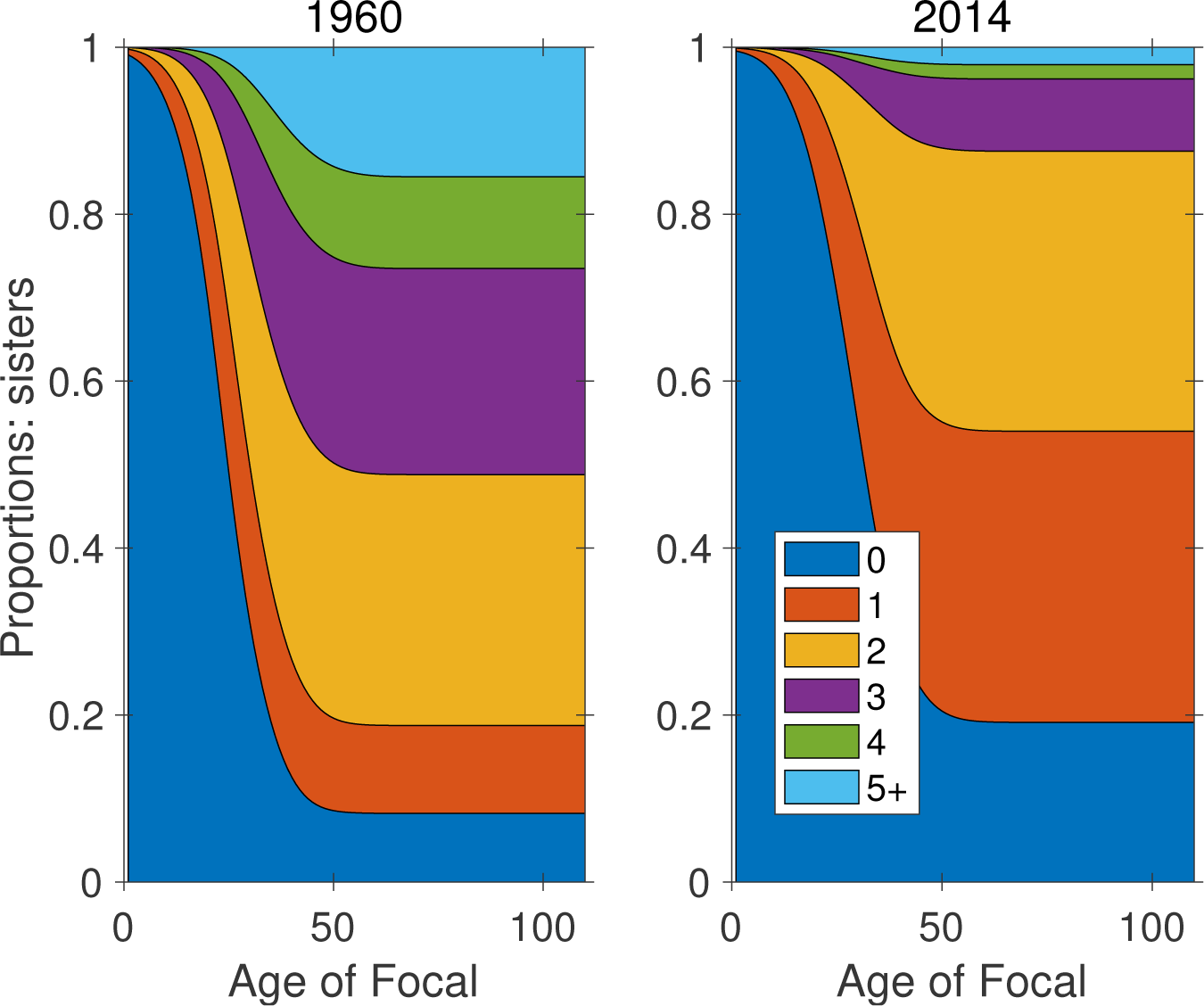
The marginal parity distribution of sisters as a function of age of Focal.

**Figure 17:**
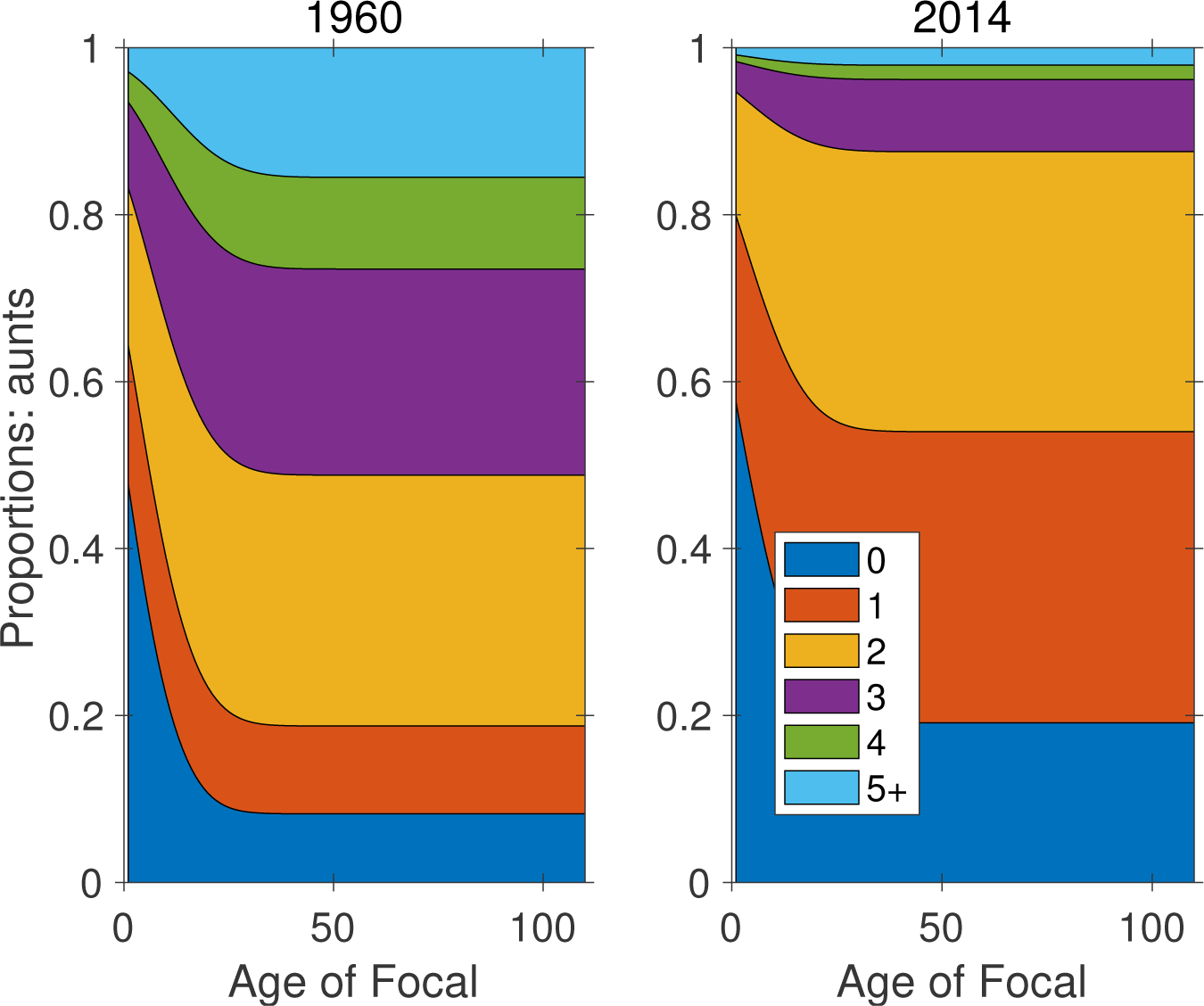
The marginal parity distribution of aunts as a function of age of Focal.

Figures 18 and 19 examine this parity shift in more detail, plotting the proportion of low parity (parity 0 or 1) sisters and aunts over time. Around 1990 there was a dramatic increase in the frequency of low parity kin.

**Figure 18:**
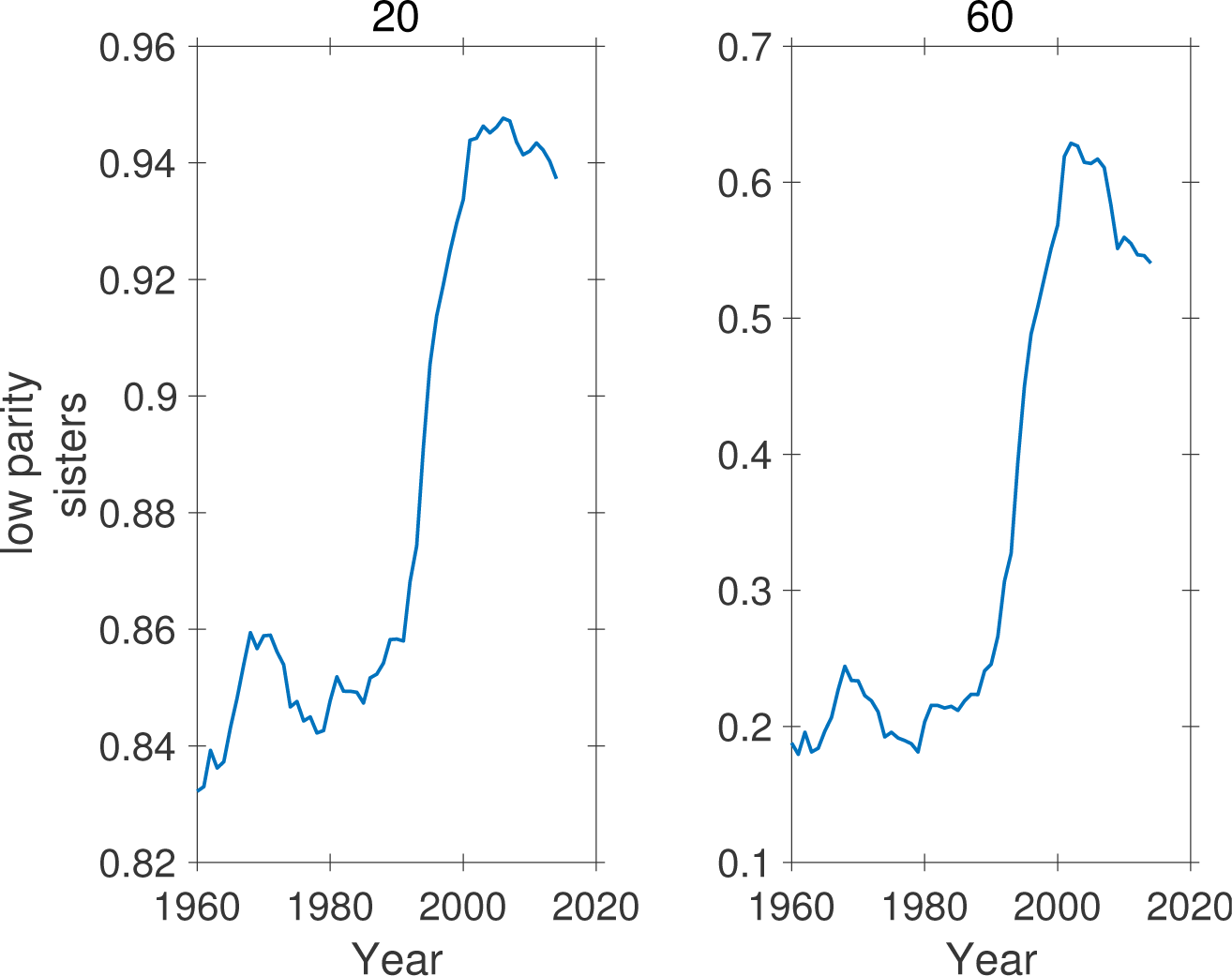
Proportion of low parity (parities 0 and 1) sisters of Focal as a function of time, for ages 20 and 60 of Focal. Note different ordinate scales.

**Figure 19:**
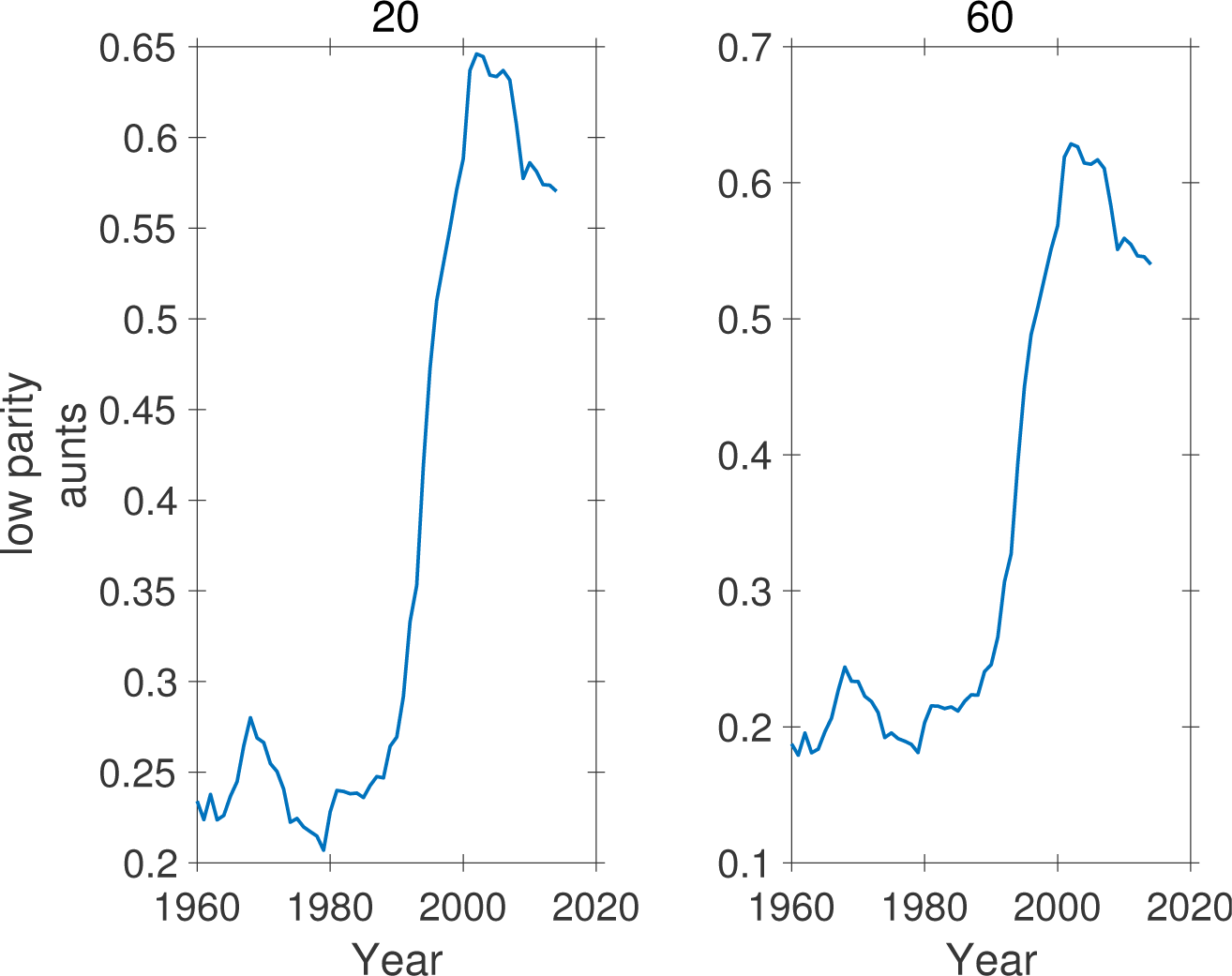
Proportion of low parity (parities 0 and 1) aunts of Focal as a function of time, for ages 20 and 60 of Focal. Note different ordinate scales.

## 7 Sibship size from parity

As pointed out by Schoen (2019a,b), the parity structure of a population provides important information on sibship size. We use the term sibship to refer to the children ever born in a family. The multistate model provides this information for any kind of kin, at any specified age of Focal; cf. the terms ‘family size’ (Preston, 1976) and ‘sibsize’ (Fahey, 2017; Schoen, 2019b).

The key to the calculation is the observation that the parity distribution in one generation gives (with some modifications, developed below) the sibship distribution in the next generation. For example, the probability that Focal is in parity class 2 tells something about the probability that the daughters of Focal are in a sibship of size 2. The probability that a daughter of Focal is in parity class 2 is the probability that the granddaughters of Focal are in a sibship of size 2. And so on.

Some previous analyses of sibship size have assumed that all women survive through their reproductive years (Preston, 1976; Schoen, 2019b). Our model requires no such assumption and incorporates mortality and its effects on sibship size.

To account for mortality, we note that the sibships among, say, the granddaughters of Focal include some whose mother (who is one of the children of Focal) has died. The chance of this event depends on the age of Focal and the mortality schedule to which she is subject (see the dramatic examples of the experience of death of relatives under the mortality and fertility schedules of Japan in Caswell 2019a).Thus the sibship size distribution of granddaughters is given by the marginal parity distribution of daughters, including daughters who have died. To include dead kin in the calculations we apply the approach of Caswell (2019a) to the age×parity-classified model.

### Notation alert

We are adding an additional dimension to the model (living vs. dead). This requires some additional notation. We have been using, e.g., **k** to represent an age structure vector, and 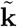 to represent an age×stage structure vector. Carrying on in the same way, we define 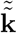, with two tildes^6^, to represent an age-stage-living structure vector, which includes both living and dead,

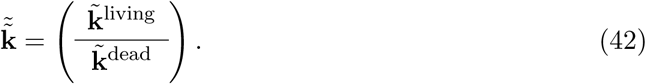

That is, 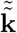 is a block-structured vector whose blocks (living and dead) are themselves block-structured (age×stage) vectors. Following Caswell (2019a), the dynamics of 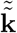 are given by

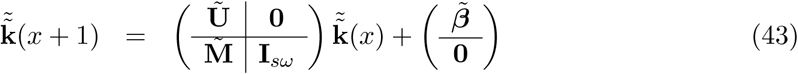

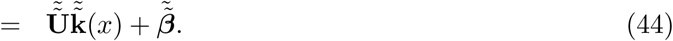

where **Ũ** is given by equation (12) and 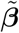 is the age×stage-classified subsidy term appropriate for the given type of kin, as given in Table 2. The **0** matrix in the upper right quadrant of 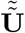 insures that dead kin to not return to life. The mortality matrix 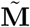 collects the dead kin into absorbing states corresponding to their age and stage at death, and is given by

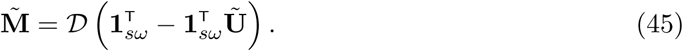

The identity matrix **I**_*sω*_ in the lower right quadrant traps the dead kin in the age-stage combination in which they died.

The fertility matrix is

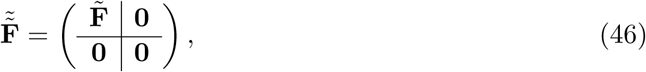

showing that fertility produces only living kin, and that dead kin do not reproduce.

Given the structure vector 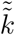 in (42), the age×stage structure vector of kin ever born is obtained by summing the living and dead kin,

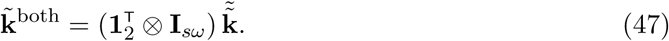

The marginal age and stage structures, combining both living and dead kin, are

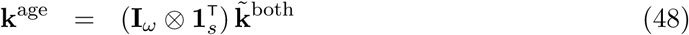

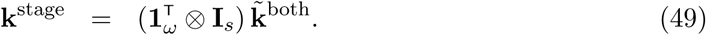

The marginal parity distribution, **k**^stage^(*x*)*/*‖**k**^stage^(*x*)‖ of kin of type **k** gives the proportional sibship size distribution of the offspring of those kin, at age *x* of Focal.

A few examples are shown in Figures 20–22, comparing the sibship size distributions in 1960 and 2014. The sibship size distribution of Focal herself, obtained from the parity of Focal’s mother, has no support for size of 0, because Focal is assumed to be exist, meaning her sibship size is at least 1 (Figure 20). By the time Focal has reached the age of about 25 years, her sibship size distribution has stabilized (her mother being unlikely to reproduce after that age). In 1960 the probability of a sibship size of 1 or 2 was about 0.4; in 2014 that probability had almost doubled.

**Figure 20:**
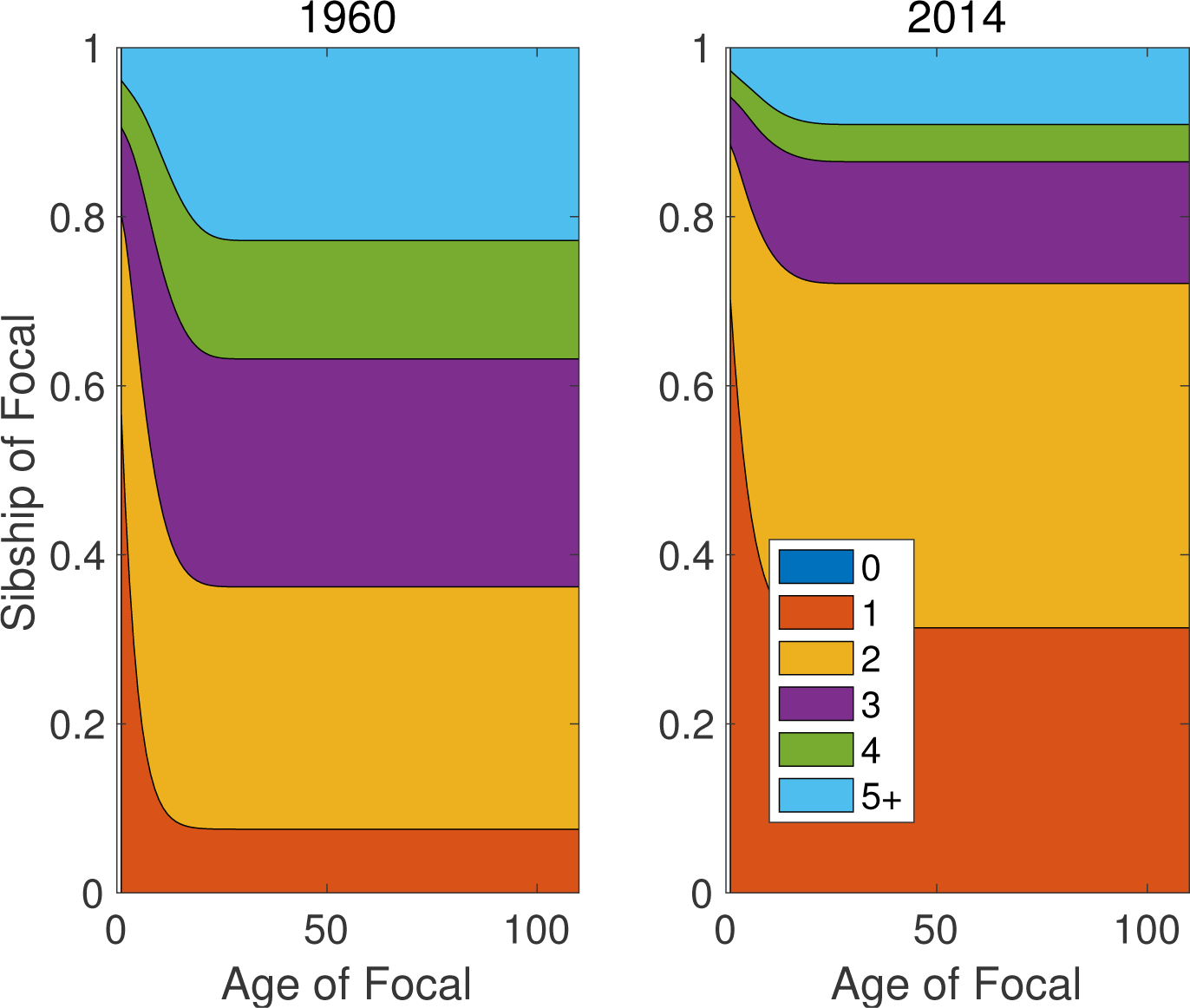
The distribution of the size of the sibship of Focal, as a function of the age of Focal, for years 1960 and 2014.

**Figure 21:**
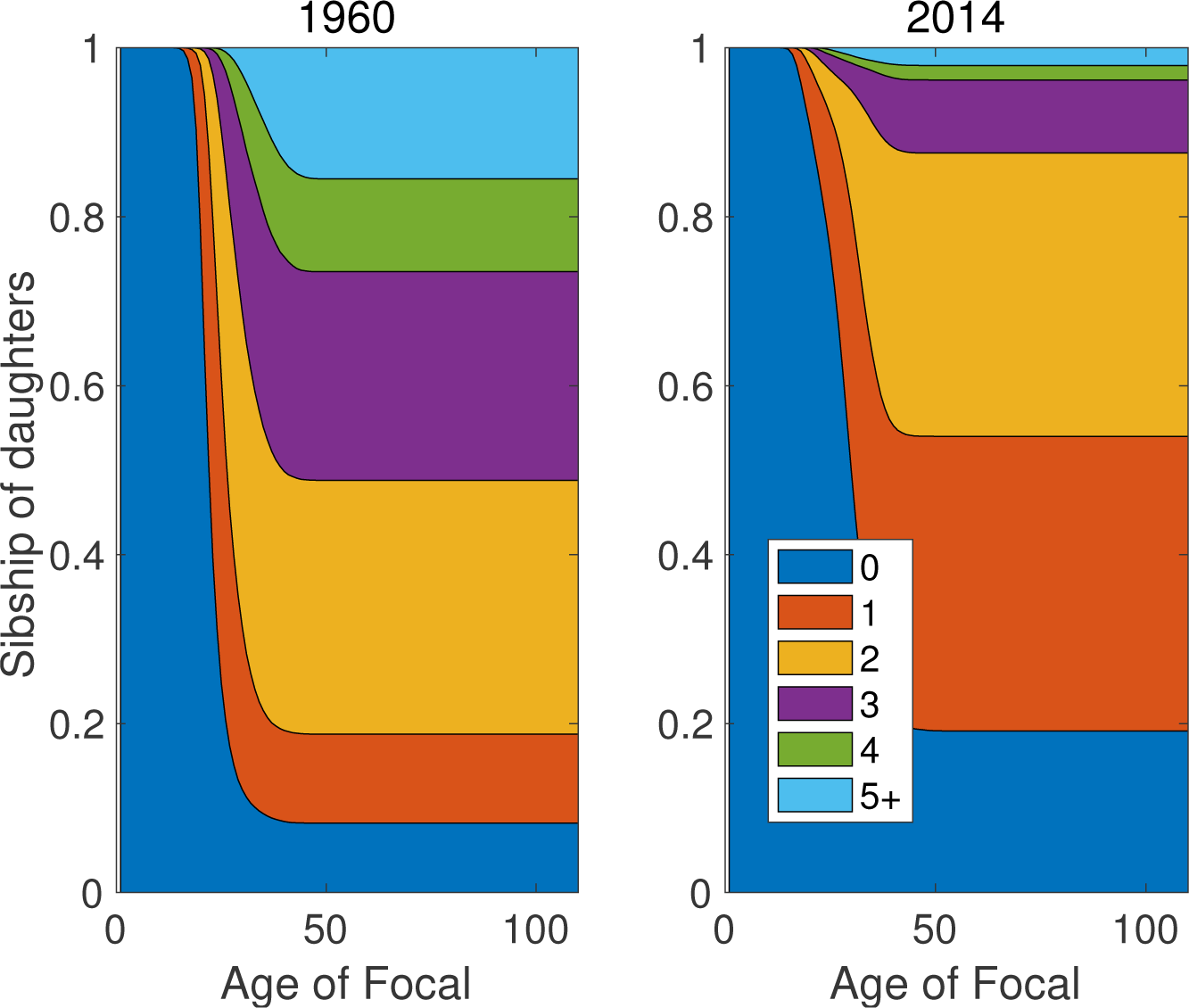
The distribution of the size of the sibship of the daughters of Focal, as a function of the age of Focal, for years 1960 and 2014.

**Figure 22:**
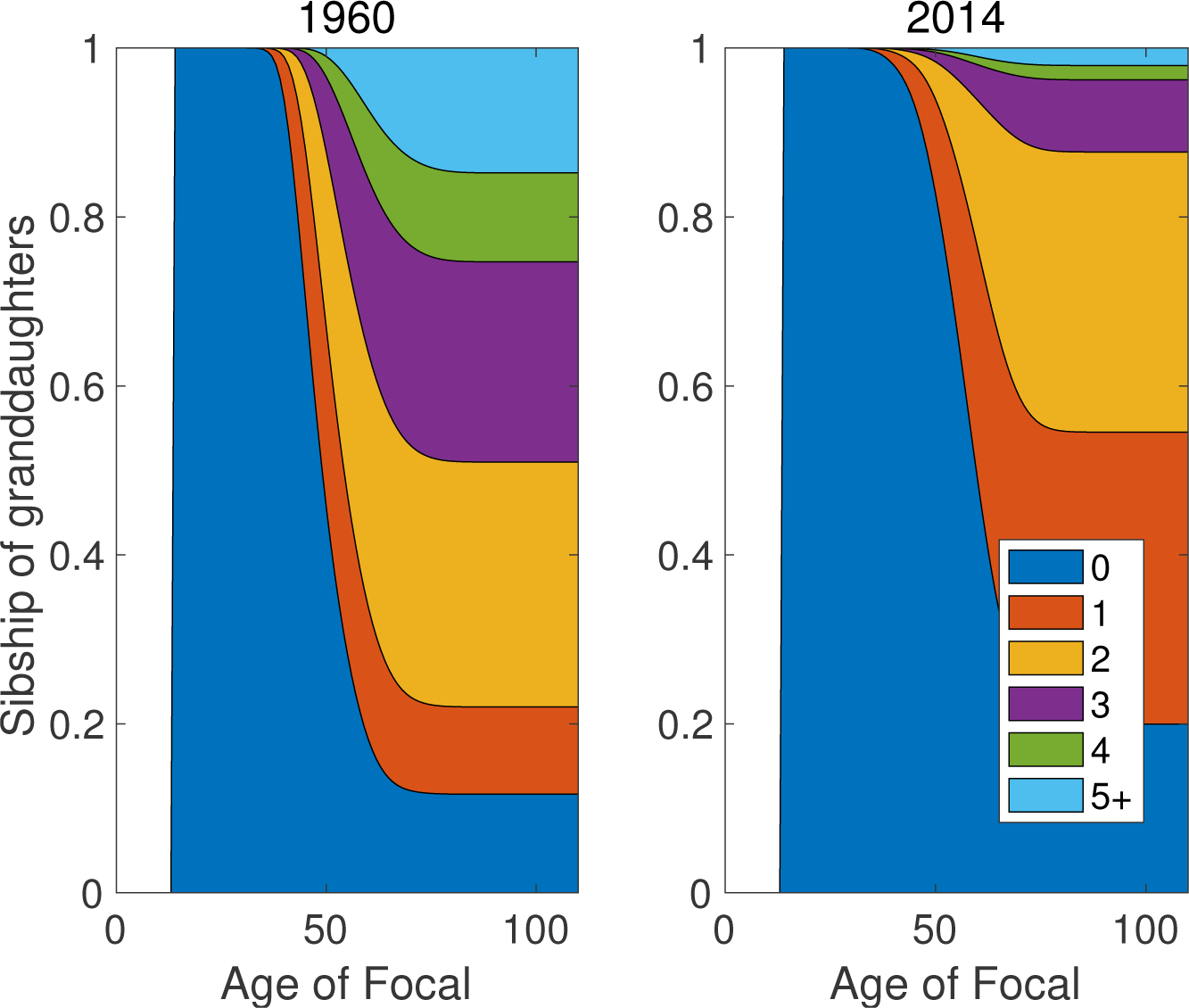
The distribution of the size of the sibship of the granddaughters of Focal, as a function of the age of Focal, for years 1960 and 2014.

A similar dramatic change in sibship size of the daughters and granddaughters of Focal is apparent in Figures 21 and 22. The probability of sibship sizes 0 or 1 more than doubled from 1960 to 2014, accompanied by a great decrease in the probability of sibships of 4 or 5+.

In populations with low mortality, the effects of including dead kin will be small. Compare, for example, the parity distribution of the daughters of Focal in Figure 14, calculated without incorporation of death, to the sibship size distribution of the granddaughters of Focal in Figure 22.

## 8 Discussion

Any individual is surrounded by a network of kin. The ages and other properties of those kin have important implications for intergenerational interactions, family sizes, and genetic relatedness. The kinship network is shaped by the mortality and fertility schedules of the population; all else being equal, higher mortality leads to fewer kin, higher fertility to more kin. Goodman, Keyfitz, and Pullum (1974) formulated kinship calculations by asking about the kin of a focal individual of a specified age. This Focal-centric view makes the kinship network a property of the individual. Caswell (2019a) took advantage of this perspective to formulate the kinship network as a collection of populations (the populations of the daughters, the sisters, the cousins, etc. of Focal). Projecting the sizes and age structures of those populations gives detailed insight into the kinship network. Age, however, is not the only game in town, and age-specific mortality and fertility schedules do not necessarily capture all aspects of demography. The matrix-based analysis introduced here makes it possible, for the first time, to take a multistate view of kinship, classifying individuals by age and one or more additional state variables.

This study reports results for an age×parity model, but parity is not the only kind of stage structure that could be included. The model could also be used to study the interaction of age and relationship status, using models like those of Mills (2000) for stage transitions. It also provides an approach to two-sex kinship models, accounting for male and female kin through male and female lines of descent. Characteristics like maternal age can be incorporated using a model like that of Hernández et al. (2019). We have not explored them here, but the same kinds of kin properties investigated by Caswell (2019a) can be applied directly to multistate models (e.g., prevalence of diseases, dependency ratios, mean age (and now also mean parity), experiences of the death of kin).

It is important to think carefully about what this model and its results imply. As emphasized by Goodman, Keyfitz, and Pullum (1974), formal demographic calculations of kinship assume that mortality and fertility schedules apply uniformly to everyone, and have been in operation long enough to make stable population theory useful. The results are not expected to agree with results of a census or sample of a population. Such a census will also reflect the history of changes in those schedules and distortions of the population structure. As with other counterfactual assertions in demographic theory (about homogeneity, or time invariance, or stability) the value of such formal analyses is that they reveal the interaction of factors that influence kinship and provides a background against which to interpret census results.^7^ Comparisons of the model calculations with population counts will be useful in the same way that comparison of observed population properties with stable population calculations reveal the effects (bulges and gaps in the age structure, fluctuations in population growth) of the violation their counterfactual assumptions. Relaxing the assumption of time-invariant rates in the kinship model remains an important open problem.

The model presented here is based on deterministic projections of the kin of Focal. As a result, the age×stage structures produced by the model, and reported here, are expected values. But because the populations of kin are small, demographic stochasticity will generate variance around those expected values. Incorporating demographic stochasticity, perhaps using generalizations of multitype branching processes (Pollard, 1973; Caswell, 2001; Caswell and Vindenes, 2018) will yield important information on the covariance structure of ages and stages of kin. Pullum and Wolf (1991) have suggested such an approach in the Goodman-Keyfitz-Pullum model.

The vec-permuation matrix approach used to construct the multistate models here is a powerful method for integrating the joint action of age and other variables (Caswell et al., 2018). In the present context, one of its advantages is that it leads to a matrix formulation that parallels exactly the age-classified kinship model of Caswell (2019a). Because there are two dimensions rather than one, the ability to go easily from the full joint age×stage structure to the marginal age and stage structures, via equations (18) and (19), is particularly useful.

The model presented here provides a basic theoretical structure for the demography of kinship. Like any theory, it identifies a set of processes (survival, fertility, the concept of subsidized production of kin, stage transitions) in terms of which to analyze the system. It provides a framework linking the phenomena to other theoretical constructs, particularly the theory of population projection. And like any theory, it clarifies the gaps in the theoretical structure. These include, inter alia, time variation, two-sex analyses, and stochasticity. The theory provides hints, sometimes very strong hints, about how to fill those gaps. Burch (2018) has argued that models form the core of theory in demography, and we believe that this kinship model is a clear example.

The analysis here of data from Slovakia is a non-exhaustive sample of ways to explore the kinship network (joint and marginal age and parity structures, dynamics of numbers and parity structures over time and over the lifetime of Focal, parity distributions, sibship size distributions). The output data structure (Figure 3) suggests additional possibilities. This diversity of endpoints emphasizes the additional richness of the model compared to the age-classified case.

Between 1960 and 2014, fertility and mortality in Slovakia both declined (TFR dropped by about 60%; life expectancy increased by 28%). This was accompanied by steep declines in the numbers of most types of kin (daughters, granddaughters, great-granddaughters, sisters, nieces, aunts, cousins). Even more dramatic, however, were the changes in parity composition of kin. An examination of the marginal parity composition (Figs. 5 – 8) shows that the decline in numbers of kin was accompanied by a great reduction in the abundance of high parity kin. A comparison of marginal parity distributions over the lifetime of Focal at the beginning and end of the time period shows the same (Figs. 9 – 12). The proportion of low parity individuals among the kin showed a dramatic increase, by several fold, around 1990 (Figs. 18 – 19).

Because the parity distribution of kin gives the sibship size distribution of the offspring of those kin, these results imply that the changes in the vital rates of Slovakia are capable of generating a sibship size revolution similar to that documented in the U.S. by Fahey (2017). It remains to be seen what kinds of patterns will emerge from the application of multistate kinship analysis to other situations.

## 9 Acknowledgments

This research was supported by the European Research Council under the European Union’s Horizon 2020 research and innovation program through ERC Advanced Grant 788195. Motion Coffee in Amsterdam provided a pleasant environment. I thank Robert Schoen for a helpful comment on parity and sibship. Discussions with Xi Song, Silke van Daalen, and the Theoretical Ecology group at the University of Amsterdam were helpful throughout.

## Appendix A Derivation of the multistate kinship equations

The derivation of the model for of each of the 14 types of kin shown in Figure 1 are given here. The derivations follow, as closely as possible, those in Caswell (2019a), and hence have been relegated to this Appendix. To keep the presentation as parallel as possible to the age-classified model, the text of this Appendix has been modified from Section 2.1 of Caswell (2019a) under the terms of a Creative Commons Attribution License.

The new aspects of the derivations, required to incorporate stage- as well as age-classification, are (1) the use of age×stage structure vectors (distinguished with a tilde; e.g., **ã**) in the place of age structure vectors (e.g., **a**), (2) the use of the marginal age distribution ***π***^age^ of the mothers of children in the stable population, which is calculated from the joint distribution 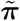 of ages and stages of those mothers, and (3) the calculation of the age×stage structure vector 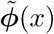 of Focal herself, which appears in Section 4.2.1.

### A.1 Daughters and descendants

Each type of descendent depends on the reproduction of another type of descendent, or of Focal herself.

**ã**(*x*) **= daughters of Focal.** Daughters are the the children of Focal. Focal is assumed to be alive at age *x*, with an age×stage distribution given by 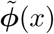. The subsidy vector for daughters is obtained by applying the fertility matrix 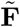 to that age×stage distribution, so 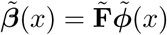. We know that Focal has no daughters when she is born, so the initial condition is **ã**_0_ = **0**. Thus

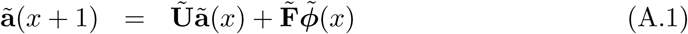

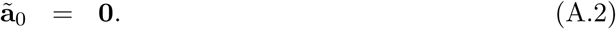

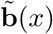 **= granddaughters of Focal.** Granddaughters are the children of the daughters of Focal. At age *x* of Focal, those daughters have an age×stage distribution **ã**(*x*), so the subsidy term for granddaughters is 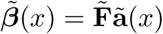. Because Focal has no granddaughters at birth, the initial condition is **0**; thus

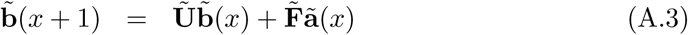

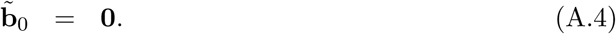

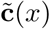 **= great-granddaughters of Focal.** Similarly, great-granddaughters are the daughters of the granddaughters of Focal, with a subsidy term 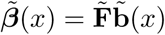, and an initial condition of **0**.

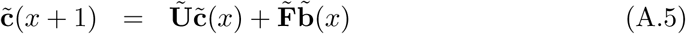

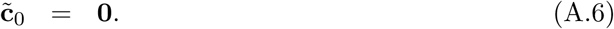

The extension to arbitrary levels of direct descendants is straightforward. Let 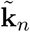, in this case, be the age×stage distribution of descendants of level *n*, where *n* = 1 denotes daughters. Then

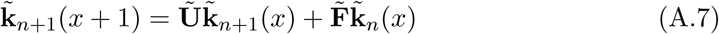

with the initial condition 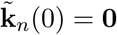.

### A.2 Mothers and ancestors

The populations of surviving mothers and other direct ancestors of Focal depend on the age of those ancestors at the time of the birth of Focal.

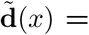 **mothers of Focal.** Focal has at most one mother (step-mothers are not considered here). Her expected age×stage structure at age *x* of Focal is denoted by 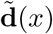. No new mothers appear after Focal’s birth, so the subsidy term is 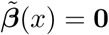.

The age of the mother of the newborn Focal is unknown, but we know the distribution of her age. The corresponding age×stage distributions must be conditioned on parity of 1 or greater. To do so, define the matrix 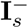 to be an identity matrix of order *s* with a 0 in the (1, 1) entry. Then the age×stage distribution given in equation (29) is replaced by the conditional distribution

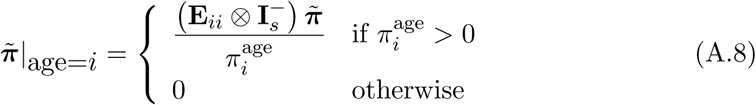

and the initial condition for mothers is given by (31)

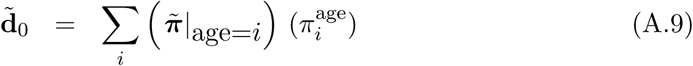

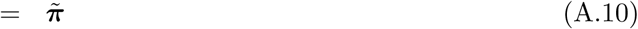

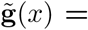 **grandmothers of Focal.** The grandmothers of Focal are the mothers of the mother of Focal. No new grandmothers appear, so the subsidy term ***β***(*x*) = **0**. The age structure of grandmothers at the birth of Focal is the age structure of the mothers of Focal’s mother, at the age of Focal’s mother at Focal’s birth. The age of Focal’s mother at Focal’s birth is unknown, so the initial age structure of grandmothers is a mixture of the age×stage structure 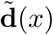 of mothers, with mixing distribution ***π***^age^:

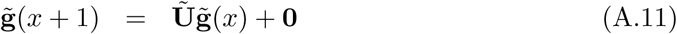

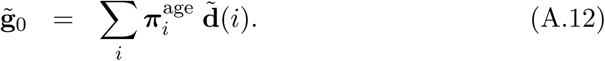

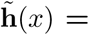 **great-grandmothers of Focal.** The great-grandmothers of Focal are the grandmothers of the mother of Focal. No new great-grandmothers appear, so the subsidy term is ***β***(*x*) = **0**. The initial condition is a mixture of the age×stage structures 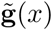 of the grandmothers of Focal, with mixing distribution ***π***^age^:

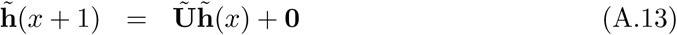

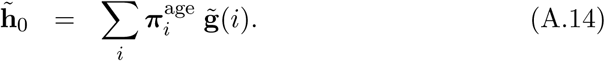

The extension to arbitrary levels of direct ancestry is clear. Let 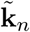 be, in this case, the age structure vector of ancestors of level *n*, where *n* = 1 denotes mothers. Then the dynamics are

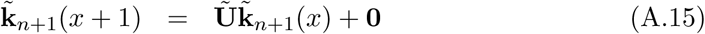

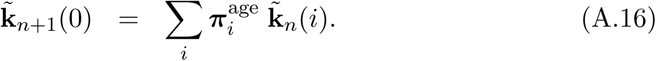

#### A.2.1 Sisters and nieces

The sisters of Focal, and their children, who are the nieces of Focal, form the first set of side branches in the kinship network of Figure 1. Following Goodman, Keyfitz, and Pullum (1974), we divide the sisters of Focal into older and younger sisters, because they follow different dynamics.

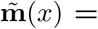 **older sisters of Focal.** Once Focal is born, she accumulates no more older sisters, so the subsidy term is ***β***(*x*) = **0**. At Focal’s birth, her older sisters are the children **ã**(*i*) of the mother of Focal at the age *i* of Focal’s mother at Focal’s birth. This age is unknown, so the initial condition 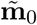 is a mixture of the age structures of daughters with mixing distribution ***π***^age^.

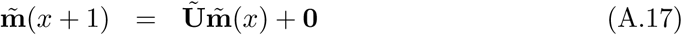

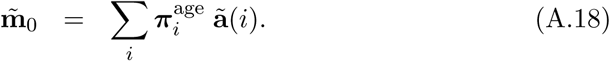

**ñ**(*x*) **= younger sisters of Focal.** Focal has no younger sisters when she is born, so the initial condition is **ñ**_0_ = **0**. Younger sisters are the children of Focal’s mother, so the subsidy term is the reproduction of mothers at age *x* of Focal.

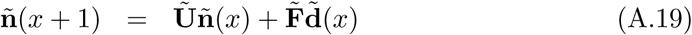

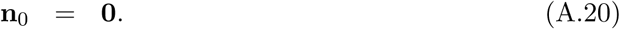

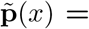 **nieces through older sisters of Focal.** At the birth of Focal, these nieces are the granddaughters of the mother of Focal, so the initial condition is mixture of granddaughters with mixing distribution ***π***^age^. New nieces through older sisters are the result of reproduction by the older sisters of Focal, so the subsidy term applies the fertility matrix to 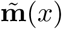.

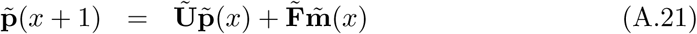

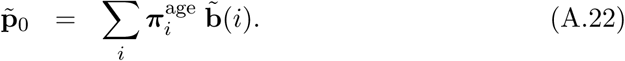

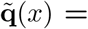 **nieces through younger sisters of Focal.** At the birth of Focal she has no younger sisters, and hence has no nieces through these sisters, so the initial condition is 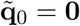. New nieces are the result of reproduction of the younger sisters of Focal, so the subsidy term applies the fertility matrix to **ñ**(*x*).

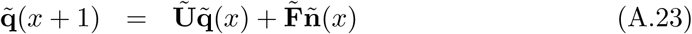

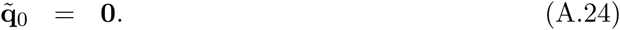

#### A.2.2 Aunts and cousins

Aunts and cousins form another level of side branching on the kinship network; their dynamics follow the same principles as those for sisters and nieces.

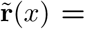 **aunts older than mother of Focal.** These are the older sisters of the mother of Focal. Focal’s mother accumulates no new older sisters, so the subsidy term is ***β***(*x*) = **0**. The initial age×stage structure of these aunts, at the birth of Focal, is a mixture of older sisters, with mixing distribution ***π***^age^

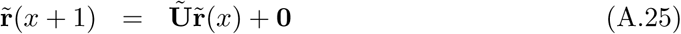

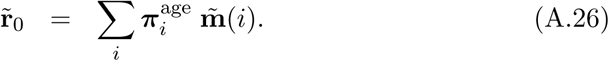

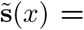 **aunts younger than mother of Focal.** These aunts are the younger sisters of the mother of Focal. They are the children of the grandmother of Focal, so the subsidy term applies the fertility matrix to the vector 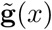 of the grandmothers of Focal. The initial age distribution of these aunts, at the birth of Focal, is a mixture of the populations **ñ** of younger sisters, with mixing distribution ***π***^age^.

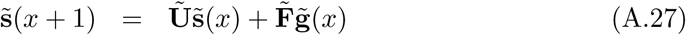

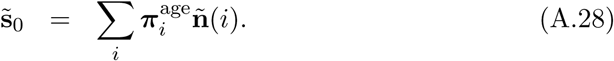

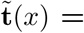 **cousins from aunts older than mother of Focal.** These cousins are the nieces of the mother of Focal through her older sisters. The subsidy term applies the fertility matrix to the population of these older sisters. The initial condition is a mixture of the population vectors T of nieces through older sisters, with mixing distribution ***π***^age^.

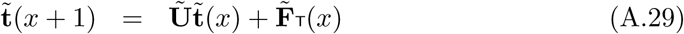

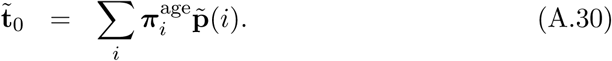

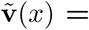 **cousins from aunts younger than mother of Focal.** These cousins are the nieces of the mother of Focal through her younger sisters. The subsidy term applies the fertility matrix to the population of these younger sisters. The initial condition is a mixture of the age distributions of nieces through younger sisters, with mixing distribution ***π***^age^.

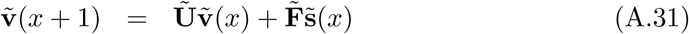

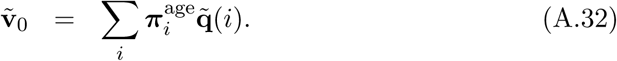

## Footnotes

1 Models that incorporate three or more characteristics have been called hyperstate models (Roth and Caswell, 2016). They generalize the construction to be developed here, and have potential applications to kinship models as well, but we do not consider them further in this study.

2 The joke about the physicist, the engineer, and the mathematician confronting a fire in a wastebasket is left as an exercise for the reader.

3 The HFD cautions that the data for Slovakia before 1960 are not reliable, so we chose to examine only 1960–2014.

4 For completists or the curious, figures for all types of kin are presented in Online Supplementary Material.

5 See websites https://draxe.com/health/benefits-of-aunts/ and http://melanienotkin.com/portfolios/pank-power/

6 The author cringes at the necessity of doing this.

7 Lotka (1939) expressed it well: “The conditions that present themselves in an actual population are always excessively complicated. Whoever has failed to grasp clearly the necessary relations among the characteristics of a theoretical population subject to simple hypotheses, will certainly be unable to manage in the much more complicated relations that exist in a real population.”

## References

Barclay, K. and Kolk, M. (2019). Parity and mortality: an examination of different explanatory mechanisms using data on biological and adoptive parents. European Journal of Population 35(1): 63–85.

Blake, J. (1989). Family Size and Achievement. Berkeley, California: University of California Press.

Burch, T.K. (2018). Model-based demography: Essays on integrating data, technique and theory. Cham, Switzerland: Springer Nature.

Caswell, H. (2001). Matrix Population Models: Construction, Analysis, and Interpretation. Sunderland, MA: Sinauer Associates, 2nd ed.

Caswell, H. (2014). A matrix approach to the statistics of longevity in heterogeneous frailty models. Demographic Research 31: 553–592.

Caswell, H. (2019a). The formal demography of kinship: A matrix formulation. Demographic Research 41: 679–712.

Caswell, H. (2019b). Inequality and variance in longevity. Unpublished.

Caswell, H., de Vries, C., Hartemink, N., Roth, G., and van Daalen, S.F. (2018). Age×stageclassified demographic analysis: a comprehensive approach. Ecological Monographs 88(4): 560–584.

Caswell, H. and Vindenes, Y. (2018). Demographic variance in heterogeneous populations: Matrix models and sensitivity analysis. Oikos 127: 648–663.

Cools, S., Markussen, S., and Strøm, M. (2017). Children and careers: How family size affects parents’ labor market outcomes in the long run. Demography 54(5): 1773–1793.

de Vries, C. and Caswell, H. (2019). Stage-structured evolutionary demography: Linking life histories, population genetics, and ecological dynamics. The American Naturalist 193(4): 545–559.

Downey, D.B. (2001). Number of siblings and intellectual development: The resource dilution explanation. American Psychologist 56(6-7): 497–504.

Fahey, T. (2017). The sibsize revolution and social disparities in children’s family contexts in the United States, 1940–2012. Demography 54(3): 813–834.

Goodman, L.A., Keyfitz, N., and Pullum, T.W. (1974). Family formation and the frequency of various kinship relationships. Theoretical Population Biology 5(1): 1–27.

Hammel, E. (2005). Demographic dynamics and kinship in anthropological populations. Proceedings of the National Academy of Sciences USA 102(6): 2248–2253.

Hartemink, N., Missov, T.I., and Caswell, H. (2017). Stochasticity, heterogeneity, and variance in longevity in human populations. Theoretical Population Biology 114: 107–117.

Hartemink, N. and Caswell, H. (2018). Variance in animal longevity: contributions of heterogeneity and stochasticity. Population Ecology 60: 89–99.

Henderson, H.V. and Searle, S.R. (1981). The vec-permutation matrix, the vec operator and Kronecker products: a review. Linear and Multilinear Algebra 9: 271–288.

Hernández, C.M., van Daalen, S.F., Caswell, H., Neubert, M.G., and Gribble, K.E. (2019). Maternal effect senescence and fitness: a demographic analysis of a novel model organism. bioRxiv doi:http://dx.doi.org/10.1101/847640.

Hrdy, S.B. (2009). Mothers and Others. Cambridge, Massachusetts: Harvard University Press.

Human Fertility Database (2019). Max Planck Institute for Demographic Research (Germany) and the Vienna Institute of Demography (Austria). www.humanfertility.org.

Human Mortality Database (2019). University of California, Berkeley (USA), and Max Planck Institute for Demographic Research (Germany). www.mortality.org URL http://www.mortality.org.

Hunter, C.M. and Caswell, H. (2005). The use of the vec-permutation matrix in spatial matrix population models. Ecological Modelling 188(1): 15–21.

Jasilioniene, A., Jdanov, D.A., Sobotka, T., Andreev, E.M., Zeman, K., Shkolnikov, V.M., Goldstein, J., Nash, E.J., Philipov, D., and Rodriguez, G. (2019). Methods protocol for the Human Fertility Database. Tech. rep., Max Planck Institute for Demographic Research (Germany) and the Vienna Institute of Demography (Austria).

Jenouvrier, S., Aubry, L.M., Barbraud, C., Weimerskirch, H., and Caswell, H. (2018). Interacting effects of unobserved heterogeneity and individual stochasticity in the life cycle of the Southern fulmar. Journal of Animal Ecology 87: 212–222.

Kalmijn, M. and van de Werfhorst, H.G. (2016). Sibship size and gendered resource dilution in different societal contexts. PLOS ONE 11(8): e0160953.

Keyfitz, N. and Caswell, H. (2005). Applied Mathematical Demography. New York, New York: Springer, 3rd ed.

Klepac, P. and Caswell, H. (2011). The stage-structured epidemic: linking disease and demography with a multi-state matrix approach model. Theoretical Ecology 4(3): 301–319.

Kramer, K.L. (2019). How there got to be so many of us: the evolutionary story of population growth and a life history of cooperation. Journal of Anthropological Research 75(4): 472–497.

Kravdal, Ø., Glattre, E., and Haldorsen, T. (1991). Positive correlation between parity and incidence of thyroid cancer: new evidence based on complete Norwegian birth cohorts. International Journal of Cancer 49(6): 831–836.

Lotka, A.J. (1939). Théorie analytique des associations biologiques: analyse démographique avec application particulière ‘a l’espèce humaine. Actualités Scientifiques et Industrielles. Paris, France: Hermann et Cie, (Published in translation as: Analytical theory of biological populations, translated by D.P. Smith and H. Rossert. Plenum Press, New York, 1998) ed.

Mills, M. (2000). The transformation of partnerships. Canada, the Netherlands, and the Russian Federation in the age of modernity. Amsterdam, The Netherlands: Thela Thesis Population Studies Series.

Nitsch, A., Faurie, C., and Lummaa, V. (2014). Alloparenting in humans: Fitness consequences of aunts and uncles on survival in historical Finland. Behavioral Ecology 25(2): 424–433.

Plesko, I., Dimitrova, E., Somogyi, J., Preston-Martin, S., Day, N., and Tzonou, A. (1985). Parity and cancer risk in Slovakia. International Journal of Cancer 36(5): 529–533.

Pollard, J.H. (1973). Mathematical Models for the Growth of Human Populations. Cambridge, UK: Cambridge University Press.

Preston, S.H. (1976). Family sizes of children and family sizes of women. Demography 13(1): 105–114.

Pullum, T.W. and Wolf, D.A. (1991). Correlations between frequencies of kin. Demography 28(3): 391–409.

Roth, G. and Caswell, H. (2016). Hyperstate matrix models: extending demographic state spaces to higher dimensions. Methods in Ecology and Evolution 7(12): 1438–1450.

Schoen, R. (2016). Hierarchical multistate models from population data: an application to parity statuses. PeerJ 4: e2535.

Schoen, R. (2019a). On the implications of age-specific fertility for sibships and birth spacing. In: Schoen, R. (ed.). Analytical Family Demography. Cham, Switzerland: Springer Nature: 201–214.

Schoen, R. (2019b). Parity progression and the kinship network. In: Schoen, R. (ed.). Analytical Family Demography. Cham, Switzerland: Springer Nature: 189–199.

Sear, R. and Mace, R. (2008). Who keeps children alive? a review of the effects of kin on child survival. Evolution and Human Behavior 29(1): 1–18.

Song, X. and Mare, R.D. (2019). Shared lifetimes, multigenerational exposure, and educational mobility. Demography 56(3): 891–916.

Sonneveldt, E., Plosky, W.D., and Stover, J. (2013). Linking high parity and maternal and child mortality: What is the impact of lower health services coverage among higher order births? BMC Public Health 13(3): S7.

Tanskanen, A.O. and Danielsbacka, M. (2019). Intergenerational Family Relations: An Evolutionary Social Science Approach. Routledge Advances in Sociology. New York, New York, USA: Routledge.

Valtorta, N.K., Kanaan, M., Gilbody, S., Ronzi, S., and Hanratty, B. (2016). Loneliness and social isolation as risk factors for coronary heart disease and stroke: systematic review and meta-analysis of longitudinal observational studies. Heart 102(13): 1009–1016.

van Daalen, S.F. and Caswell, H. (2020). Variance as a life history outcome: Sensitivity analysis of the contributions of stochasticity and heterogeneity. Ecological Modelling 147: https://doi.org/10.1016/j.ecolmodel.2019.108856.

van den Broek, T., Tosi, M., and Grundy, E. (2019). Offspring and later-life loneliness in Eastern and Western Europe. Journal of Family Research 31(2): 199–215.

Wachter, K.W. (1997). Kinship resources for the elderly. Philosophical Transactions of the Royal Society of London. Series B: Biological Sciences 352(1363): 1811–1817.

Willekens, F., van Imhoff, E., and Wright, J.D. (2015). Formal demography of families and households. International Encyclopedia of the Social & Behavioral SciencesInternational Encyclopedia of the Social & Behavioral Sciences 8(2): 725–730.

Zhao, M. and Zhang, Y. (2019). Parental childcare support, sibship status, and mothers’ second-child plans in urban China. Demographic Research 41: 1315–1346.

