## Supplemental figures for "The formal demography of kinship II: Multistate models, parity, and sibship"

Hal Caswell
Institute for Biodiversity and Ecosystem Dynamics
PO Box 94240
1090 GE Amsterdam
The Netherlands

Revised March 2, 2020

This supplementary document contains a complete set of kinship outputs for the age×parity model for Slovakia, used as an example in the main text. The different relatives are presented in the same order in each section.

#### Contents

**1 Parity distribution of Focal** **1**

**2 Age structures of kin** **2**

**3 Marginal parity structure of kin over time** **12**

**4 Marginal parity structure of kin by age of Focal** **17**

**5 Marginal parity distributions by age of Focal** **22**

**6 Marginal parity distribution over time** **27**

**7 Prevalence of low parity kin over time** **32**

### 1 Parity distribution of Focal

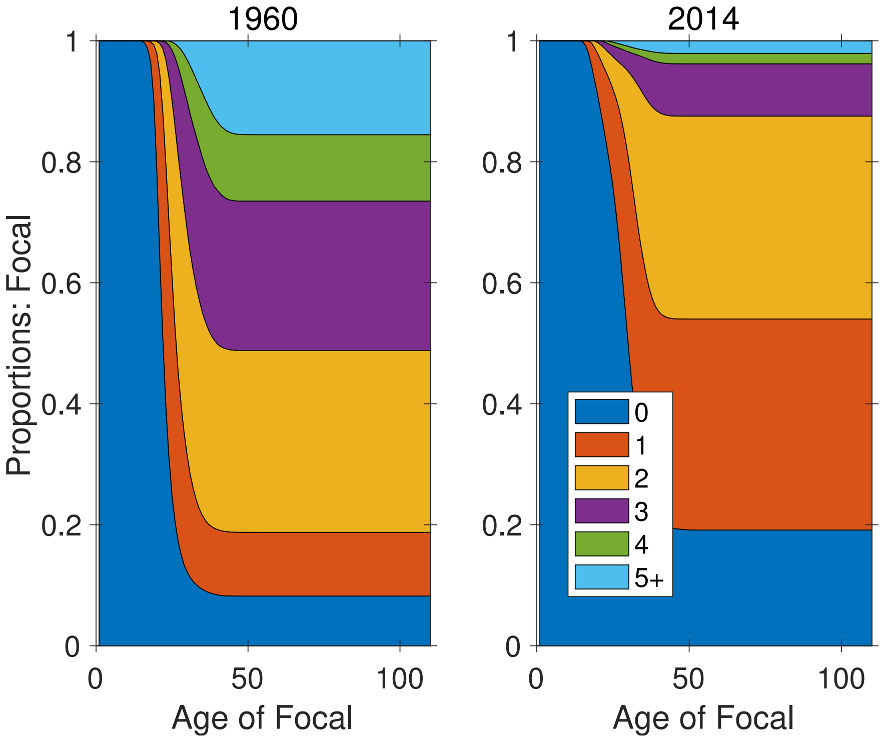


**Figure S-1:** Expected parity distribution of Focal as a function of age, in 1960 and 2014.

#### 2 Age structures of kin

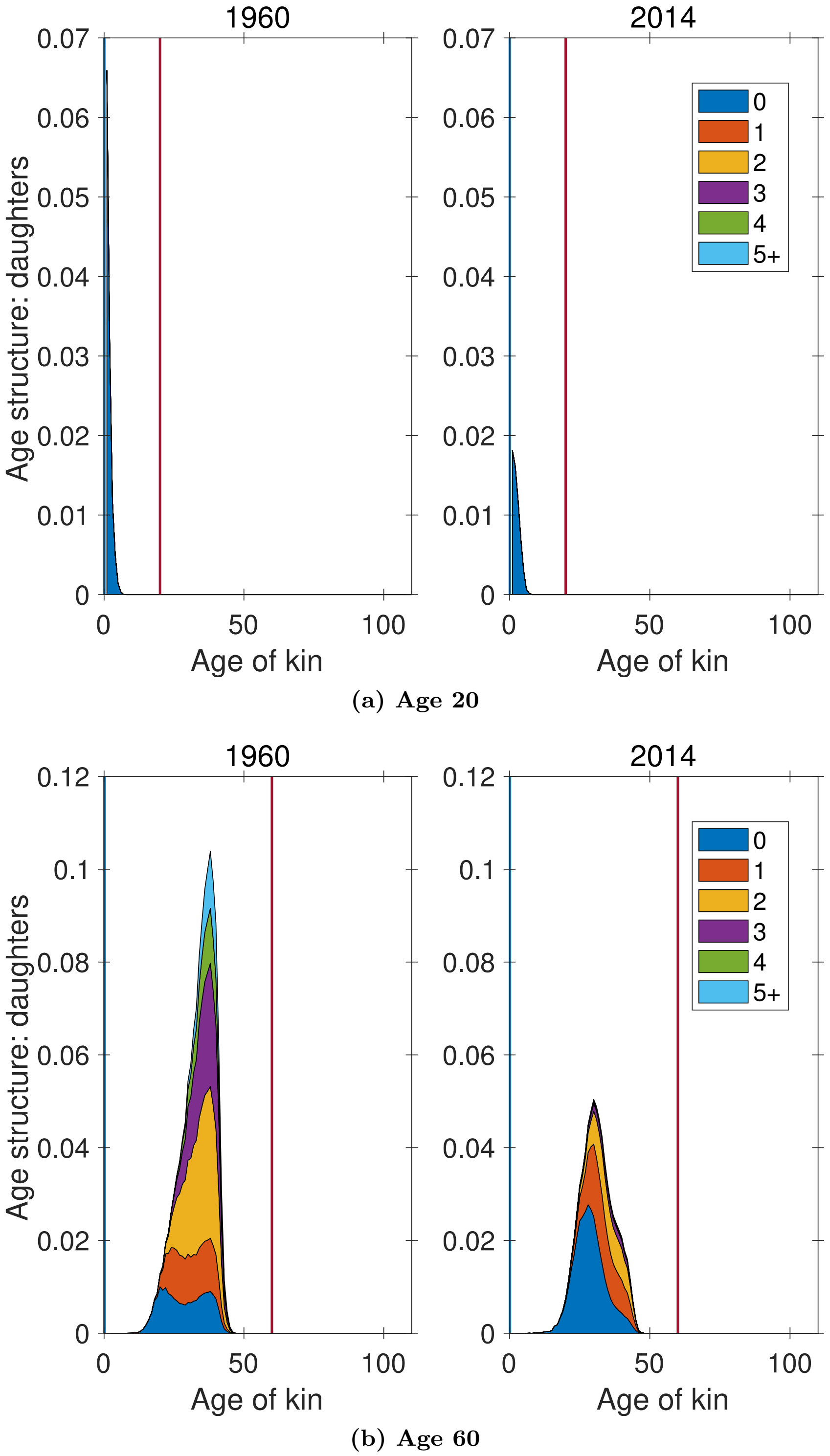


**Figure S-2:** Age×parity structure of daughters, at ages 20 and 60 of Focal. Vertical line indicates age of Focal for reference.

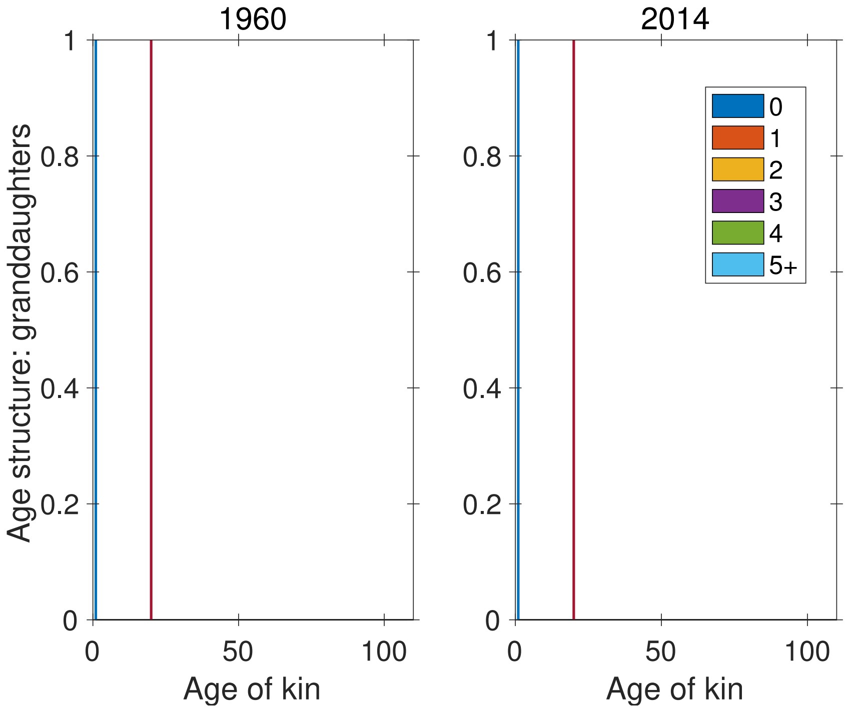


**(a) Age 20**

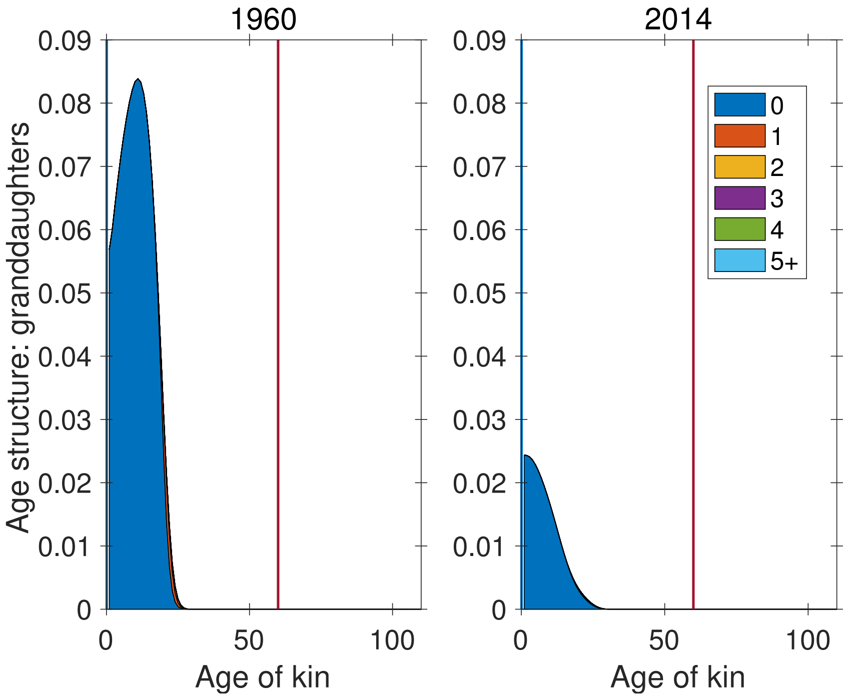


**(b) Age 60**

**Figure S-3:** Age×parity structure of granddaughters, at ages 20 and 60 of Focal. Vertical line indicates age of Focal for reference.

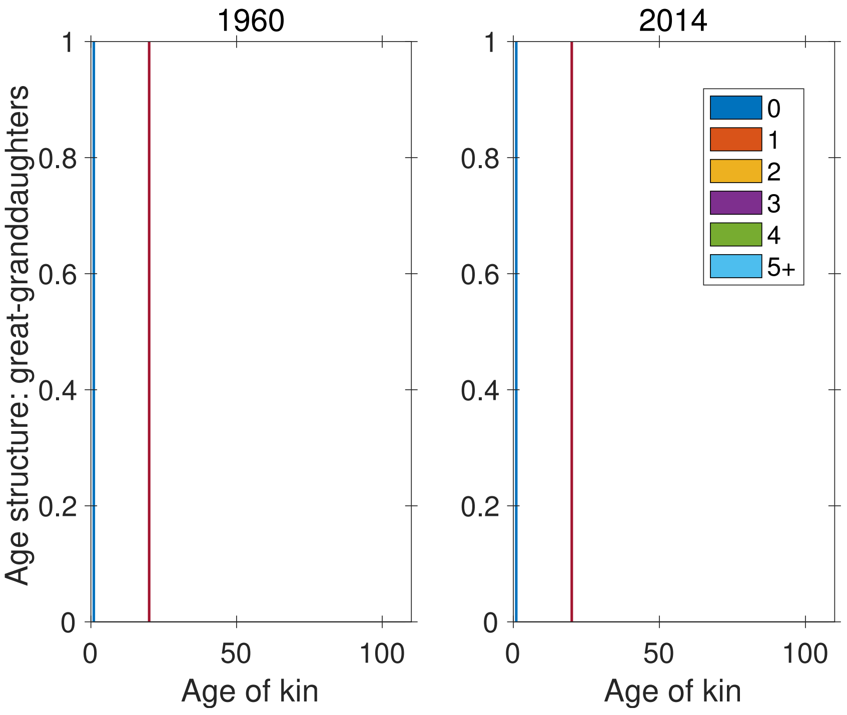


(a) **Age 20**

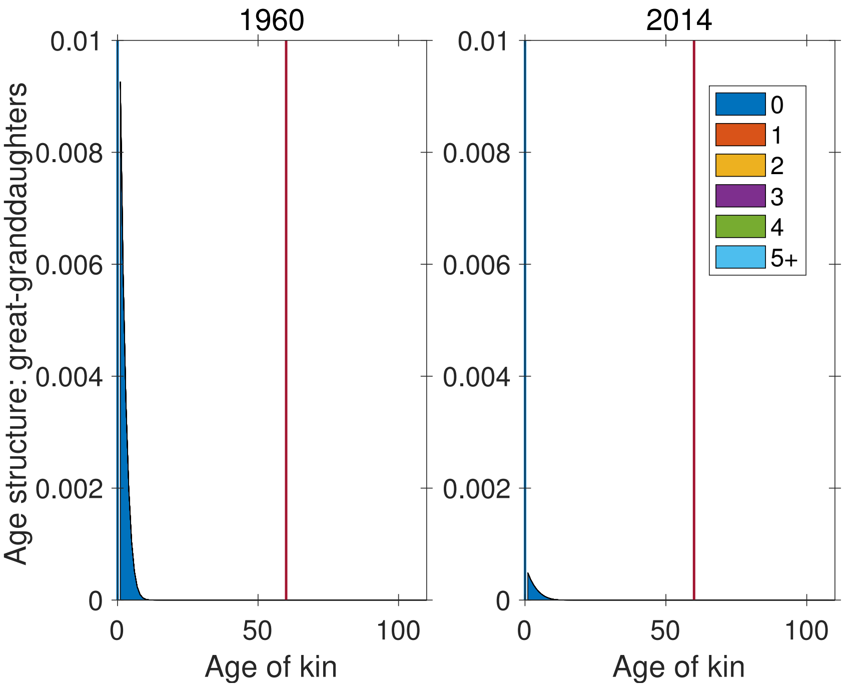


(b) **Age 60**

**Figure S-4:** Age×parity structure of great-granddaughters, at ages 20 and 60 of Focal. Vertical line indicates age of Focal for reference.

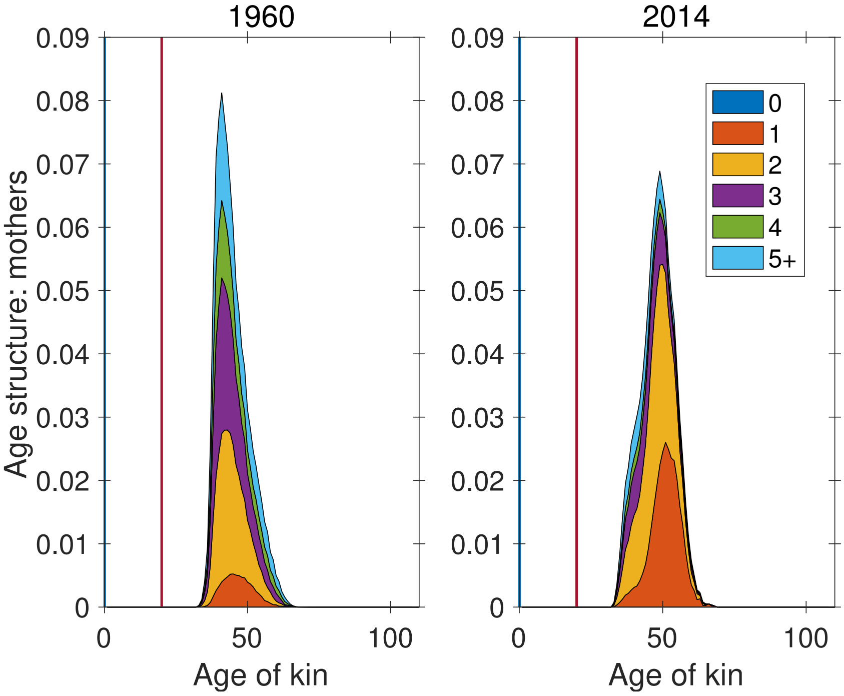


**(a) Age 20**

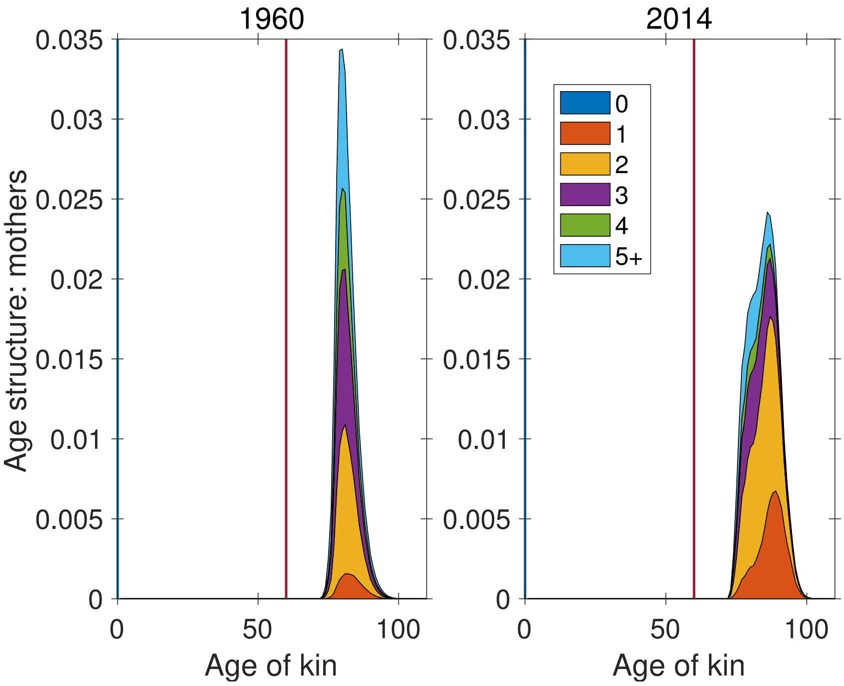


**(b) Age 60**

**Figure S-5:** Age×parity structure of mothers, at ages 20 and 60 of Focal. Vertical line indicates age of Focal for reference.

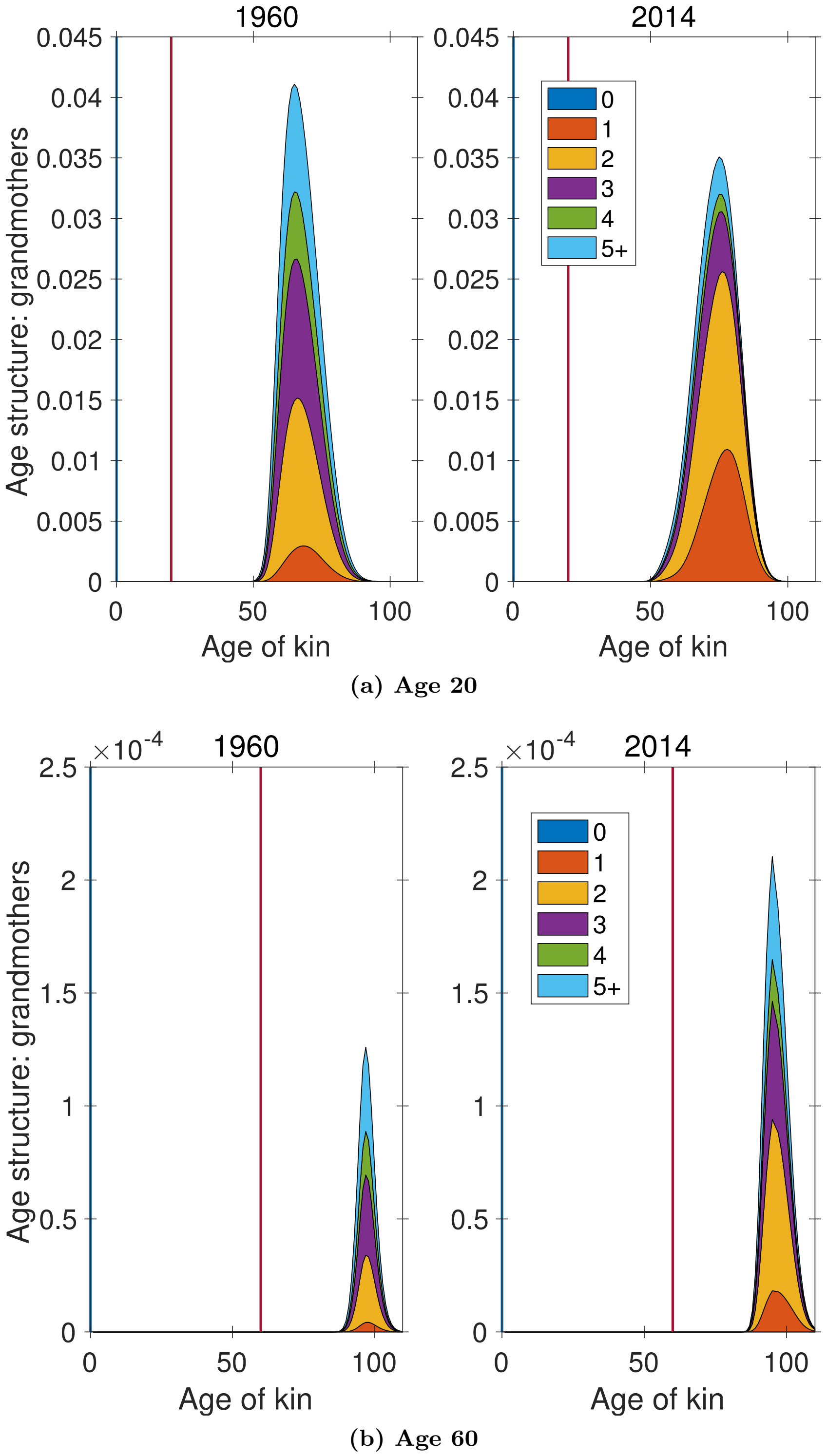


**Figure S-6:** Age×parity structure of grandmothers, at ages 20 and 60 of Focal. Vertical line indicates age of Focal for reference.

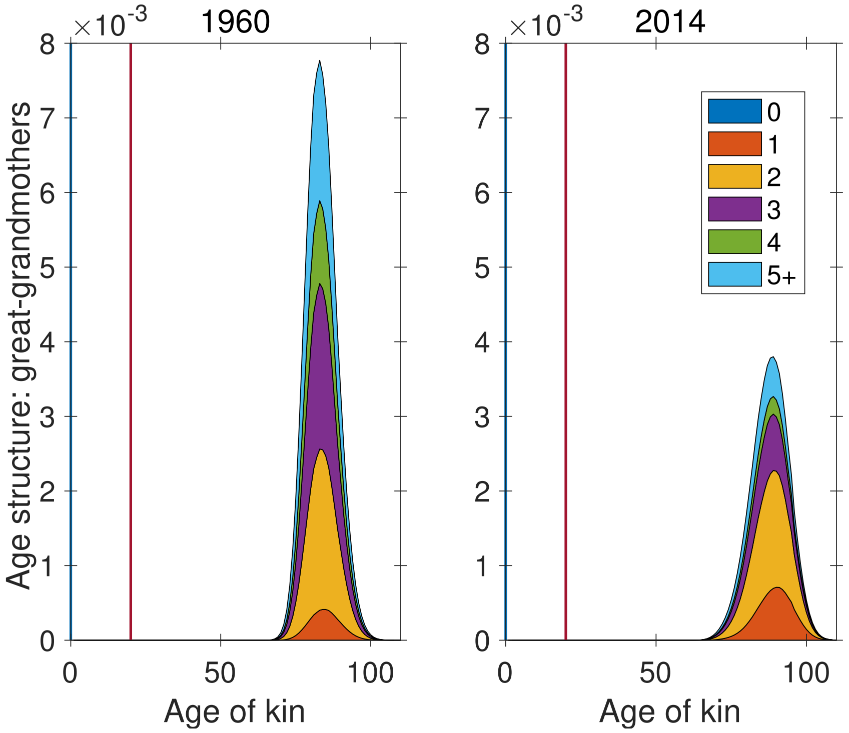


**(a) Age 20**

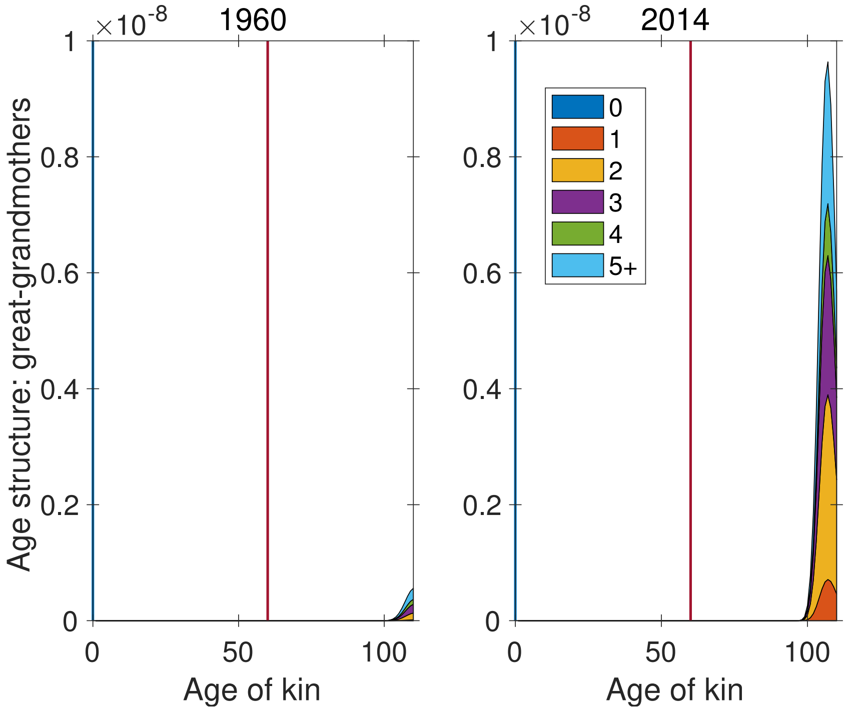


**(b) Age 60**

**Figure S-7:** Age×parity structure of great-grandmothers, at ages 20 and 60 of Focal. Vertical line indicates age of Focal for reference.

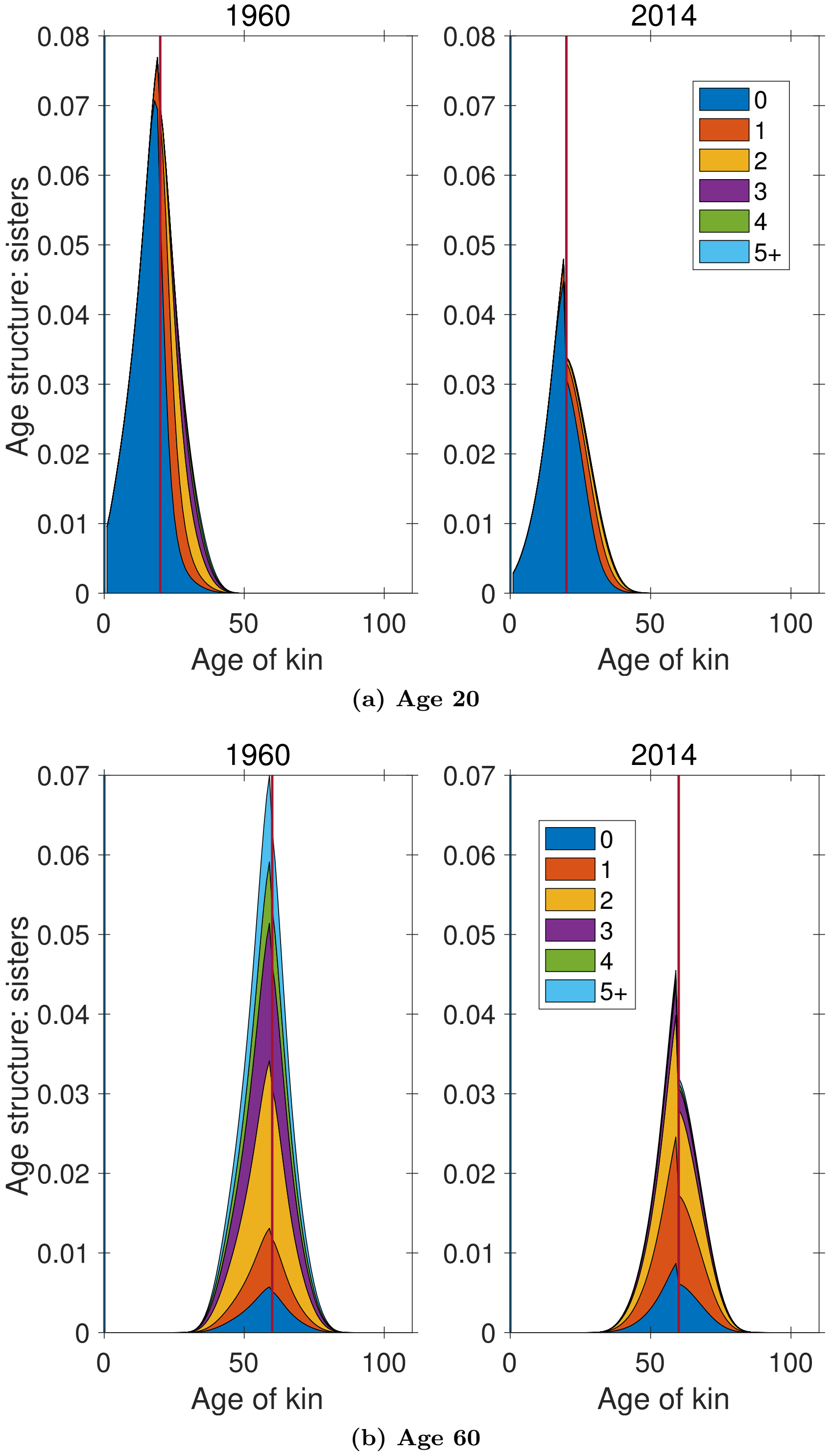


**Figure S-8:** Age×parity structure of sisters, at ages 20 and 60 of Focal. Vertical line indicates age of Focal for reference.

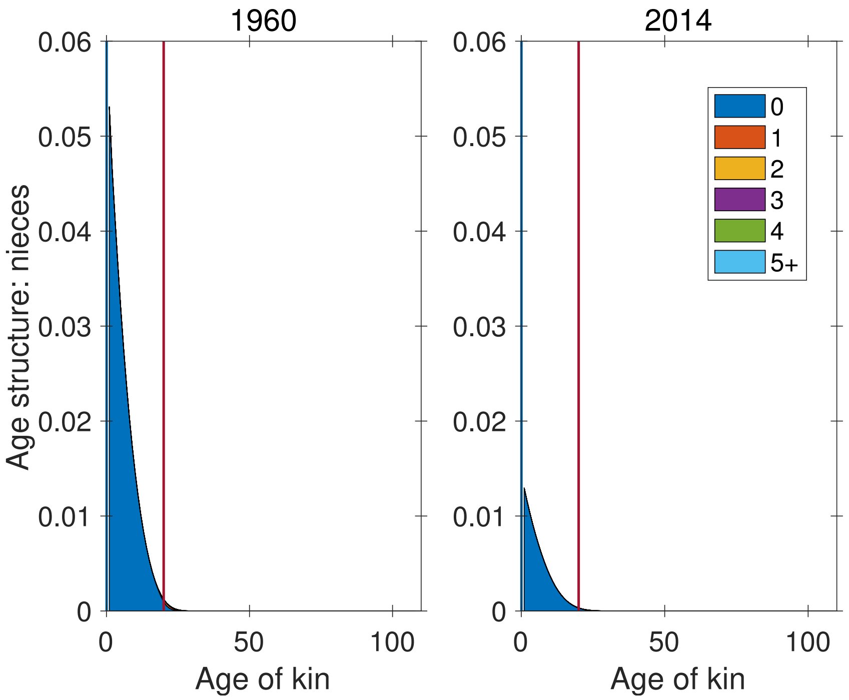


**(a) Age 20**

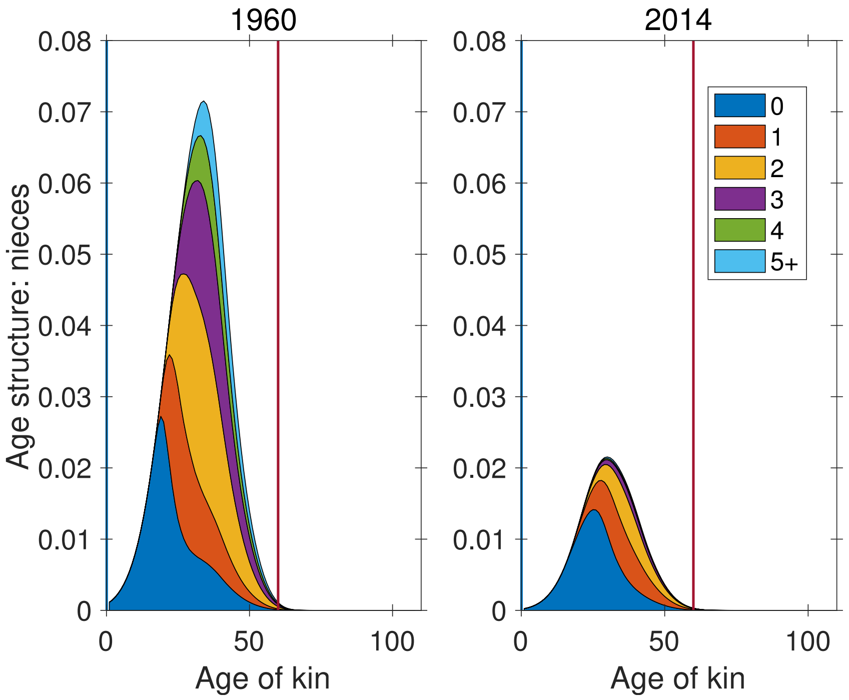


**(b) Age 60**

**Figure S-9:** Age×parity structure of nieces, at ages 20 and 60 of Focal. Vertical line indicates age of Focal for reference.

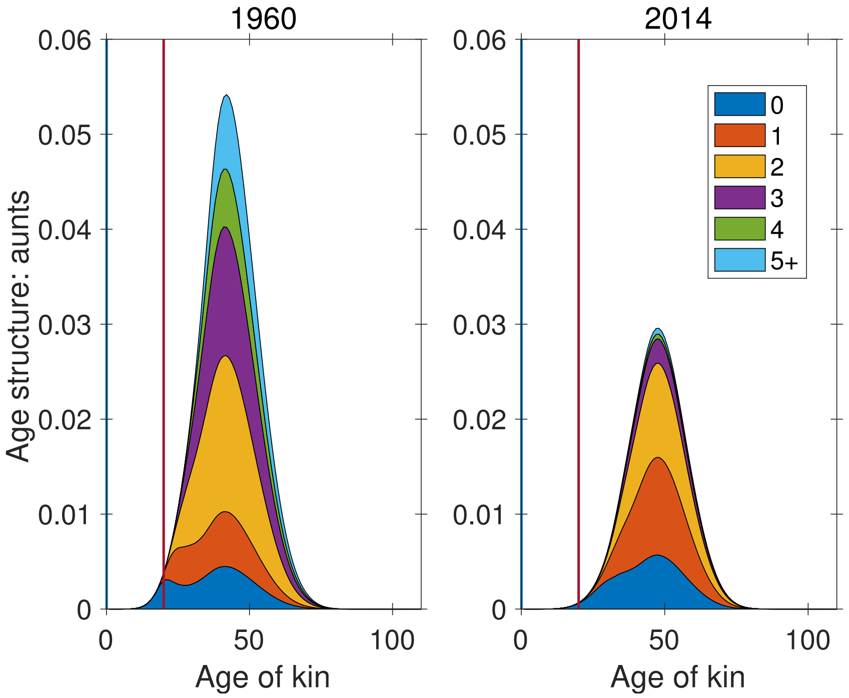


**(a) Age 20**

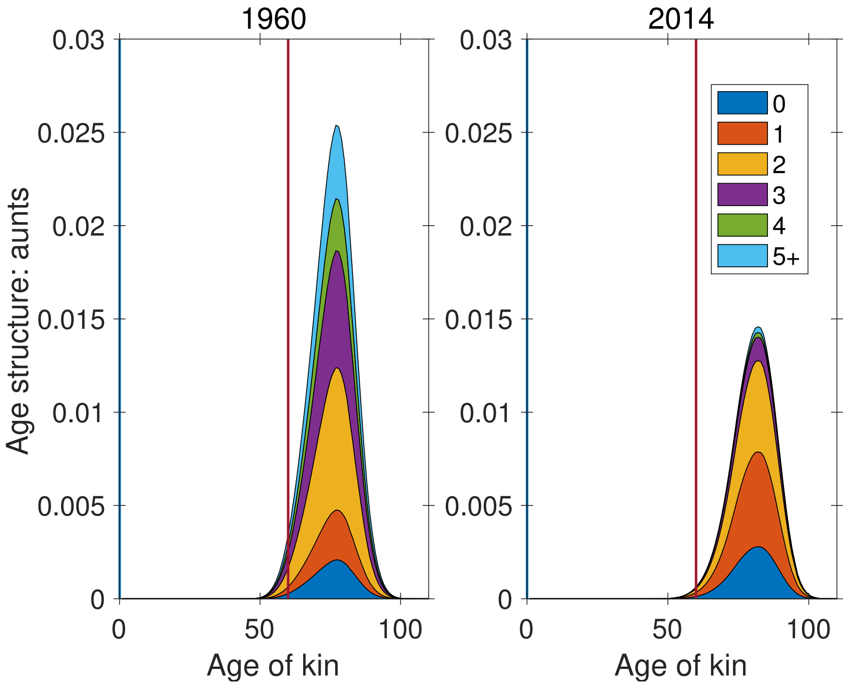


**(b) Age 60**

**Figure S-10:** Age×parity structure of aunts, at ages 20 and 60 of Focal. Vertical line indicates age of Focal for reference.

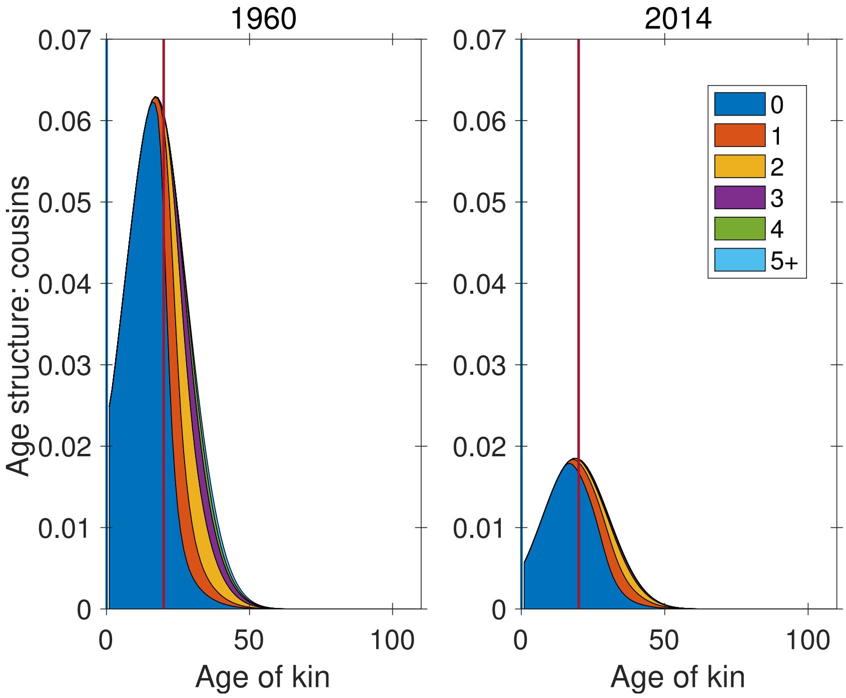


**(a) Age 20**

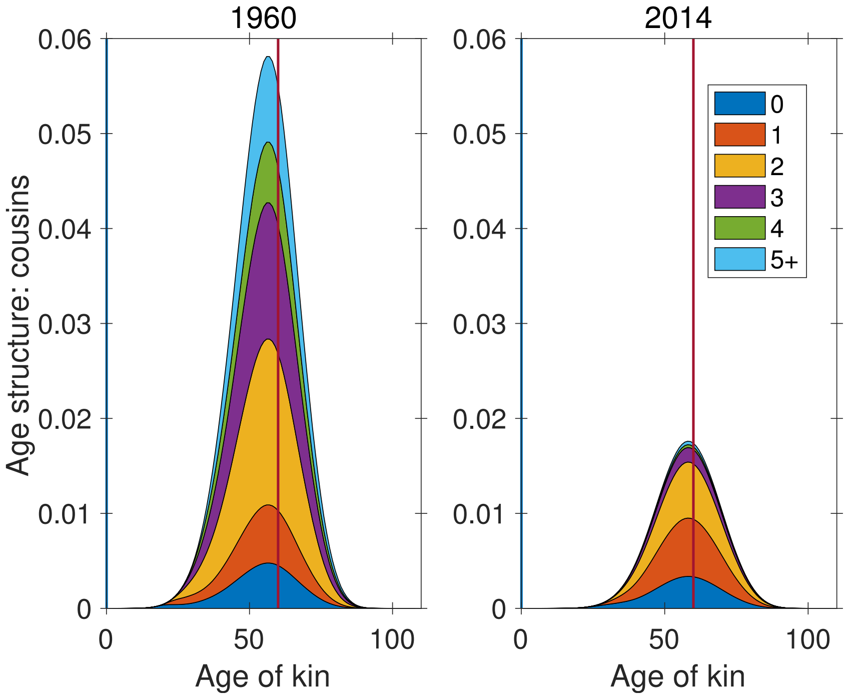


**(b) Age 60**

**Figure S-11:** Age×parity structure of cousins, at ages 20 and 60 of Focal. Vertical line indicates age of Focal for reference.

#### 3 Marginal parity structure of kin over time

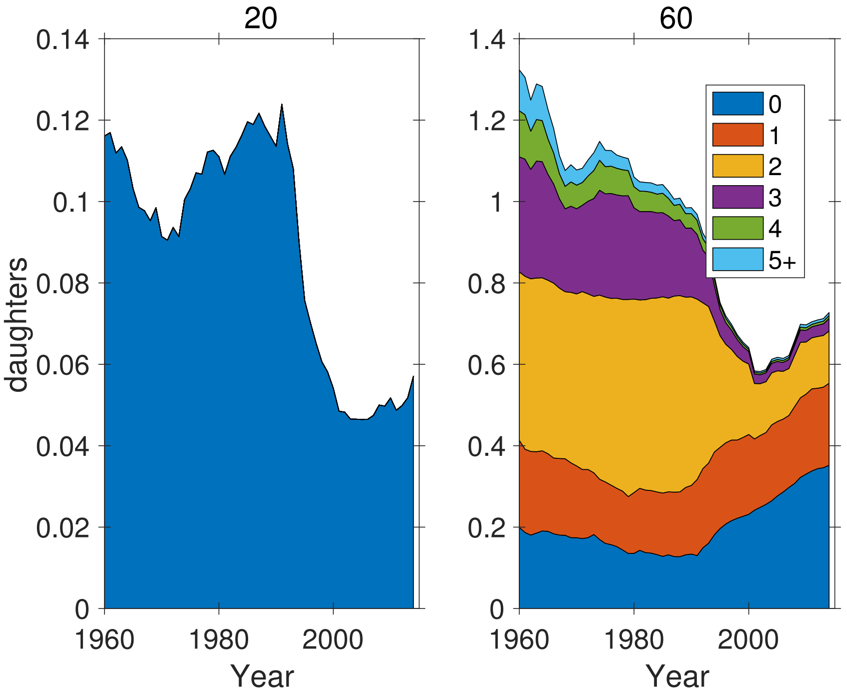


**Figure S-12:** Marginal parity structure of daughters as a function of time, for ages 20 and 60 of Focal. Note different ordinate scales.

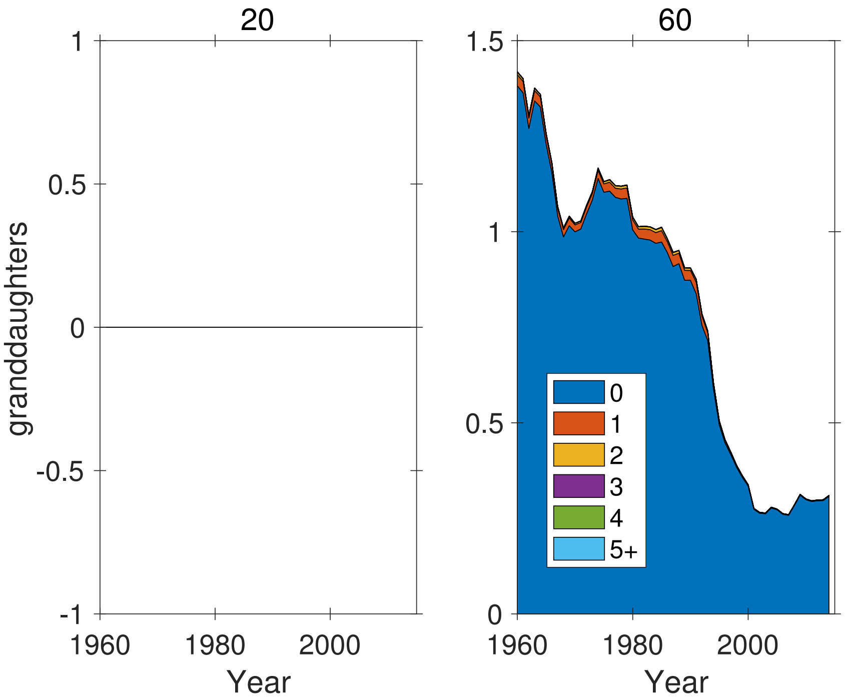


**Figure S-13:** Marginal parity structure of granddaughters as a function of time, for ages 20 and 60 of Focal. Note different ordinate scales.

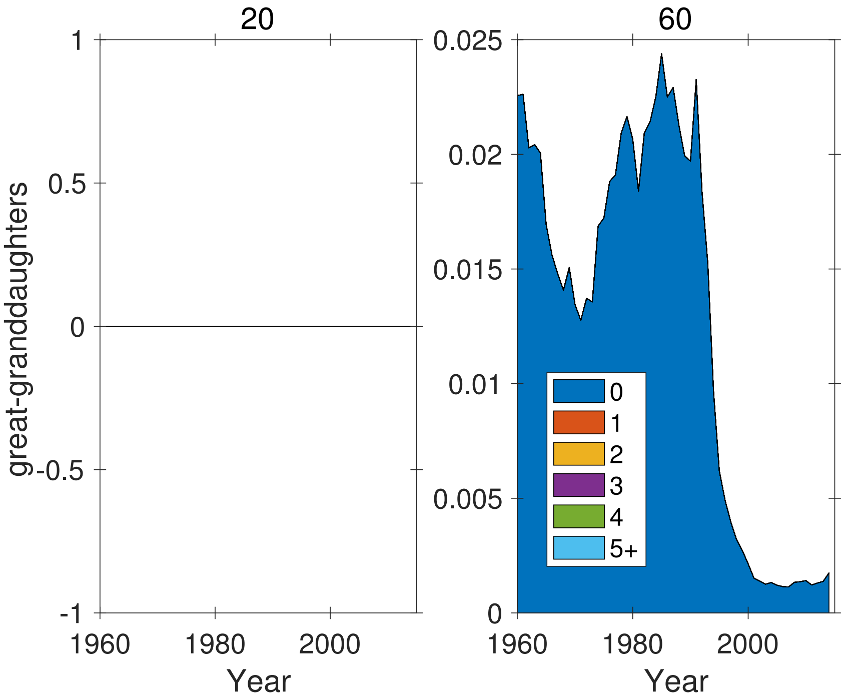


**Figure S-14:** Marginal parity structure of great-granddaughters as a function of time, for ages 20 and 60 of Focal. Note different ordinate scales.

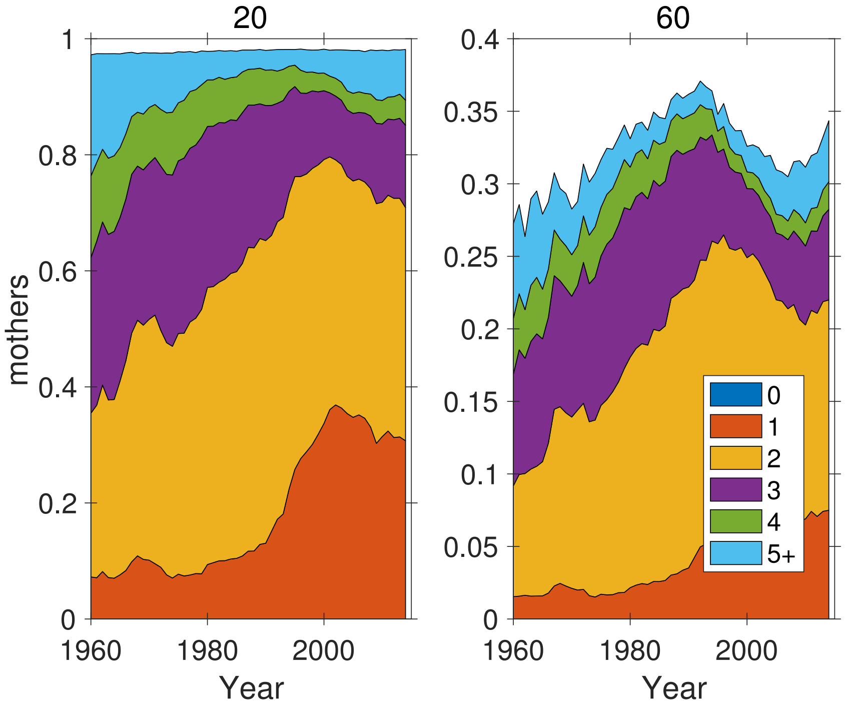


**Figure S-15:** Marginal parity structure of mothers as a function of time, for ages 20 and 60 of Focal. Note different ordinate scales.

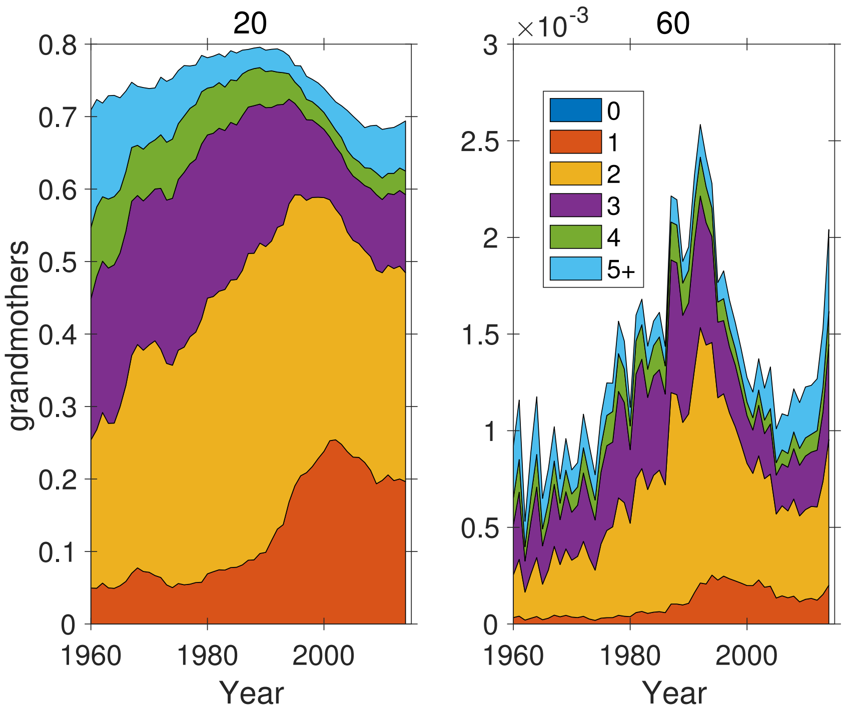


**Figure S-16:** Marginal parity structure of grandmothers as a function of time, for ages 20 and 60 of Focal. Note different ordinate scales.

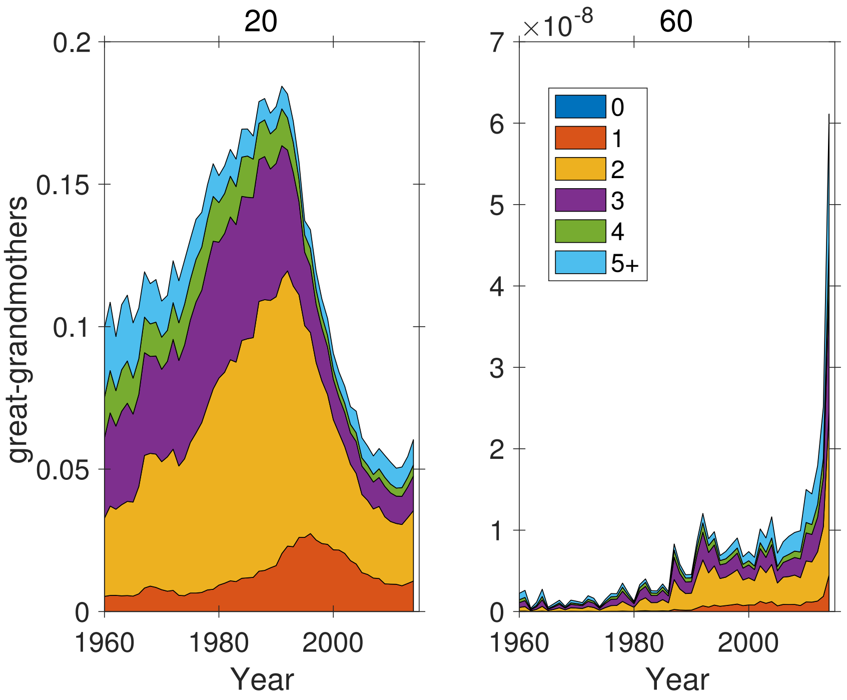


**Figure S-17:** Marginal parity structure of great-grandmothers as a function of time, for ages 20 and 60 of Focal. Note different ordinate scales.

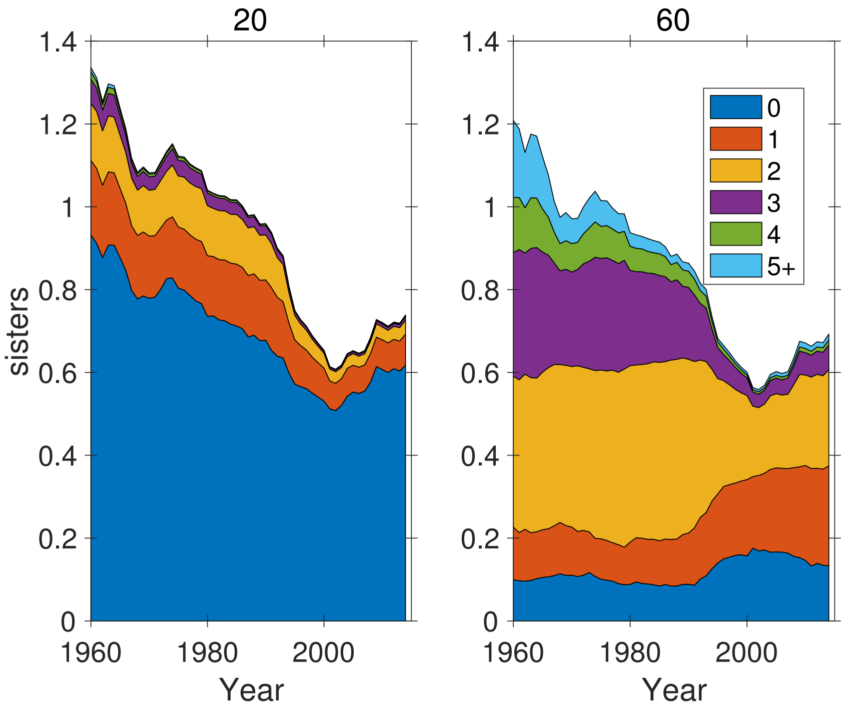


**Figure S-18:** Marginal parity structure of sisters as a function of time, for ages 20 and 60 of Focal. Note different ordinate scales.

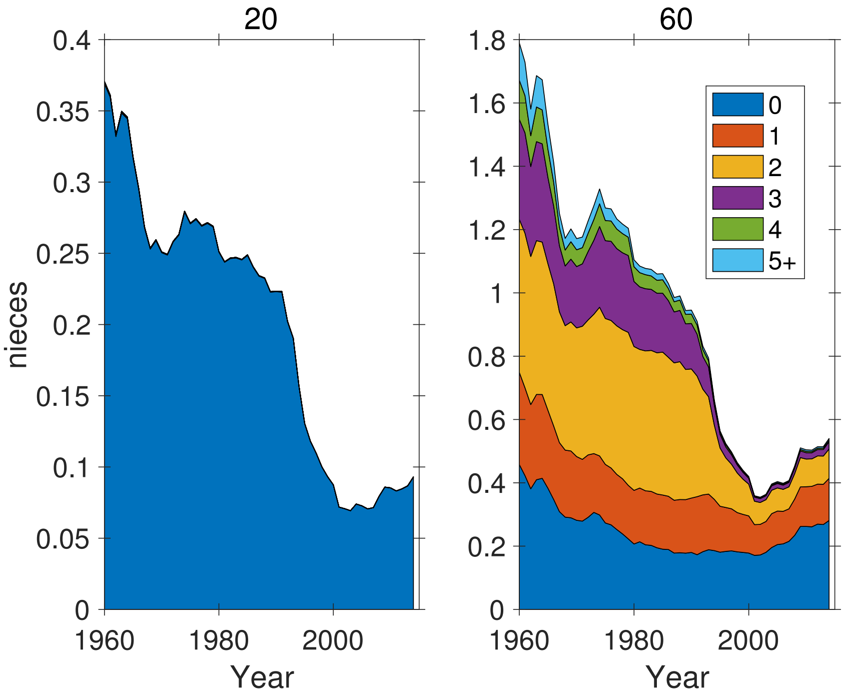


**Figure S-19:** Marginal parity structure of nieces as a function of time, for ages 20 and 60 of Focal. Note different ordinate scales.

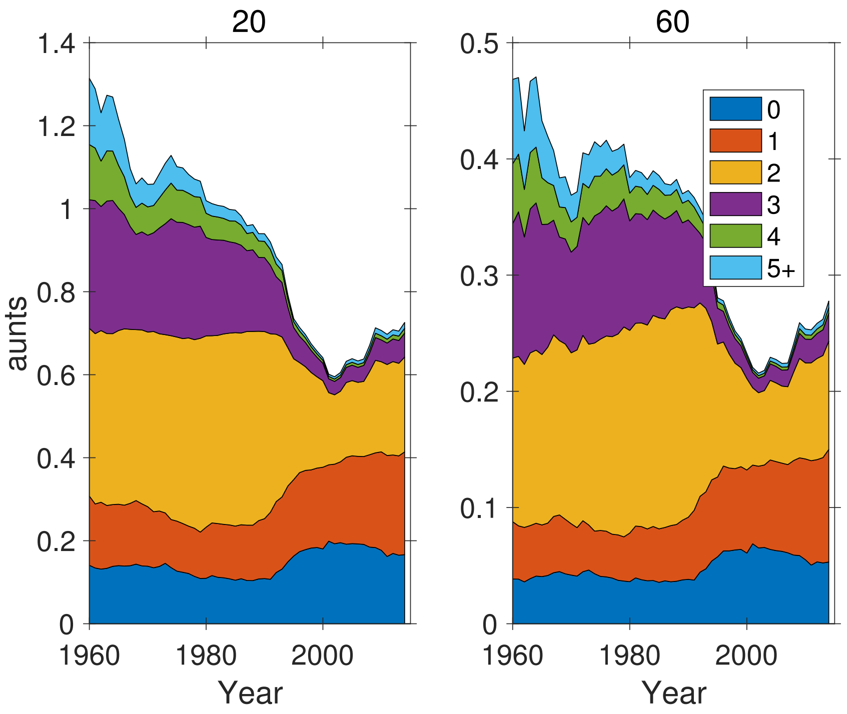


**Figure S-20:** Marginal parity structure of aunts as a function of time, for ages 20 and 60 of Focal. Note different ordinate scales.

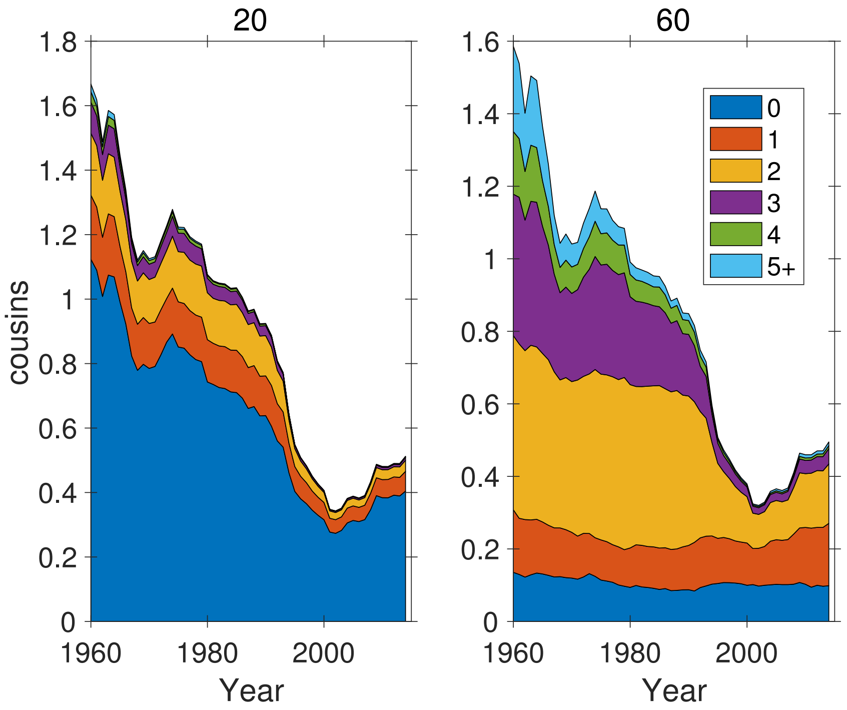


**Figure S-21:** Marginal parity structure of cousins as a function of time, for ages 20 and 60 of Focal. Note different ordinate scales.

### 4 Marginal parity structure of kin by age of Focal

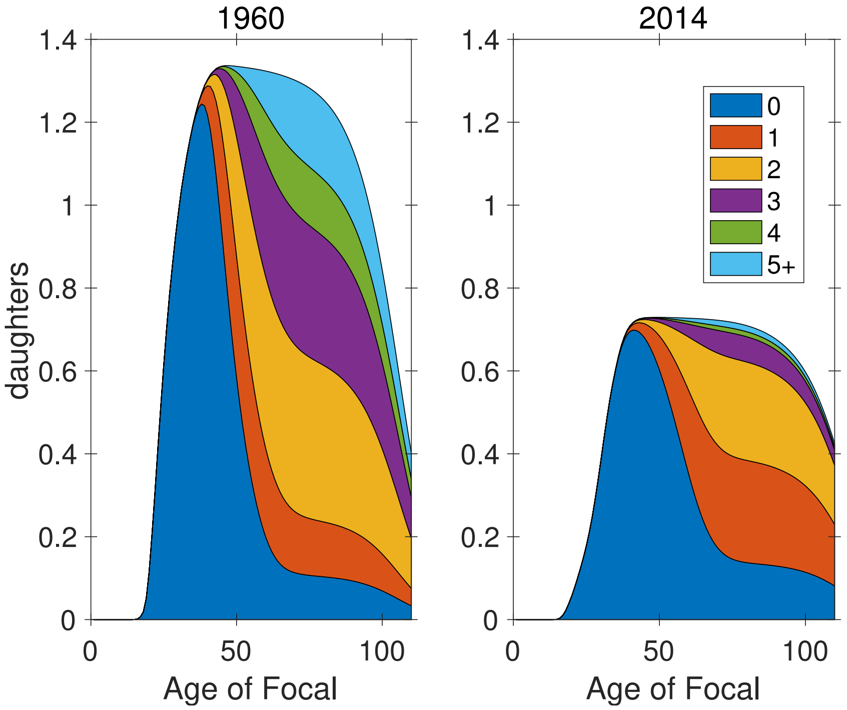


**Figure S-22:** Marginal parity structure of daughters as a function of the age of Focal.

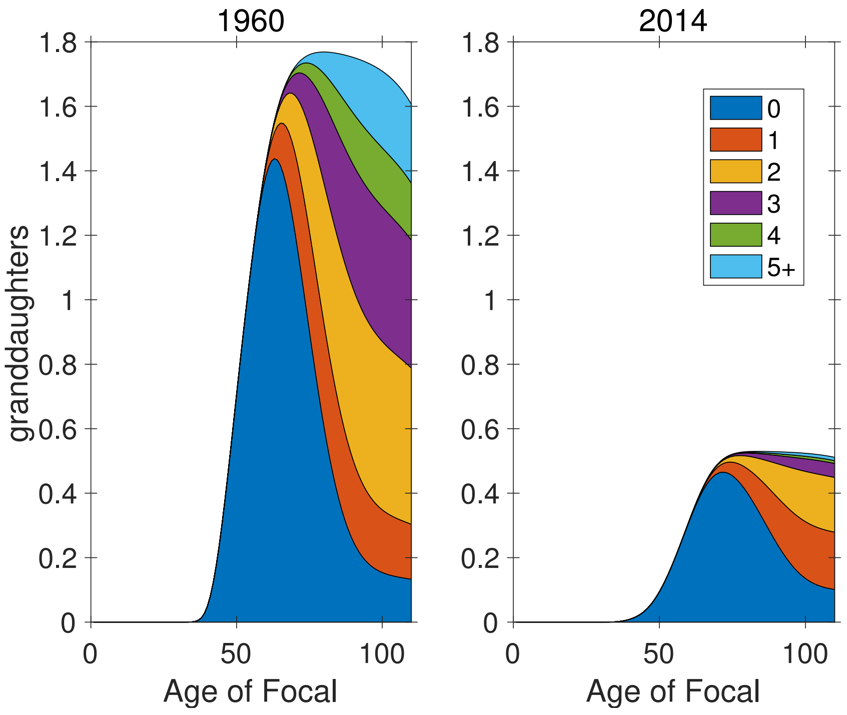


**Figure S-23:** Marginal parity structure of granddaughters as a function of the age of Focal.

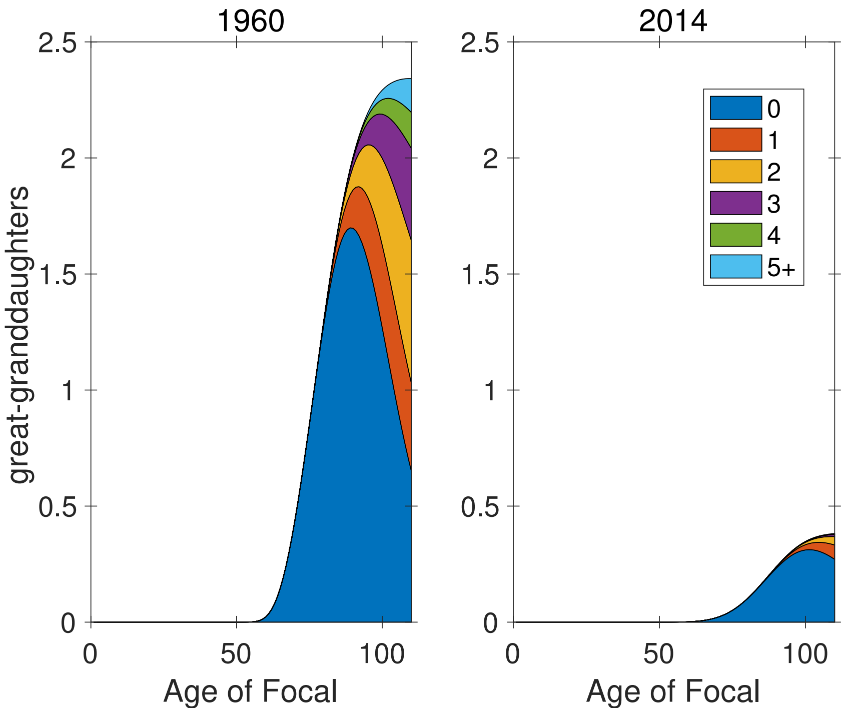


**Figure S-24:** Marginal parity structure of great-granddaughters as a function of the age of Focal.

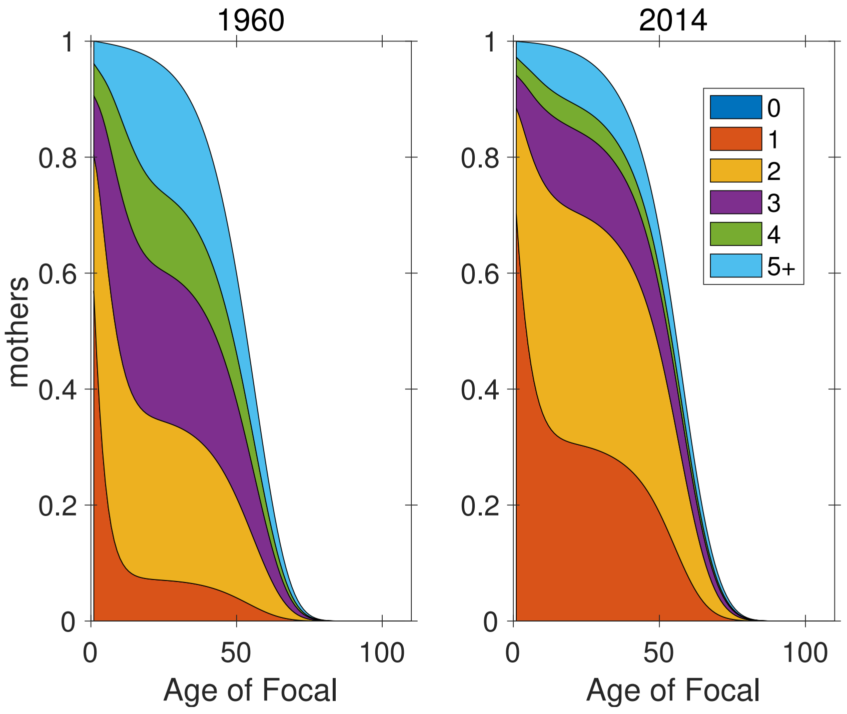


**Figure S-25:** Marginal parity structure of mothers as a function of the age of Focal.

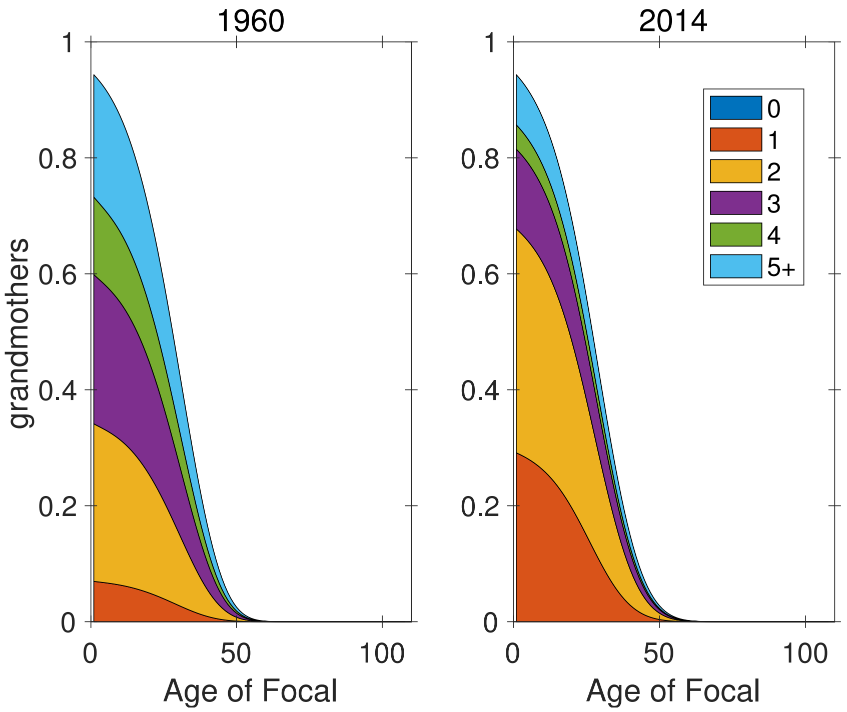


**Figure S-26:** Marginal parity structure of grandmothers as a function of the age of Focal.

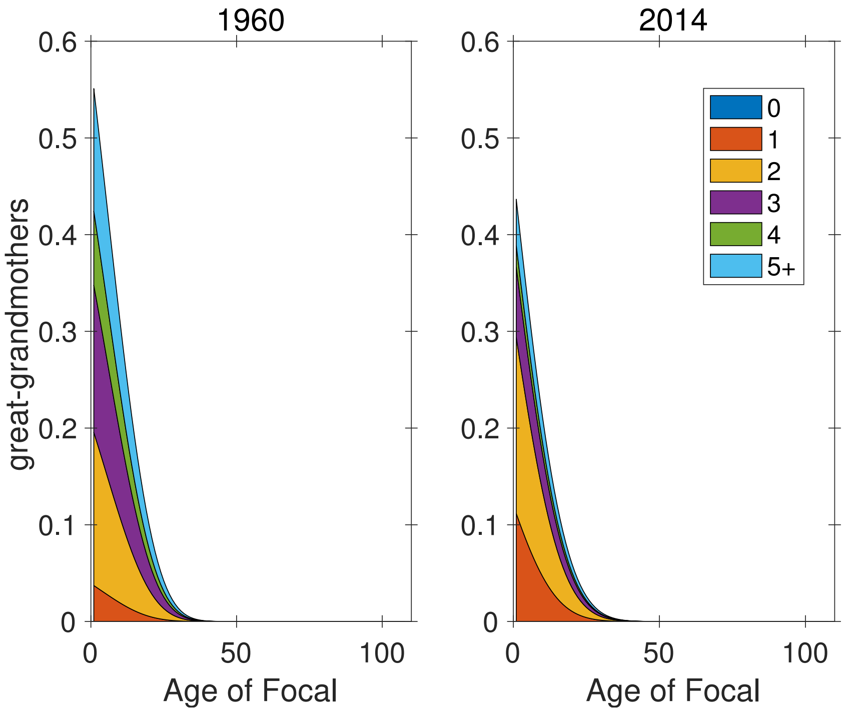


**Figure S-27:** Marginal parity structure of great-grandmothers as a function of the age of Focal.

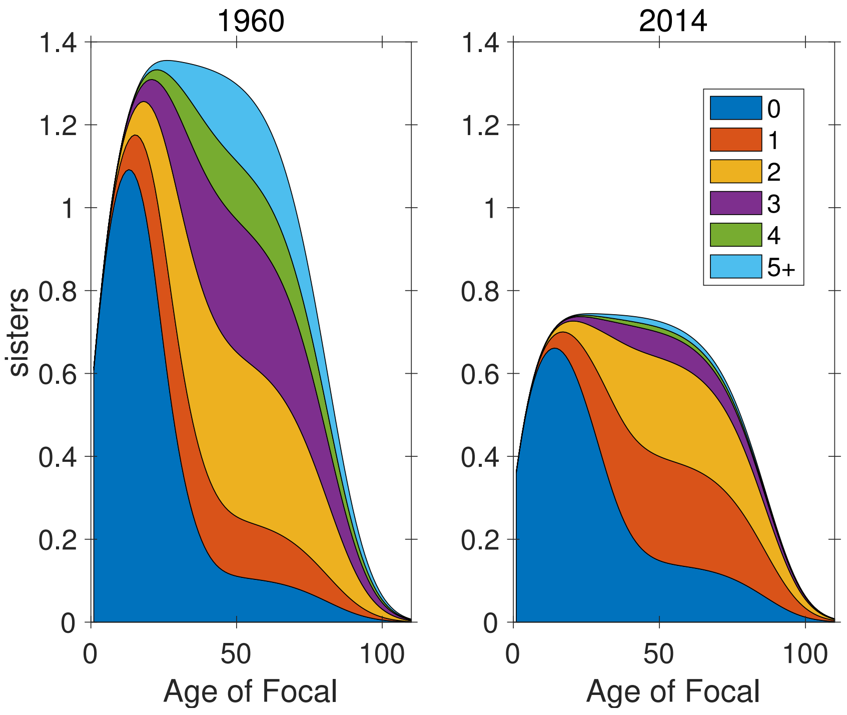


**Figure S-28:** Marginal parity structure of sisters as a function of the age of Focal

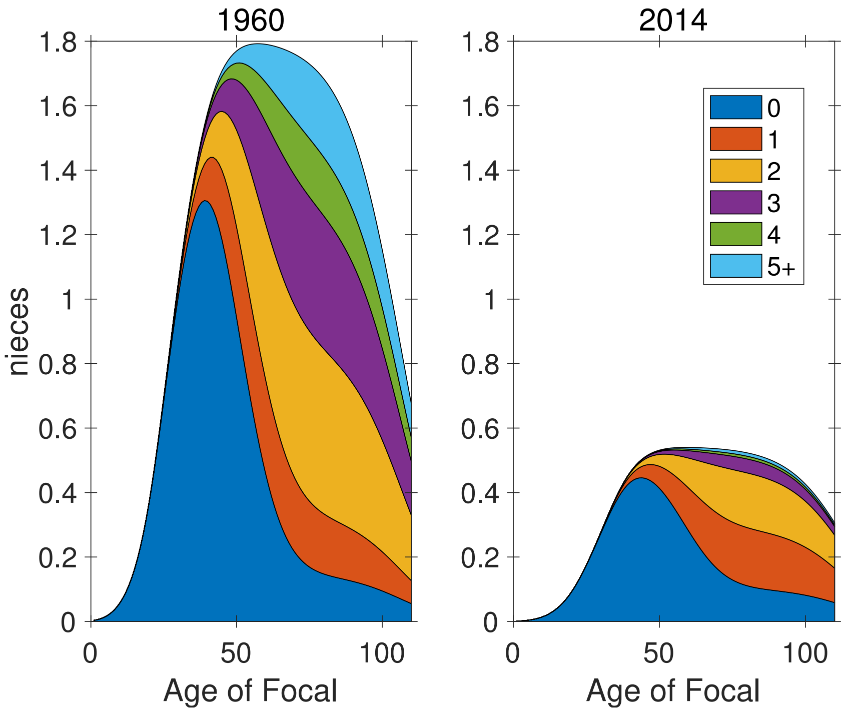


**Figure S-29:** Marginal parity structure of nieces as a function of the age of Focal.

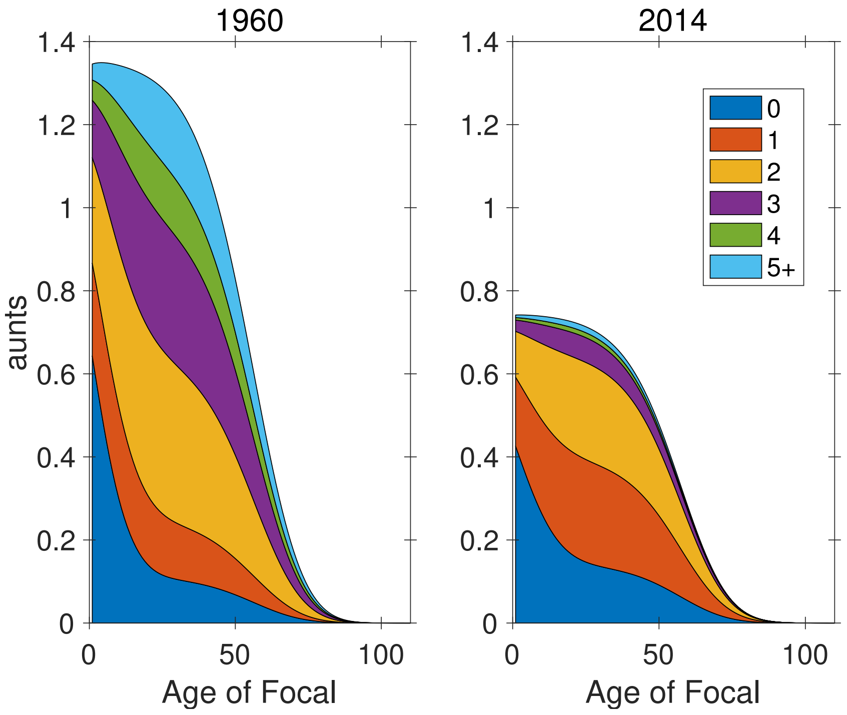


**Figure S-30:** Marginal parity structure of aunts as a function of the age of Focal.

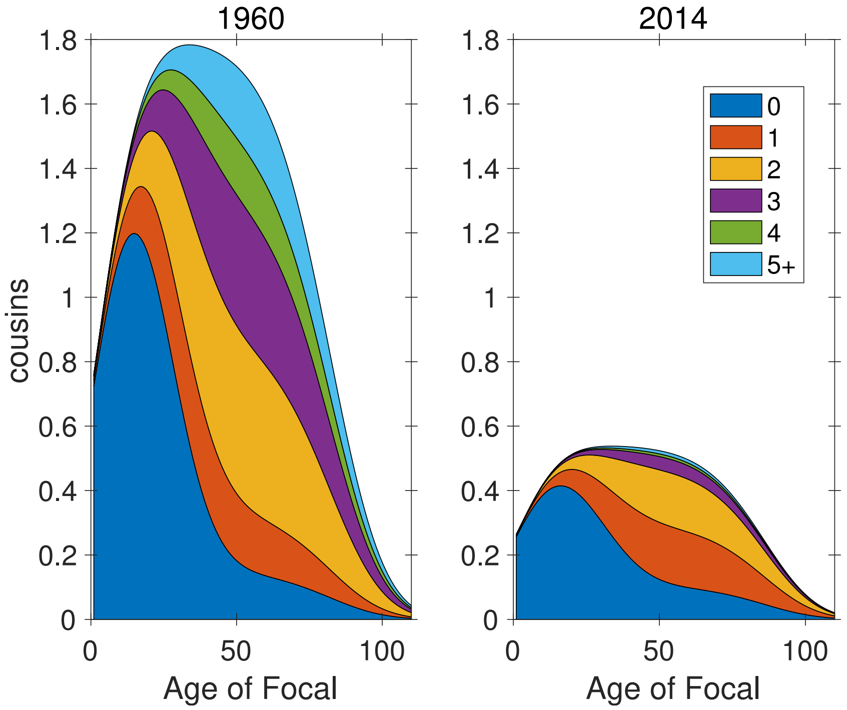


**Figure S-31:** Marginal parity structure of cousins as a function of the age of Focal.

#### 5 Marginal parity distributions by age of Focal

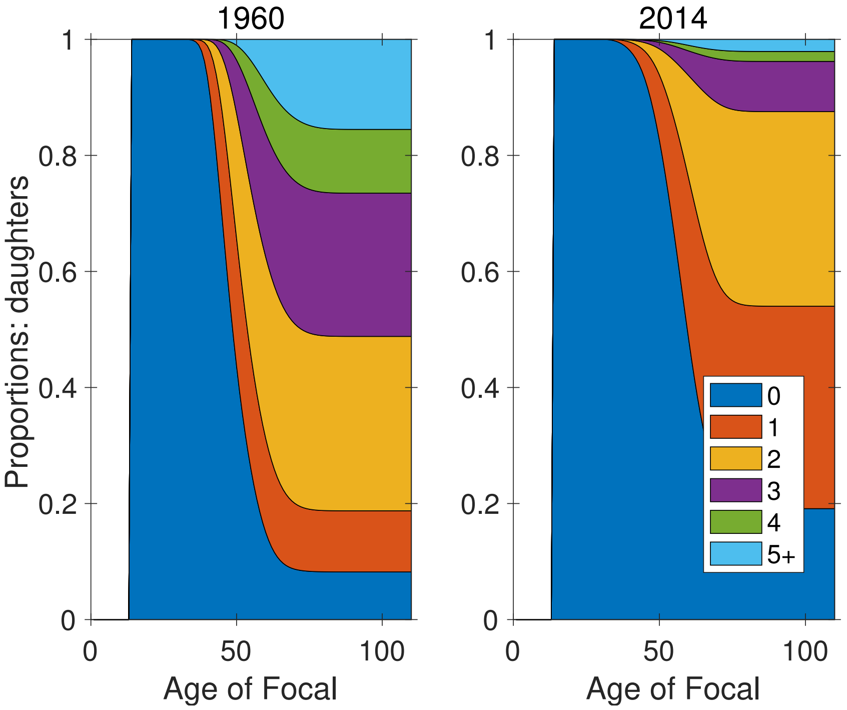


**Figure S-32:** Marginal parity distribution of daughters, as function of age of Focal

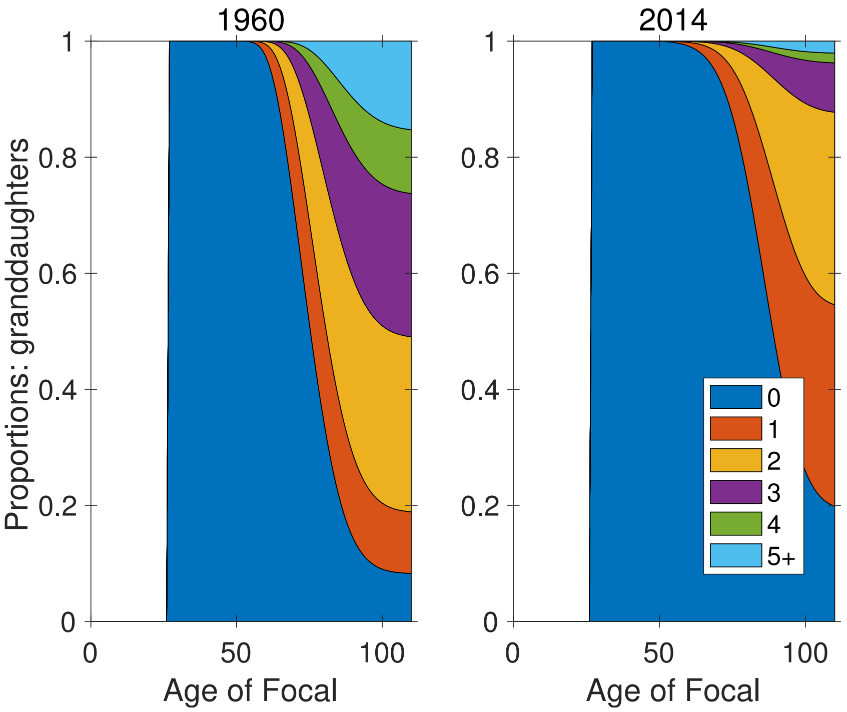


**Figure S-33:** Marginal parity distribution of granddaughters, as function of age of Focal

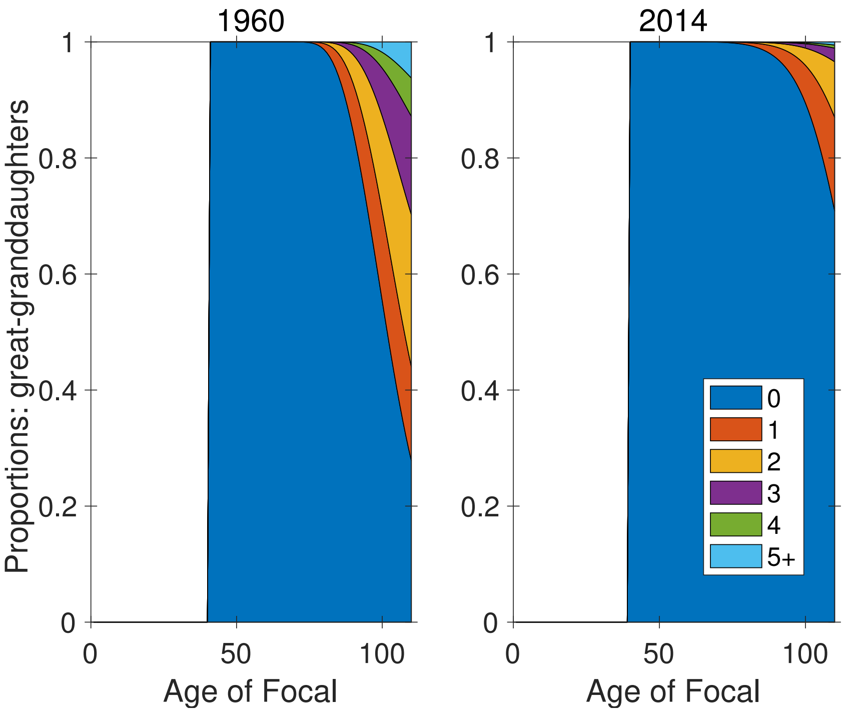


**Figure S-34:** Marginal parity distribution of great-granddaughters, as function of age of Focal

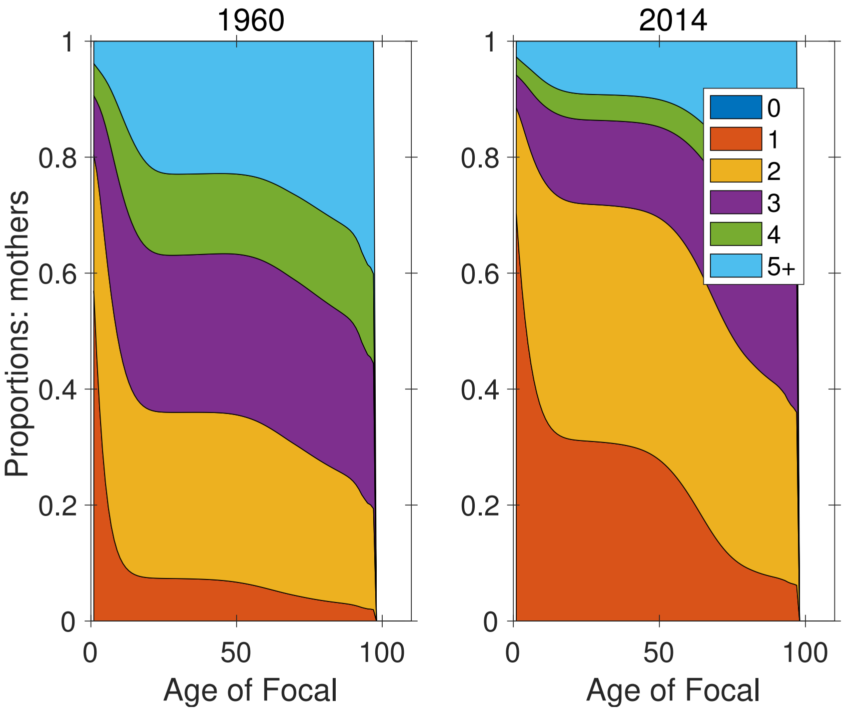


**Figure S-35:** Marginal parity distribution of mothers, as function of age of Focal

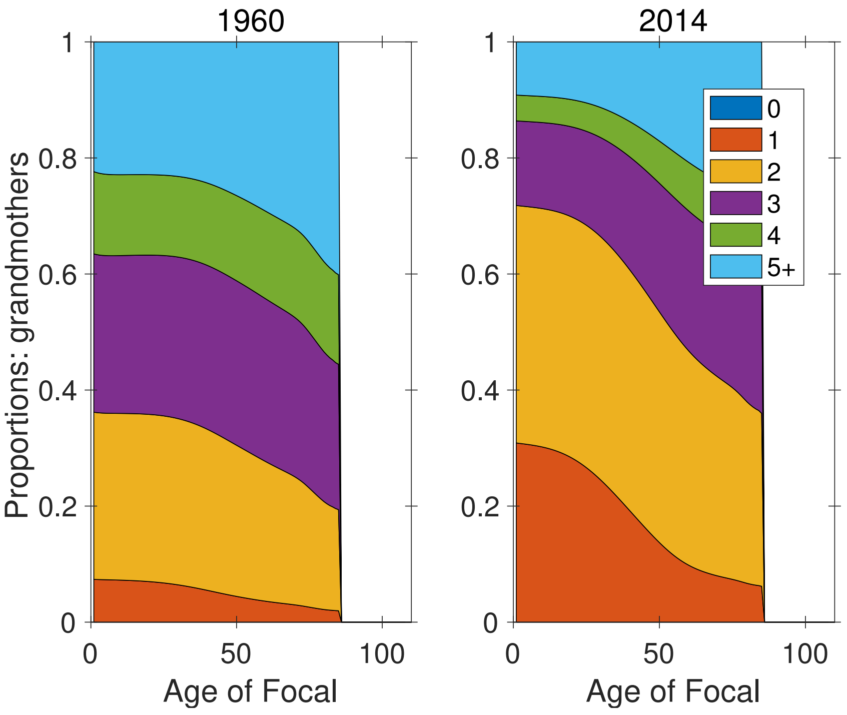


**Figure S-36:** Marginal parity distribution of grandmothers, as function of age of Focal

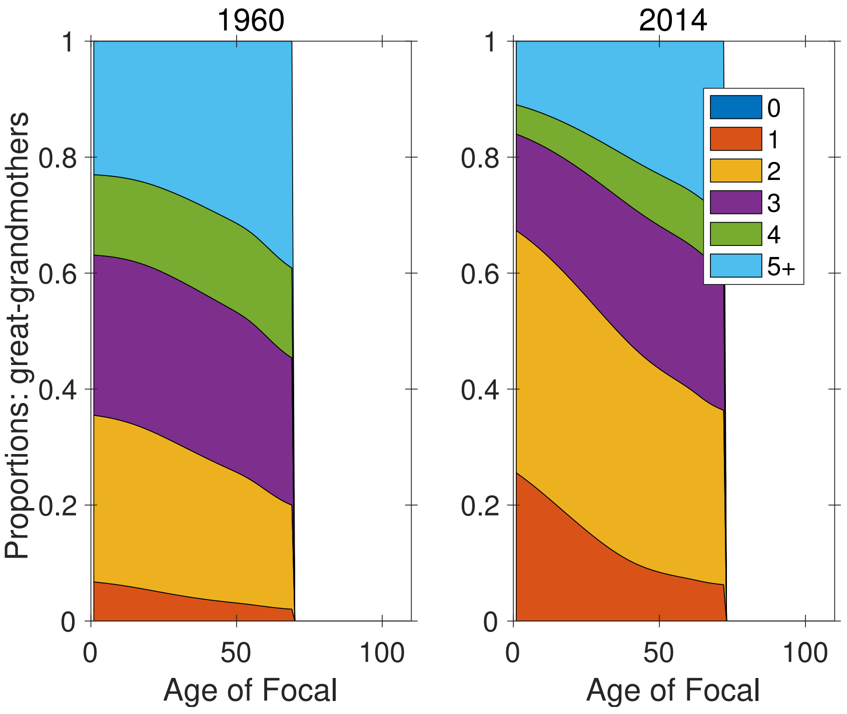


**Figure S-37:** Marginal parity distribution of great-grandmothers, as function of age of Focal

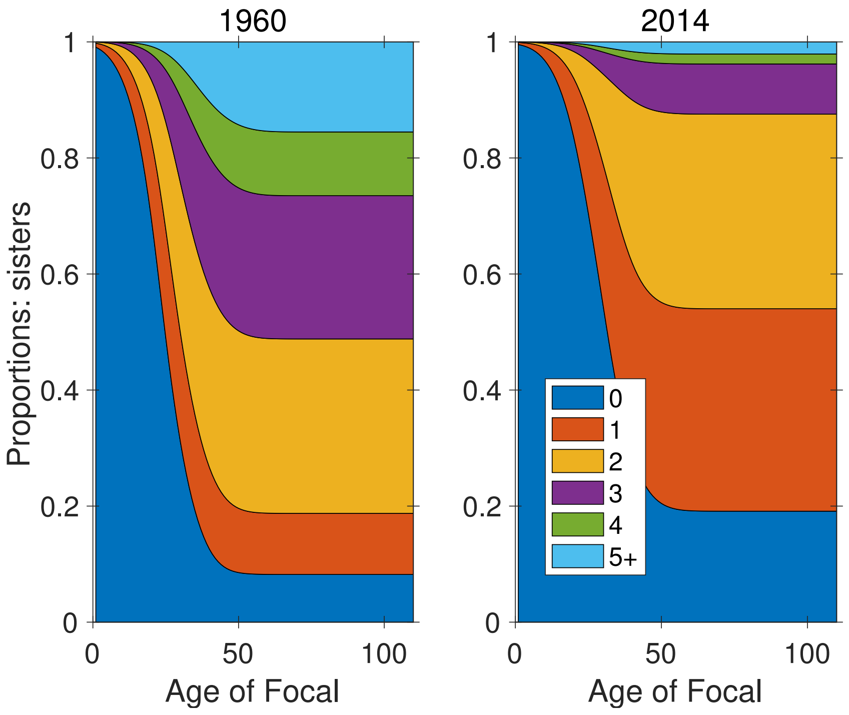


**Figure S-38:** Marginal parity distribution of sisters, as function of age of Focal

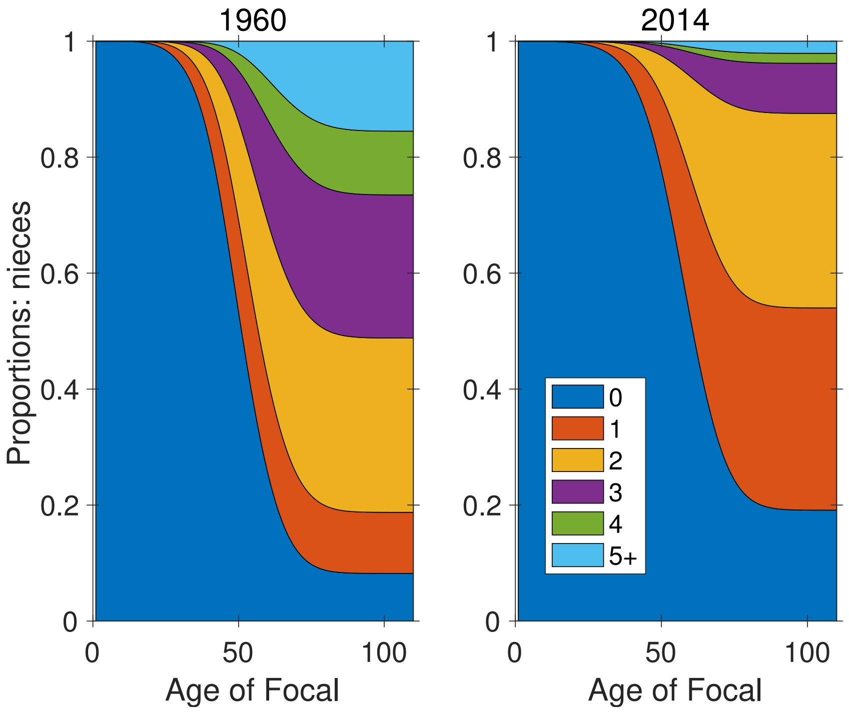


**Figure S-39:** Marginal parity distribution of nieces, as function of age of Focal

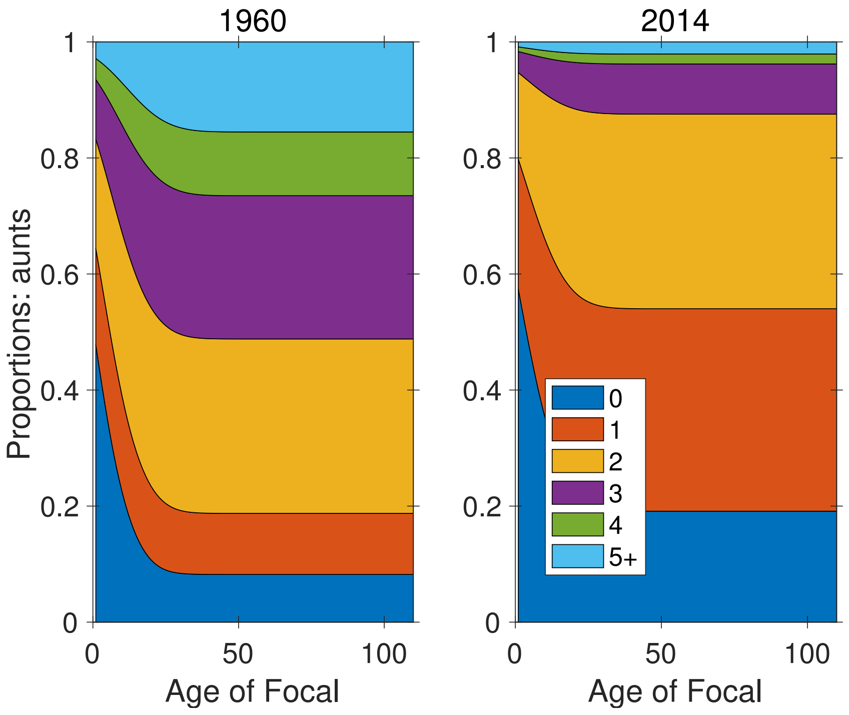


**Figure S-40:** Marginal parity distribution of aunts, as function of age of Focal

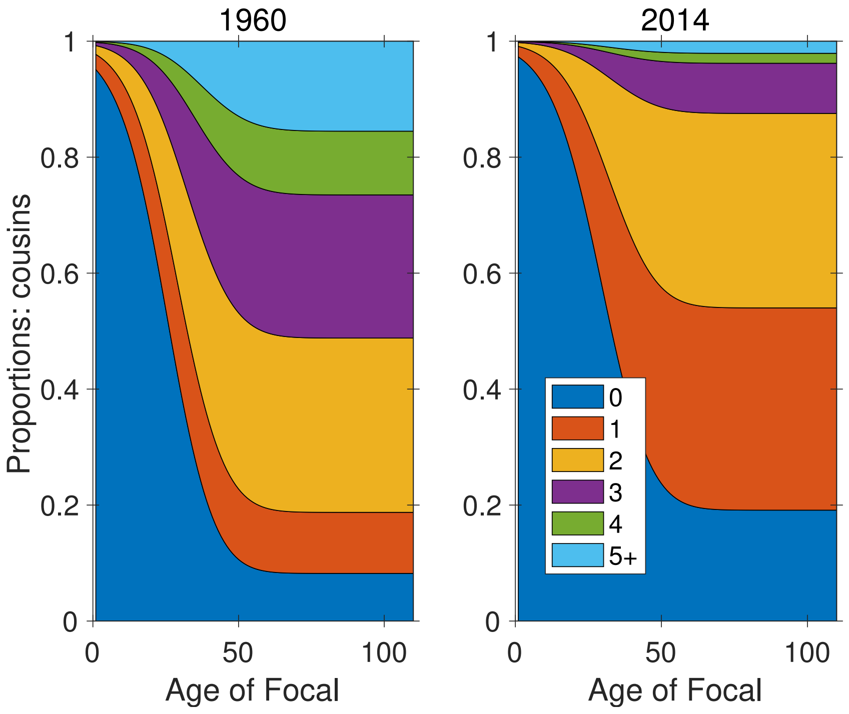


**Figure S-41:** Marginal parity distribution of cousins, as function of age of Focal

#### 6 Marginal parity distribution over time

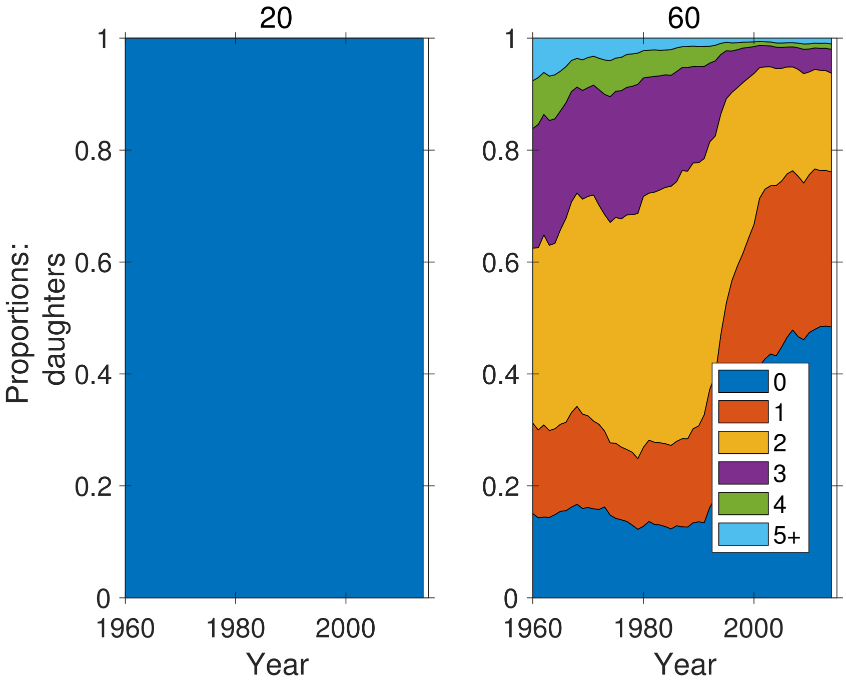


**Figure S-42:** Marginal parity distribution of daughters of Focal, as function of time, at ages 20 and 60 of Focal.

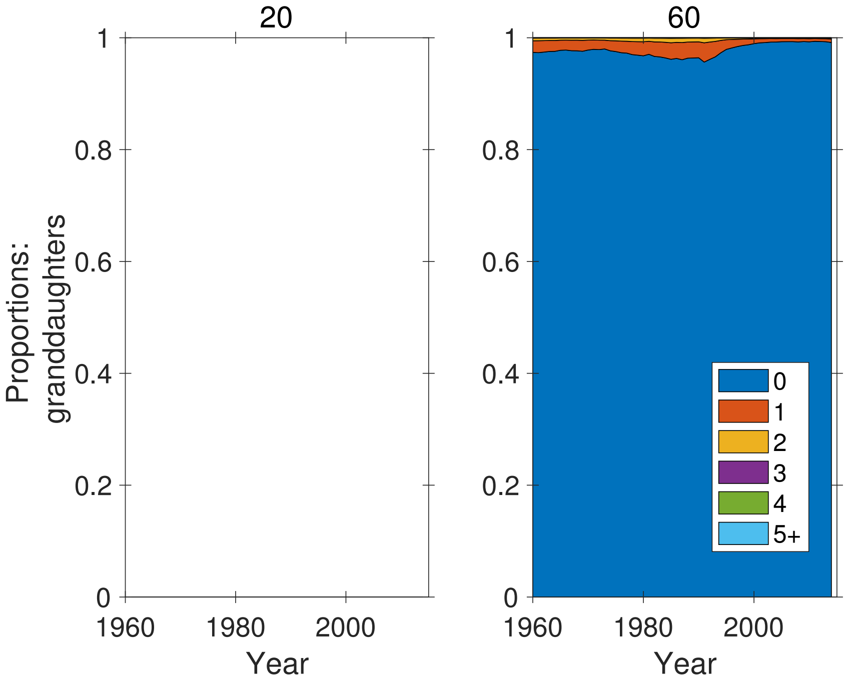


**Figure S-43:** Marginal parity distribution of granddaughters of Focal, as function of time, at ages 20 and 60 of Focal.

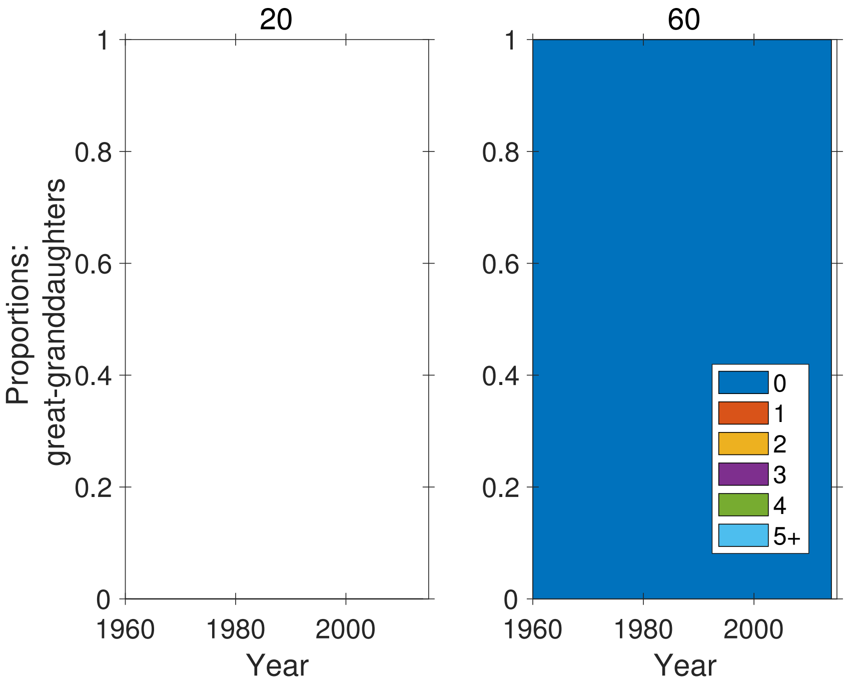


**Figure S-44:** Marginal parity distribution of great-granddaughters of Focal, as function of time, at ages 20 and 60 of Focal.

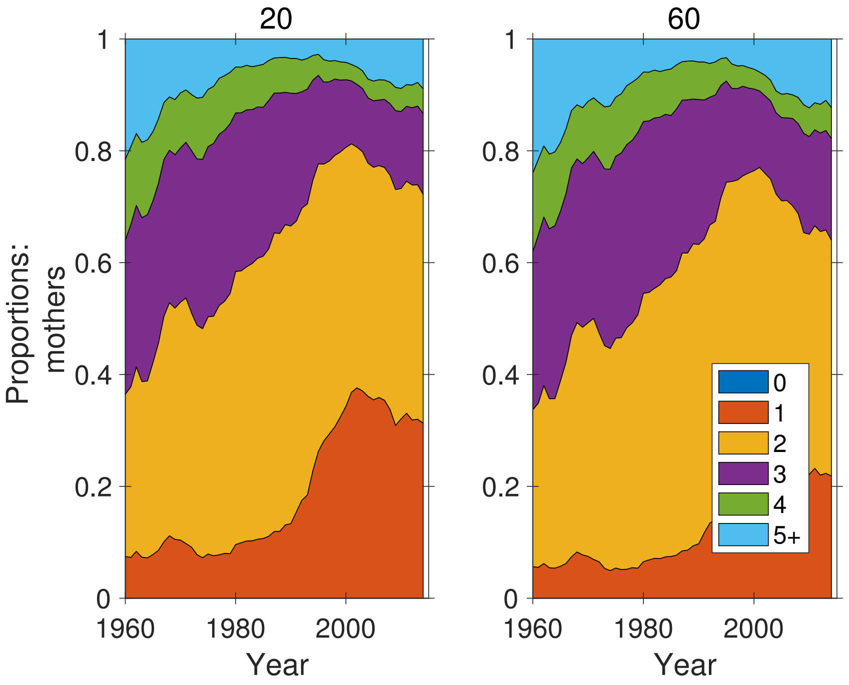


**Figure S-45:** Marginal parity distribution of mothers of Focal, as function of time, at ages 20 and 60 of Focal.

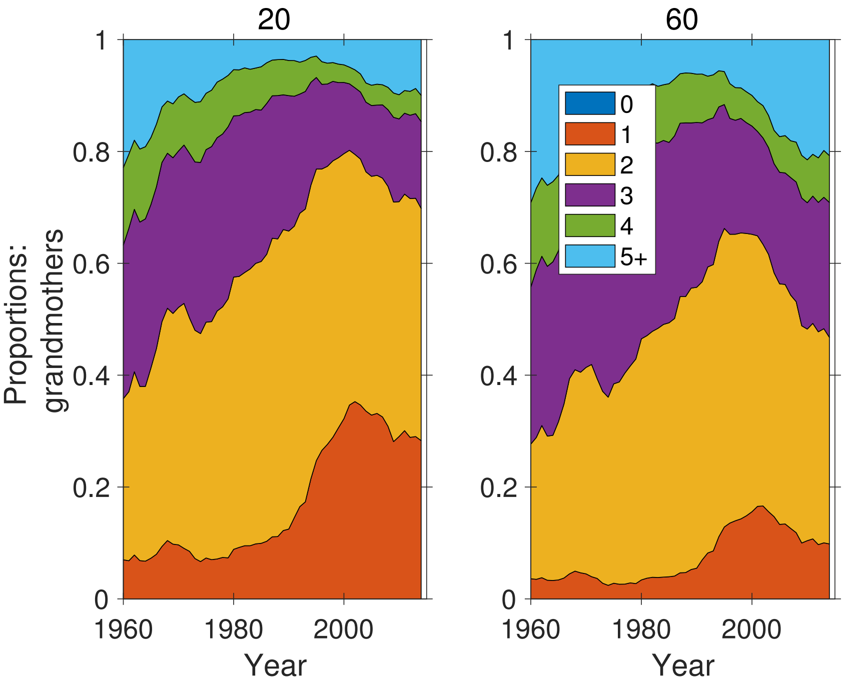


**Figure S-46:** Marginal parity distribution of grandmothers of Focal, as function of time, at ages 20 and 60 of Focal.

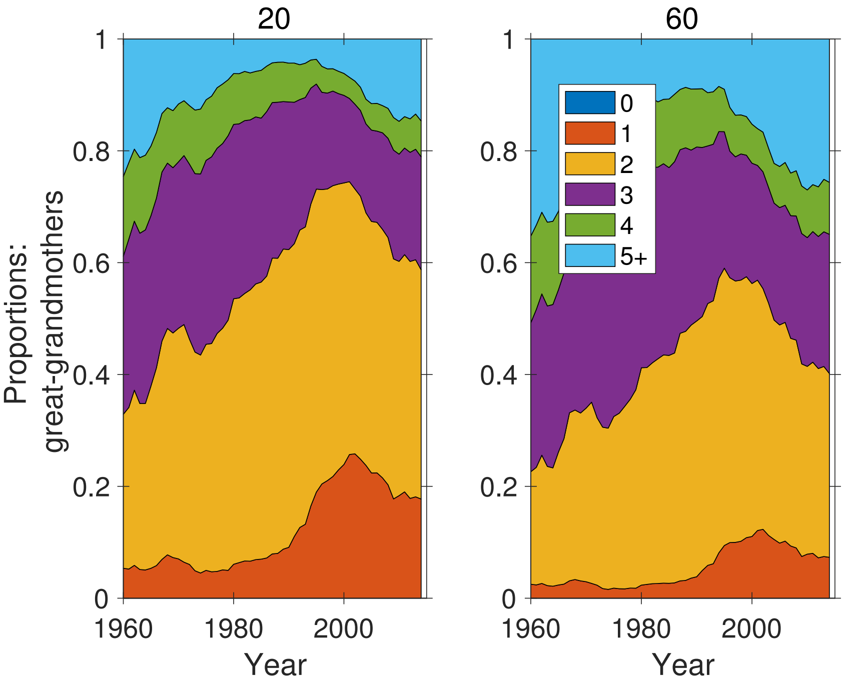


**Figure S-47:** Marginal parity distribution of great-grandmothers of Focal, as function of time, at ages 20 and 60 of Focal.

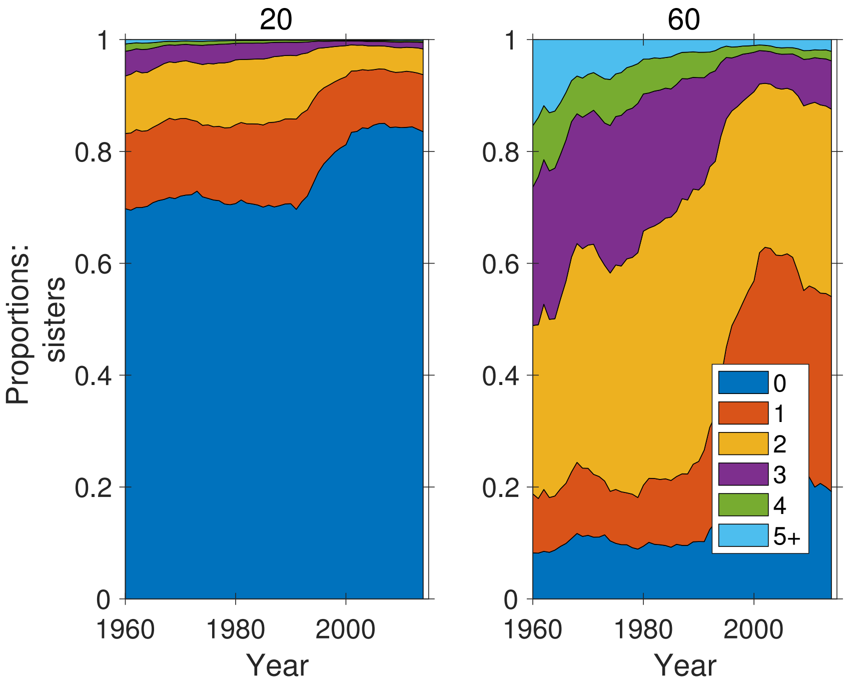


**Figure S-48:** Marginal parity distribution of sisters of Focal, as function of time, at ages 20 and 60 of Focal.

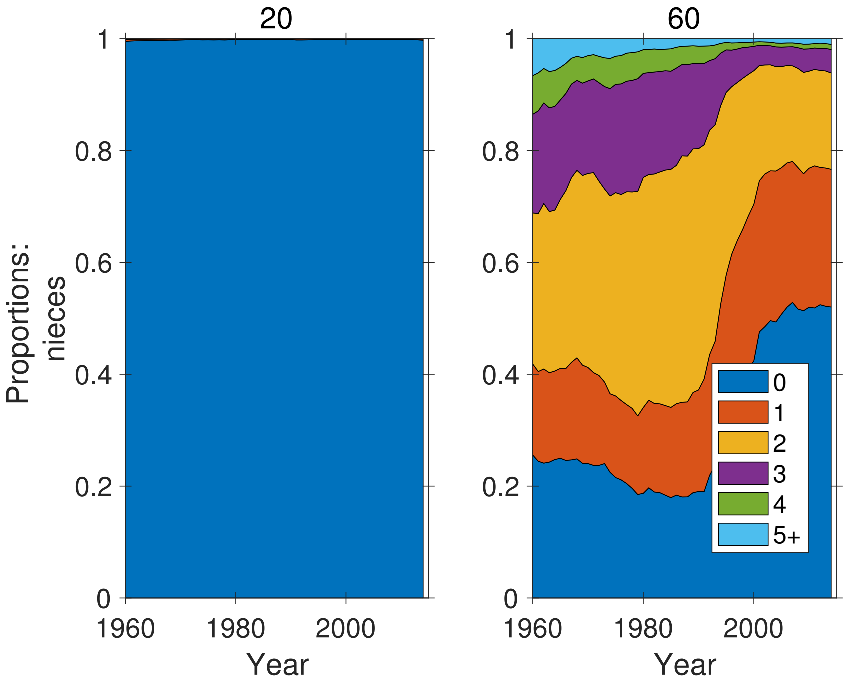


**Figure S-49:** Marginal parity distribution of nieces of Focal, as function of time, at ages 20 and 60 of Focal.

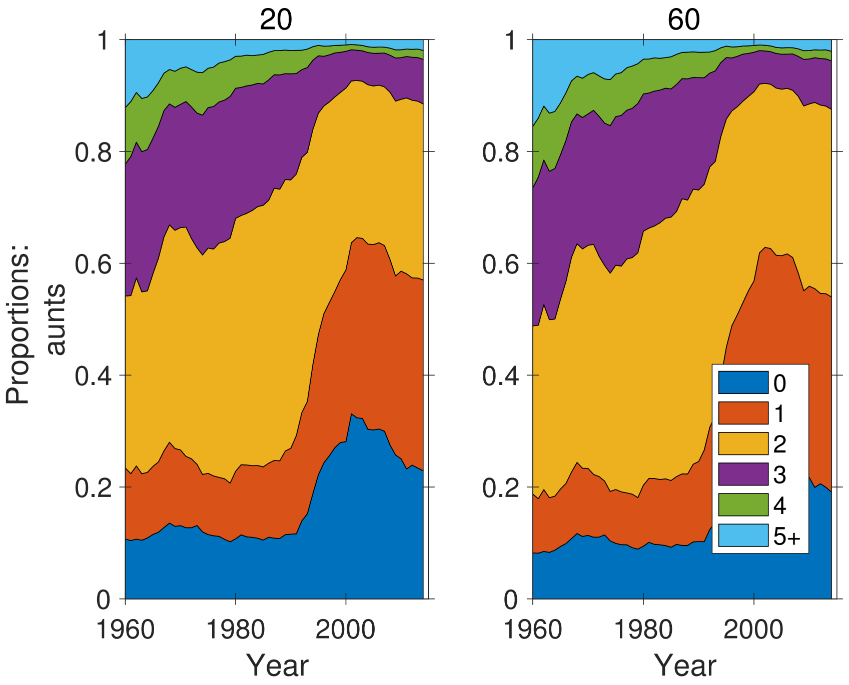


**Figure S-50:** Marginal parity distribution of aunts of Focal, as function of time, at ages 20 and 60 of Focal.

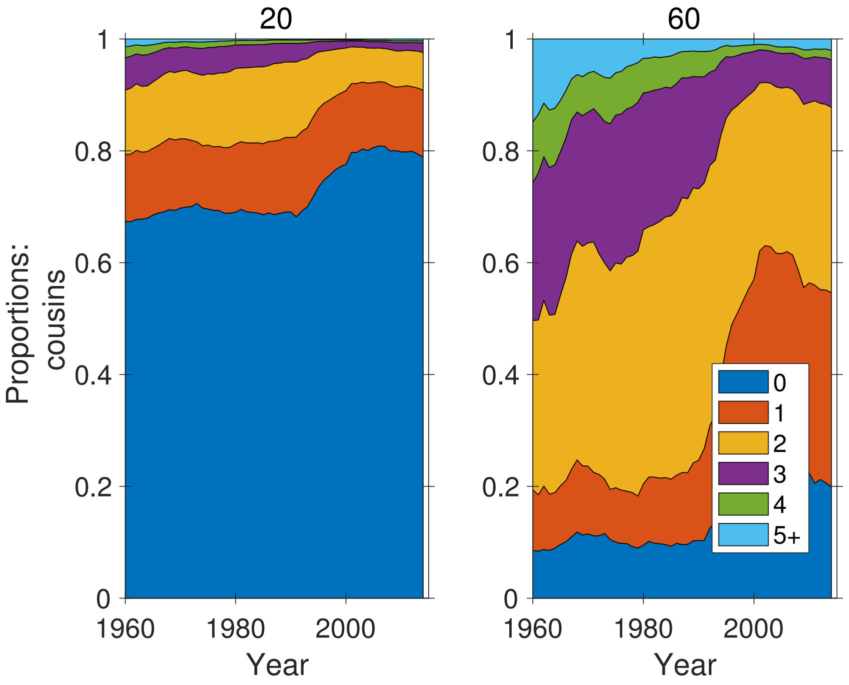


**Figure S-51:** Marginal parity distribution of cousins of Focal, as function of time, at ages 20 and 60 of Focal.

#### 7 Prevalence of low parity kin over time

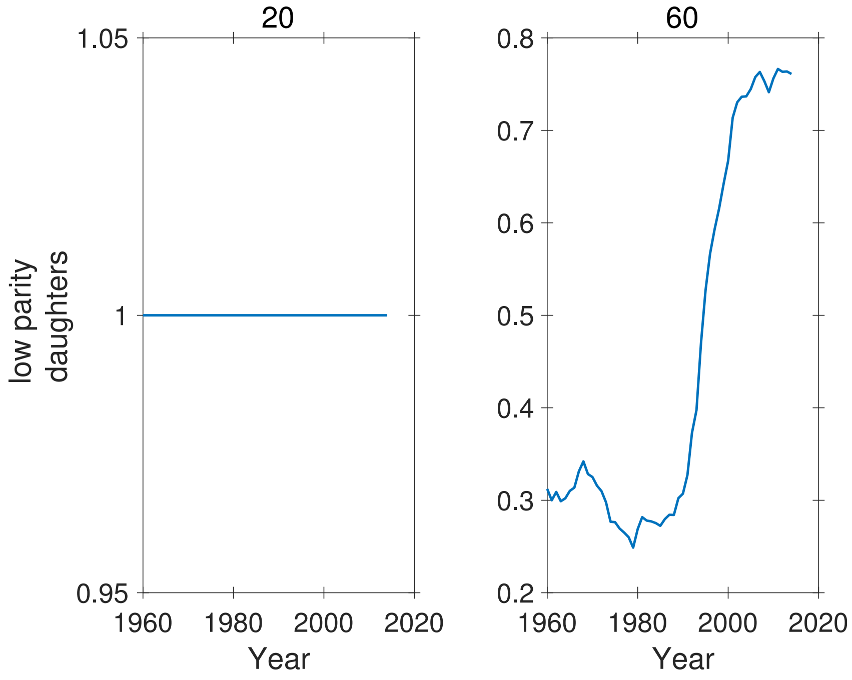


**Figure S-52:** Proportion of low parity (parities 0 and 1) daughters of Focal as a function of time, for ages 20 and 60 of Focal. Note different ordinate scales.

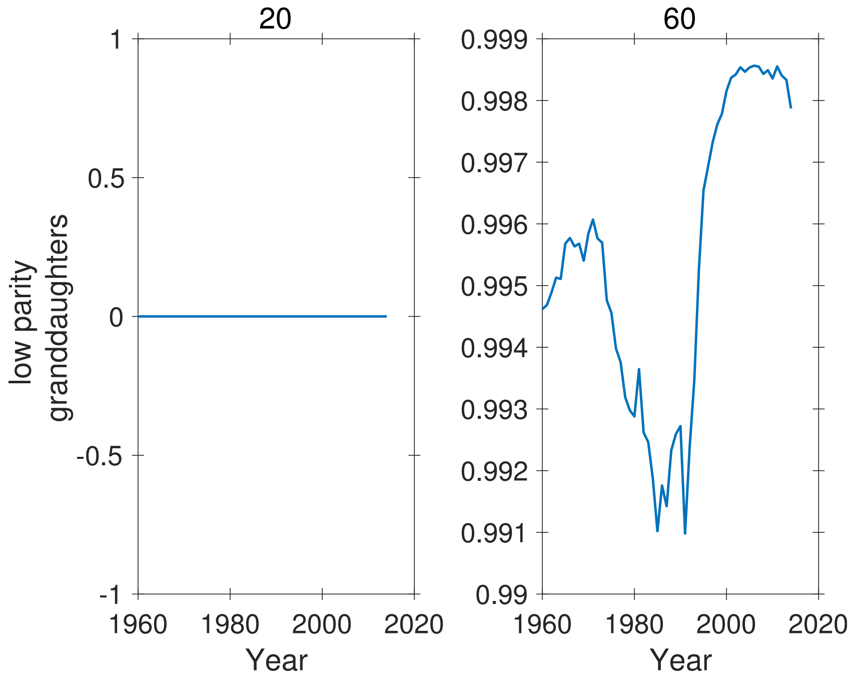


**Figure S-53:** Proportion of low parity (parities 0 and 1) granddaughters of Focal as a function of time, for ages 20 and 60 of Focal. Note different ordinate scales.

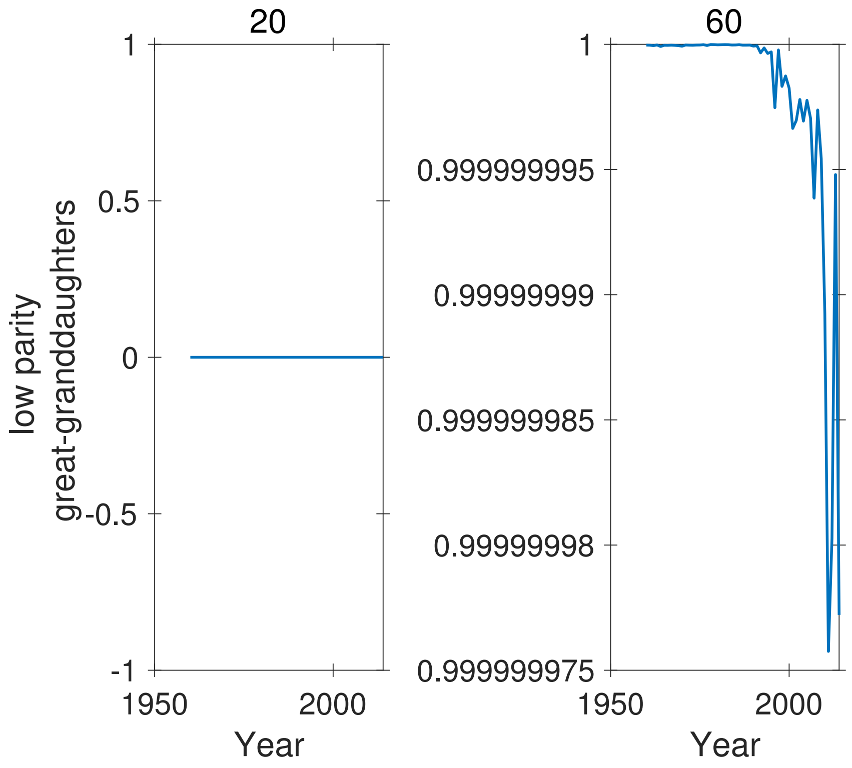


**Figure S-54:** Proportion of low parity (parities 0 and 1) great-granddaughters of Focal as a function of time, for ages 20 and 60 of Focal. Note different ordinate scales.

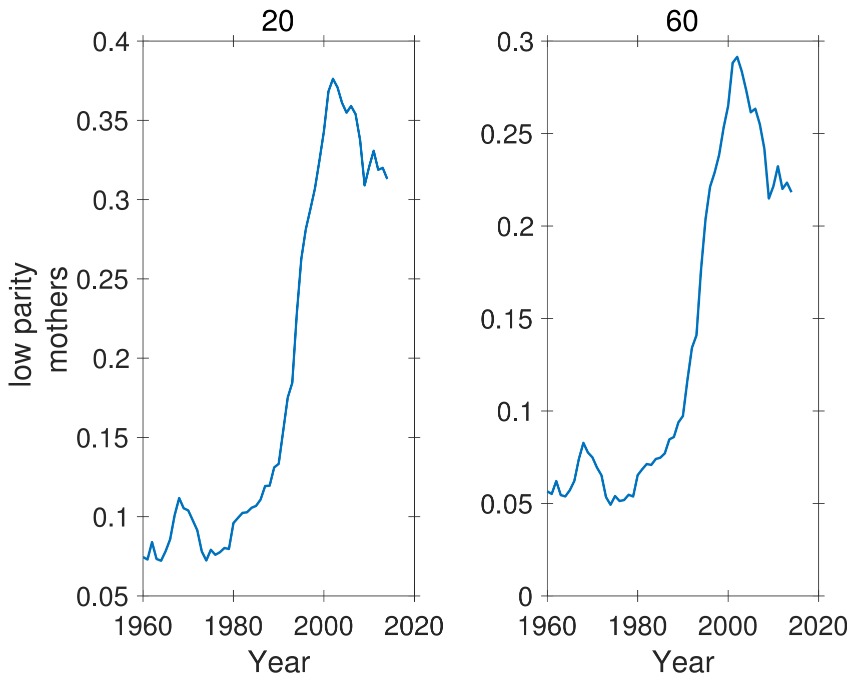


**Figure S-55:** Proportion of low parity (parities 0 and 1) mothers of Focal as a function of time, for ages 20 and 60 of Focal. Note different ordinate scales.

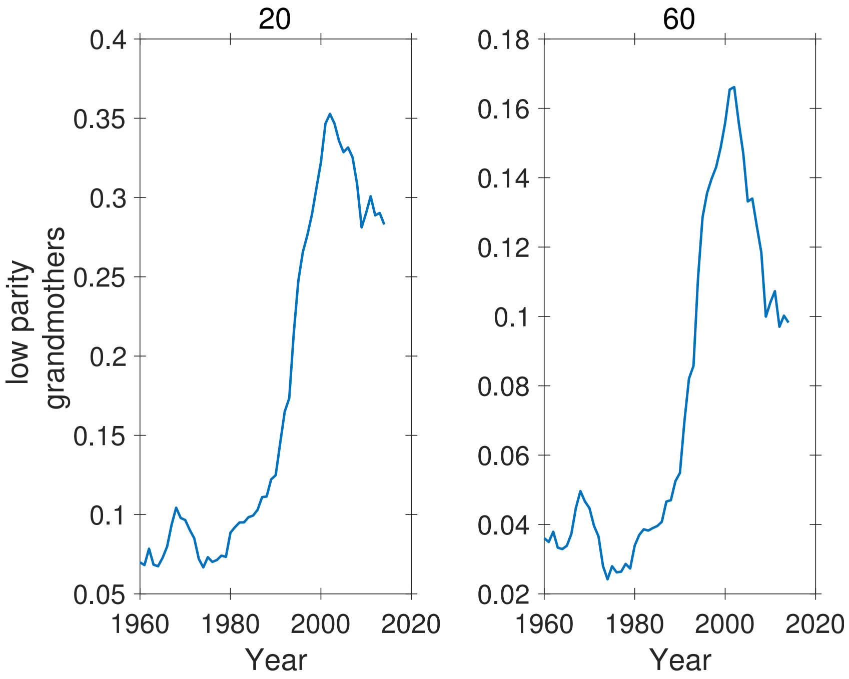


**Figure S-56:** Proportion of low parity (parities 0 and 1) grandmothers of Focal as a function of time, for ages 20 and 60 of Focal. Note different ordinate scales.

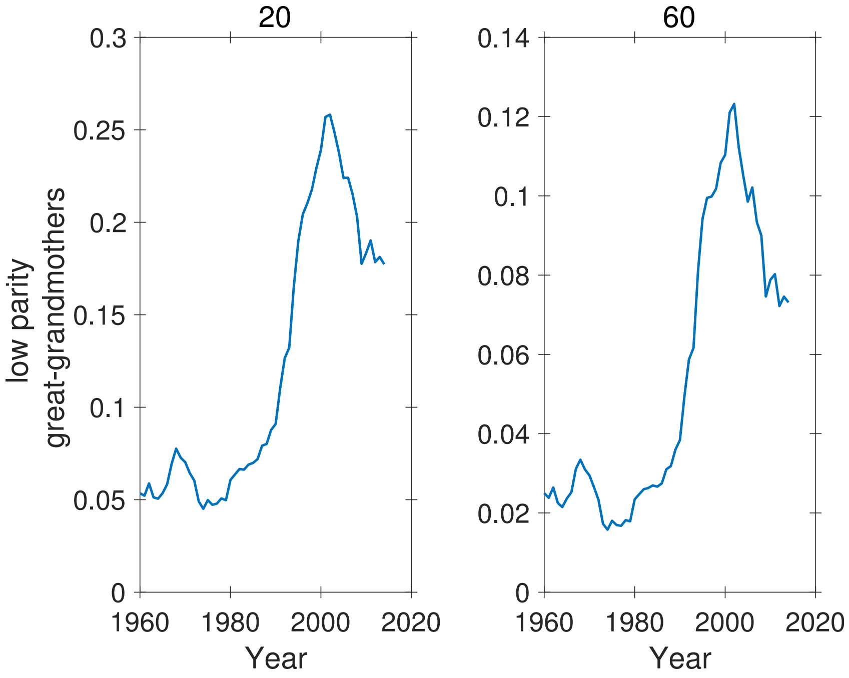


**Figure S-57:** Proportion of low parity (parities 0 and 1) great-grandmothers of Focal as a function of time, for ages 20 and 60 of Focal. Note different ordinate scales.

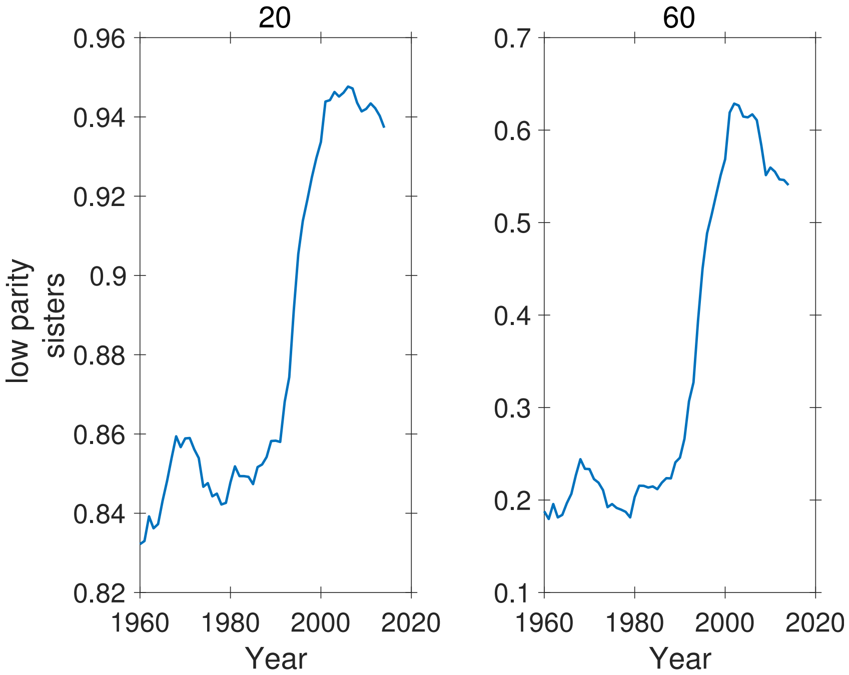


**Figure S-58:** Proportion of low parity (parities 0 and 1) sisters of Focal as a function of time, for ages 20 and 60 of Focal. Note different ordinate scales.

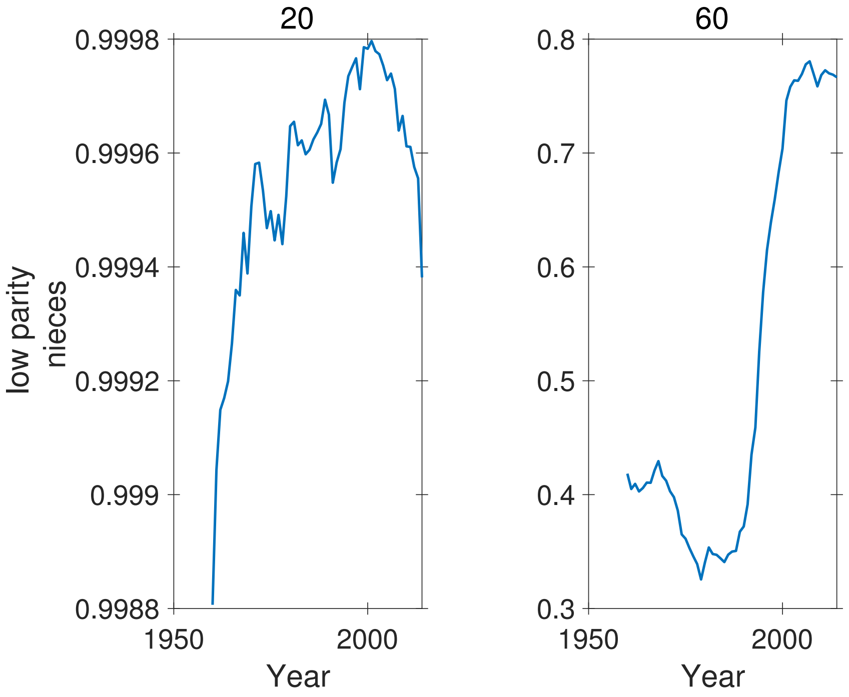


**Figure S-59:** Proportion of low parity (parities 0 and 1) nieces of Focal as a function of time, for ages 20 and 60 of Focal. Note different ordinate scales.

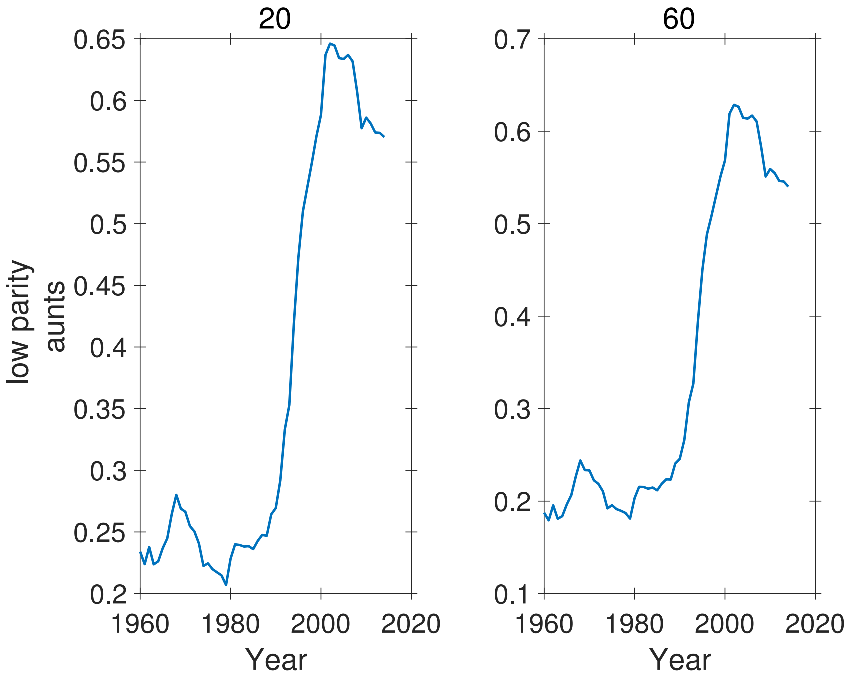


**Figure S-60:** Proportion of low parity (parities 0 and 1) aunts of Focal as a function of time, for ages 20 and 60 of Focal. Note different ordinate scales.

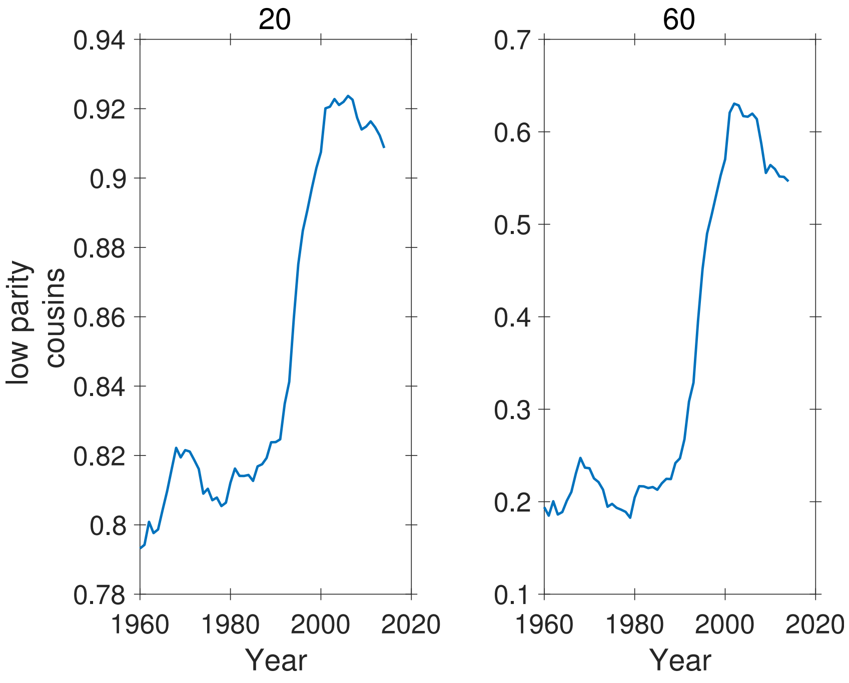


**Figure S-61:** Proportion of low parity (parities 0 and 1) cousins of Focal as a function of time, for ages 20 and 60 of Focal. Note different ordinate scales.
